# Visualizing proteins by expansion microscopy

**DOI:** 10.1101/2022.08.03.502284

**Authors:** Ali H. Shaib, Abed Alrahman Chouaib, Rajdeep Chowdhury, Daniel Mihaylov, Chi Zhang, Vanessa Imani, Svilen Veselinov Georgiev, Nikolaos Mougios, Mehar Monga, Sofiia Reshetniak, Tiago Mimoso, Han Chen, Parisa Fatehbasharzad, Dagmar Crzan, Kim-Ann Saal, Nadia Alawar, Janna Eilts, Jinyoung Kang, Luis Alvarez, Claudia Trenkwalder, Brit Mollenhauer, Tiago F. Outeiro, Sarah Köster, Julia Preobraschenski, Ute Becherer, Tobias Moser, Edward S. Boyden, A Radu Aricescu, Markus Sauer, Felipe Opazo, Silvio O. Rizzoli

## Abstract

Fluorescence imaging is one of the most versatile and widely-used tools in biology^1^. Although techniques to overcome the diffraction barrier were introduced more than two decades ago, and the nominal attainable resolution kept improving^2, 3^, fluorescence microscopy still fails to image the morphology of single proteins or small molecular complexes, either purified or in a cellular context^4, 5^. Here we report a solution to this problem, in the form of <u>o</u>ne-step <u>n</u>anoscale <u>e</u>xpansion (ONE) microscopy. We combined the 10-fold axial expansion of the specimen (1000-fold by volume) with a fluorescence fluctuation analysis^6, 7^ to enable the description of cultured cells, tissues, viral particles, molecular complexes and single proteins. At the cellular level, using immunostaining, our technology revealed detailed nanoscale arrangements of synaptic proteins, including a quasi-regular organisation of PSD95 clusters. At the single molecule level, upon main chain fluorescent labelling, we could visualise the shape of individual membrane and soluble proteins. Moreover, conformational changes undergone by the ∼17 kDa protein calmodulin upon Ca^2+^ binding were readily observable. We also imaged and classified molecular aggregates in cerebrospinal fluid samples from Parkinson’s Disease (PD) patients, which represents a promising new development towards improved PD diagnosis. ONE microscopy is compatible with conventional microscopes and can be performed with the software we provide here as a free, open-source package. This technology bridges the gap between high-resolution structural biology techniques and light microscopy, and provides a new avenue for discoveries in biology and medicine.

## Introduction

Optical microscopy has been one of the most valuable tools in biology for more than two centuries, and has been considerably enhanced by the introduction of super-resolution microscopy, two decades ago^2, 3^. Nevertheless, optical imaging remains difficult to perform below 10-20 nm^4, 5^. Several recent works have presented localization precisions down to 1-2 nm^8-10^, or even below^11^, but the application of such imaging resolution to biological samples has been severely limited by two fundamental problems. First, the achievable structural resolution is determined by the labeling density, which is limited by the size of the fluorescent probes (typically 1 nanometer or larger)^12^. Second, fluorophores can interact via energy transfer at distances below 10 nm, which results in accelerated photoswitching (blinking) and photobleaching, and thus in substantially lower localization probabilities^13^.

The solution to these two problems would be to separate the fluorophores spatially by the physical expansion of the specimen, in what is termed expansion microscopy (ExM^14^). To then reach the molecular scale, one would combine ExM with optics-based super-resolution. This has been attempted numerous times^15–22^, but the resulting performance typically reached only ∼10 nm. The ExM gels are dim because the fluorophores are diluted by the third power of the expansion factor, thus limiting optics techniques that prefer bright samples, as stimulated emission depletion (STED^23^), or saturated structured illumination (SIM^24^). In addition, the ExM gels need to be imaged in distilled water, since the ions in buffered solutions shield the charged moieties of the gels and diminish the expansion factor. The use of distilled water reduces the performance of techniques that rely on special buffers, as single molecule localization microscopy, SMLM^14, 25^ (**Extended Data Fig. 1**).

A third class of optical super-resolution approaches is based on determining the higher-order statistical analysis of temporal fluctuations measured in a movie, *e.g.* super-resolution optical fluctuation imaging (SOFI^26^) or super-resolution radial fluctuations (SRRF^6, 7^). The resolution of these approaches is inversely correlated to the distance between the fluorophores^6, 7, 27^ and they do not require especially bright samples or special buffers, implying that they should benefit from ExM. To test this hypothesis, we combined X10 expansion microscopy^28, 29^ with SRRF^6, 7, 27^ and established a technique we term <u>o</u>ne-step <u>n</u>anoscale <u>e</u>xpansion (ONE) microscopy (**Extended Data Fig. 1**). ONE was implemented using conventional confocal or epifluorescence microscopes and reached an imaging performance that enables imaging individual protein shapes. To aid in its implementation, we generated a ONE software platform, as a plug-in for the popular freeware ImageJ (Fiji) (**Supplementary Fig. 1**; Supplementary Software). A shortcoming of this technique, in comparison to more established procedures as *d*STORM-ExM, is that its axial resolution is equivalent to the resolution of the confocal microscope, divided by the expansion factor, and is thus substantially poorer than the XY-plane resolution.

### Principles and validation of ONE microscopy

We first attached a gel-compatible anchor (Acryloyl-X) to protein molecules, either purified or in a cellular context, and then embedded these samples into a swellable X10 gel^28, 29^. Proteins were hydrolysed by proteinase K or by heating in alkaline buffers, leading to main chain breaks. This enables a highly-isotropic 10-fold expansion of the sample, which is achieved by distilled water incubations^28, 29^. The alkaline peptide hydrolysis we use, at >100°C, has been initially designed to generate free amino acids from protein mixtures (*e.g*. urine), for clinical investigations. It is therefore sufficient to cause the fragmenting of every protein, under our analysis conditions. Its current implementation, according to the protocol described in Methods, provides a mild fragmentation of individual proteins, while maintaining most of the fluorophores that were present on proteins before expansion (for more details see the Supplementary Discussion).

We then imaged the samples using wide-field epifluorescence or confocal microscopy, acquiring series of images (movies) of hundreds to thousands of images (ideally 1500-2000) in which the fluorescence intensity of the fluorophores fluctuates (**Extended Data Fig. 2**). Each pixel of a frame was then magnified into a large number of subpixels, and the local radial symmetries of the frame (which are due to the radial symmetry of the microscope’s point-spread-function, PSF) were measured. This parameter, termed “radiality” was analyzed throughout the image stack, by higher-order temporal statistics, to provide the final, fully resolved image^6, 7, 27^.

In theory, the precision of the SRRF technique should reach values close to 10 nm6. SRRF should therefore be able to separate fluorophores found at 20 nm from each other, provided the signal-to-noise ratio is sufficiently high. We found this to be the case, using nanorulers (provided by GATTAquant^30^), of precisely defined size (**Supplementary Fig. 2**).

In practice, most previous implementations of SRRF have reached ∼50-70 nm. This is partly due to the fact that the presence of overlapping fluorophores reduces radiality in conventional samples^6, 7^, and partly due to the aims of the respective SRRF implementations, which did not target ultimate performance in terms of resolution, and therefore did not optimize a number of parameters. First, the highest resolutions are obtained by analysing higher-order statistical correlations, whose precision is dependent on the number of frames acquired, as discussed not only for SRRF, but for SOFI as well^26^. While most publications use less than 300 frames, we found that results are optimal when using 1500-2000 frames (**Supplementary Fig. 3**). Working with low frame numbers reduces the achievable resolution, even when working with ExM gels^20, 31^. Second, the signal-to-noise ratio needs to be optimized carefully^32^, as we also demonstrate in **Supplementary Fig. 4**.

These limitations are alleviated by ExM (see Supplementary Discussion for more details). The distance between the fluorophores increases, enabling the study of intensity fluctuations from individual dye molecules independently. The signal-to-noise ratio also increased, even for idealized samples consisting only of fluorescently-conjugated nanobodies in solution (**Extended Data Fig. 3**). This approach should therefore allow an optimal SRRF performance, which, divided by the expansion factor, should bring the resulting imaging precision to the molecular scale (**Extended Data Fig. 1b**), as long as the gel expands isotropically in all dimensions. The X10 gel, based on *N,N-*dimethylacrylamide acid (DMAA), rather than the acrylamide used in typical ExM protocols, has a more homogeneous distribution of cross-links^33^, thus leading to fewer errors in expansion (see^34^ for a further discussion on gel homogeneity).

To assess the performance of ONE microscopy in a cellular context, we first analysed microtubules, a standard reference structure in super-resolution imaging techniques^17^. Gels were stabilized in specially-designed imaging chambers (**Supplementary Fig. 5**), which enabled us to image the antibody-decorated microtubules at both 10-fold and ∼3.5-fold expansion (the ZOOM ExM technique^35^ was used for the latter; **Fig. 1**).

The microtubule sizes matched previous measurements^36, 37^, ∼60 nm in diameter, when labelled with secondary antibodies, and around 30-35 nm, when labelled with secondary nanobodies (**Extended Data Fig. 4c,d**).

We then evaluated a purified ALFA-tagged EGFP construct bound simultaneously by two anti-GFP nanobodies^38^ and by an anti-ALFA nanobody^39^. This results in a triangular semi-flexible arrangement, which we termed a “triangulate smart ruler” (TSR, **Fig. 1b; Extended Data Fig. 5**). The TSR aspect observed in ONE microscopy is consistent with crystal structures of nanobody-EGFP and nanobody-ALFA complexes (**Fig. 1b,c**).

**Figure 1.**
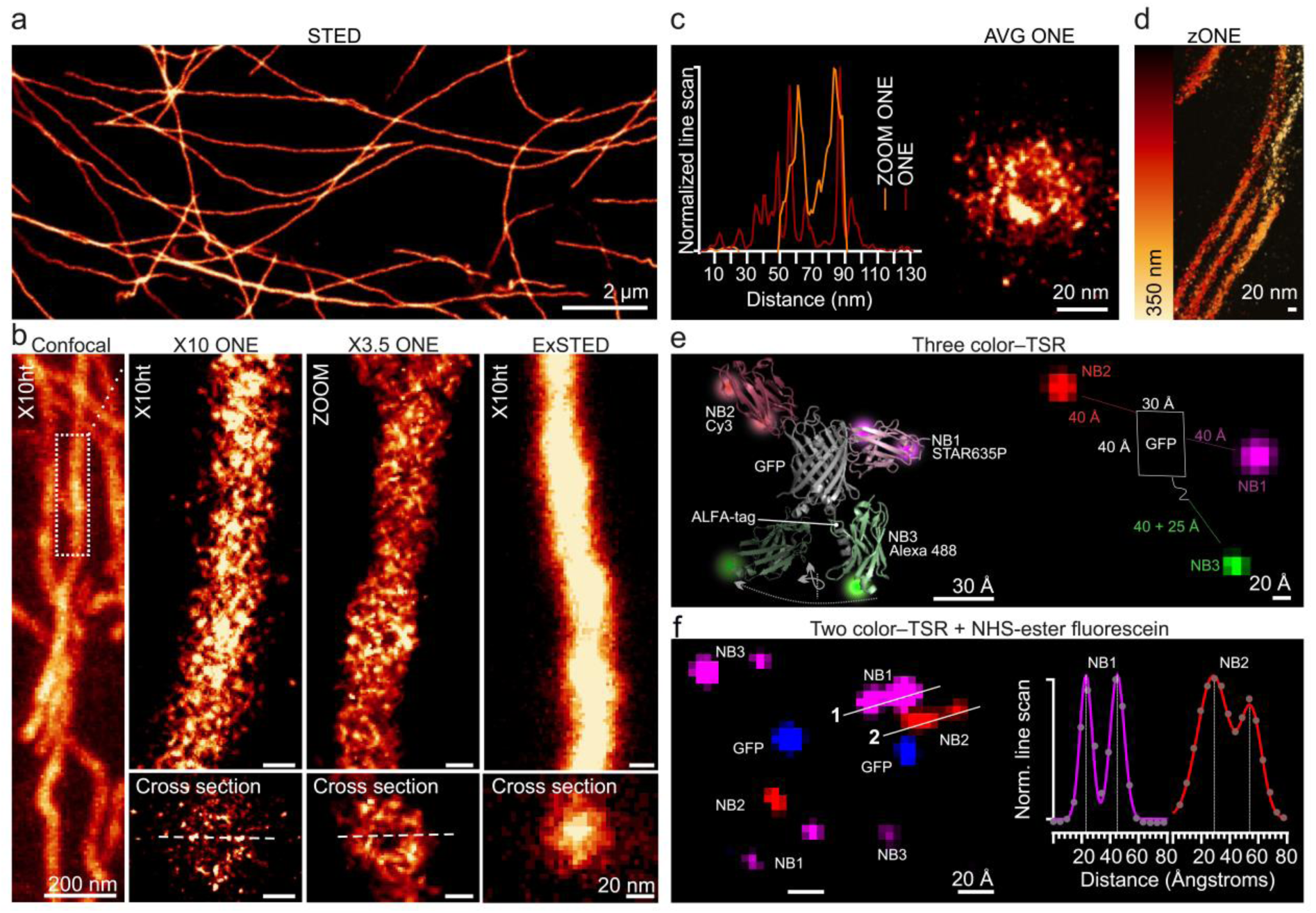
ONE performance in cellular samples and *in vitro*. **a-b**, Tubulin immunostainings (relying on primary and secondary antibodies) imaged using STED, without expansion (**a**), confocal after X10 expansion, ONE microscopy, ONE with ZOOM ExM (3.5-fold), and STED with X10 expansion (ExSTED, **b**). **c**, The graph depicts the line scans indicated by the dashed lines, which come from two different gels, with different expansion factors (10-fold for ONE, 3.5-fold for ZOOM-ONE). The image (AVG ONE) shows an average of 36 cross-sections. **d**, ONE microscopy images at different Z-axis levels, obtained by confocal scanning at different heights (zONE). **e**, The general scheme of the GFP-based assemblies (generated in Pymol, using the PDB structures 6I2G and 3K1K), along with a typical ONE microscopy image, with the rough positioning of the molecules indicated by the cartoon. The three nanobodies carry three spectrally different fluorophores. **f**, Two further examples, relying on a design in which NB1 and NB3 carry identical fluorophores. To detect GFP, the samples were labelled with NHS-ester fluorescein, after homogenization. Here we used a small pixel size, enabling the detection of two fluorophores connected to the nanobodies (see Supplementary Figure 6). Line scans across the indicated fluorophores are also shown.

To reveal the protein molecules themselves (the 10-fold expansion eliminates the endogenous GFP fluorescence for example (**Fig. 1e**)), we labelled the TSRs using NHS-ester fluorescein^40, 41^, which is sufficiently stable, under our imaging conditions, for this type of experiment (**Supplementary Fig. 7**). This is possible because proteins are broken during homogenization at multiple main chain positions, and each resulting peptide has an exposed amino terminal group that can be efficiently conjugated with N-hydroxysuccinimide ester (NHS-ester) functionalized fluorophores. It is known that nanobodies are not as strongly anchored to ExM gels as other proteins, owing to their low lysine content (only 2 lysines for the ALFA nanobody), and most of their peptides are lost after homogenization^17^. Their fluorescein signal is therefore poorer than that of GFP (**Fig. 1f**, **Extended Data Fig. 5g**). This is an unexpected bonus of nanobodies, because nanobody signals do not obscure those resulting from the protein of interest (see Supplementary Fig. 6 for a gallery of examples). In these experiments we used a smaller pixel size than in Fig. 1f (0.48 nm vs. 0.98 nm), which enabled us to often observe dual fluorophores, in agreement with the fact that these nanobodies can be labelled at two positions. The distances between the two fluorophores on one nanobody are consistent with the size of nanobody molecules (see the graph in **Fig. 1f**). Measuring the full width at half-maximum (FWHM) of the fluorescence signals resulted in an apparent particle size of ∼1 nm in the different fluorescence signals, including the fluorescein channel (**Extended Data Fig. 5**). Values within the same range are obtained when turning to an often-used technique to quantify the resolution of individual images in super-resolution fluorescence imaging, as well as in conventional electron microscopy, the Fourier Ring Correlation (FRC) determination^42, 43^. We applied this approach to our images, relying on the NanoJ-SQUIRREL package^42^ which has a blockwise implementation, to provide FRC values for different regions within individual images (**Extended Data Fig. 5**), obtaining values within the low single-digit nanometer range.

### ONE microscopy can reveal protein shapes

Considering the fact that proteins expand 1000x in volume but fluorophores do not, we hypothesized that our NHS-ester labelling method could be optimized to enable the analysis of protein shapes by ONE microscopy. In the TSR experiments (Fig. 1) we used limited NHS-ester labelling, to avoid the fluorescent labelling of the nanobodies, which also limited the GFP labelling to poorly distinguishable blobs. To observe the GFP shape better, we used optimal labelling conditions (high excess of NHS-ester fluorescein), and then proceeded to ONE microscopy. The expected shape and size were obtained for the GFP molecules (**Supplementary Fig. 8**). We next applied this approach to antibody molecules, and we could observe immediately recognizable outlines for IgGs, IgAs and IgMs (**Fig. 2a-c, Supplementary Fig. 9**). Fluorescent labels attached to secondary IgG antibodies could also be observed in the same images (**Fig. 2a; Supplementary Fig. 9**) and also in complexes between fluorescently-conjugated primary and secondary antibodies, or nanobodies (**Fig. 2d**).

We then applied the same labelling method to a membrane protein, the full-length β3 human γ-aminobutyric acid (GABA_A_) receptor homopentamer, a ligand-gated chloride channel^44^, producing imaging that resembled “front” and “side” views of the receptor, similar to its structure, as derived from crystallography and single-particle cryogenic electron microscopy (Cryo-EM) structures (**Fig. 2e,f**; **Supplementary Fig. 10**). It is worth noting that particles observed by ONE microscopy are indeed single molecules, and no averaging or classification has been performed on these datasets.

We next investigated a protein of unknown structure, the ∼225 kDa otoferlin, a Ca^2+^ sensor molecule that is essential for synaptic sound encoding^45^. The outlines provided by ONE microscopy imaging strongly resemble the AlphaFold^46^ prediction for this protein (**Fig. 2g,h, Supplementary Fig. 10**). Moreover, scanning in both the axial and lateral dimensions, using confocal laser scanning microscopy, enabled us to obtain 3D information on single otoferlin molecules (**Fig. 2i**). At the opposite end of the Ca^2+^ sensor size spectrum, we sought to visualize the small (∼17 kDa) protein calmodulin, expressed as a GFP chimera. To our surprise, even for such small particles, it was possible to observe dynamic changes in their shape upon Ca^2+^ binding (**Fig. 2j-n**). We applied both heat denaturation and proteinase K treatments for the homogenization of calmodulin, to test whether these methods would lead to different results. The proteinase K presumably removes all amino acids that are not anchored into the gel, and is therefore more aggressive than the heat denaturation^47^. However, both methods resulted in similar observations for calmodulin, implying that both can be used for observing the shape of purified proteins.

**Figure 2.**
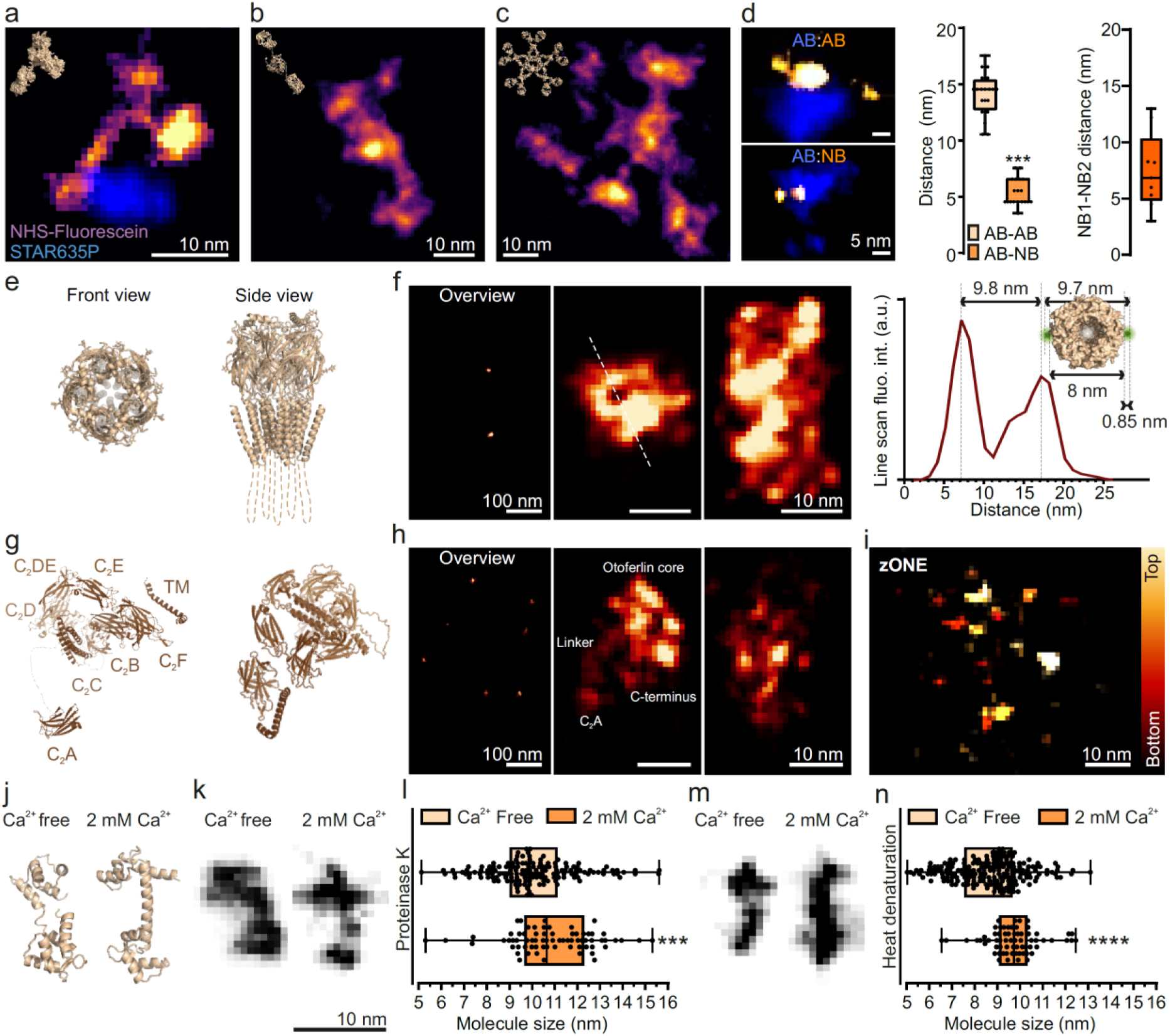
ONE analysis of single molecules. **a-c**, Images of isolated immunoglobulins (IgG, IgA, IgM), labelled with NHS-ester fluorescein after homogenization. The IgGs were secondary anti-mouse antibodies, carrying STAR635P (blue). **d**, Distances between fluorescently-conjugated IgGs and fluorescently-conjugated secondary antibodies (top) or secondary nanobodies (bottom). In panels a-d we used a different fluctuation analysis, termed temporal radiality pairwise product mean, TRPPM, unlike the temporal radiality auto-cumulant to the order of 4 (TRAC4)^6^ approach used in most other figures. Unlike TRAC4, which aims to separate the individual fluorophores, TRPPM enhances the cohesiveness of the fluorophores decorating the single antibodies, resulting in cloud-like signals whose distances are easily measured (N = 20/19, for AB:AB/AB:NB). Right panel: distance between the two secondary nanobodies binding single IgGs (N = 9). **e-f**, A TRAC4 analysis of purified GABAA receptors. Line scans across specific profiles seem to detect the receptor pore. **g**, AlphaFold-predicted model of otoferlin structure. **h**, ONE examples of otoferlin images. **i**, Z-axis ONE imaging, indicating that two components (presumably C2A and TM domains) are relatively far from the main body of the molecule. **j-k**, 1CLL and 1CFD PDB structures of the Ca^2+^ sensor calmodulin, in presence or absence of its ligand, respectively, along with ONE images, after proteinase K-based homogenization and expansion (see Supplementary Fig. 11 for more images). **l**, The expected elongation by ∼1 nm was reproduced (*p* = 0.0006, Mann-Whitney test; N = 70-155). **m-n**, Similar analysis, after homogenization using autoclaving (*p* <0001, N = 66-197). The box plot shows the medians, the 25^th^ percentile and the range of values.

### Protein averaging using ONE microscopy

One important challenge in ONE microscopy is to obtain deeper insight in the protein structure, beyond imaging the rough shape of an individual protein. In principle, one should be able to average the images of multiple proteins, but many difficulties are apparent, from our low resolution in the axial direction, which implies that every image is a projection of a relatively large volume, to difficulties in determining the orientation of the particles. Nevertheless, we attempted this approach, relying on the GABA_A_ receptor, in complex with GABA_A_R-binding nanobodies^48^ carrying fluorophores. These nanobodies recognize the receptors *in vitro*, providing the expected ONE images (**Supplementary Fig. 12**). Selected views fit very well with cryo-ET images of the receptor (**Supplementary Fig. 13**).

To generate an average receptor image, we used an automated particle detecting routine to find about 5000 particles, removing the ones with any signs of uncompensated drift (for drift examples, see **Supplementary Fig. 14**). We then inspected visually all of the remaining particles, to choose those that appeared to be in a “front view”, showing a reasonably round appearance, with nanobodies placed at the edges of the receptor (1225 particles). These particles were subjected to an analysis of the peaks of fluorescence, which were then mapped into one single matrix, which represents the “averaged receptor”. All sets of positions were rotated so that the nanobodies point “up” (**Fig. 3**).

As indicated by the different examples, not all receptor particles are decorated by five nanobodies, due to various issues, ranging from incomplete nanobody binding to the loss of fluorophores during the anchoring and expansion steps. This implies that the averaging procedure induces a bias for the nanobody image, since the nanobody with the highest fluorescence will be the one placed in the “up” position. This type of averaging will result in a prominent labelling for the nanobodies in this position, and little else in the nanobody channel. Even if more nanobodies are always present on the same receptor, small imprecisions in their placement will make them not align properly in the average image, resulting in one bright nanobody, up, and a blur elsewhere.

**Figure 3.**
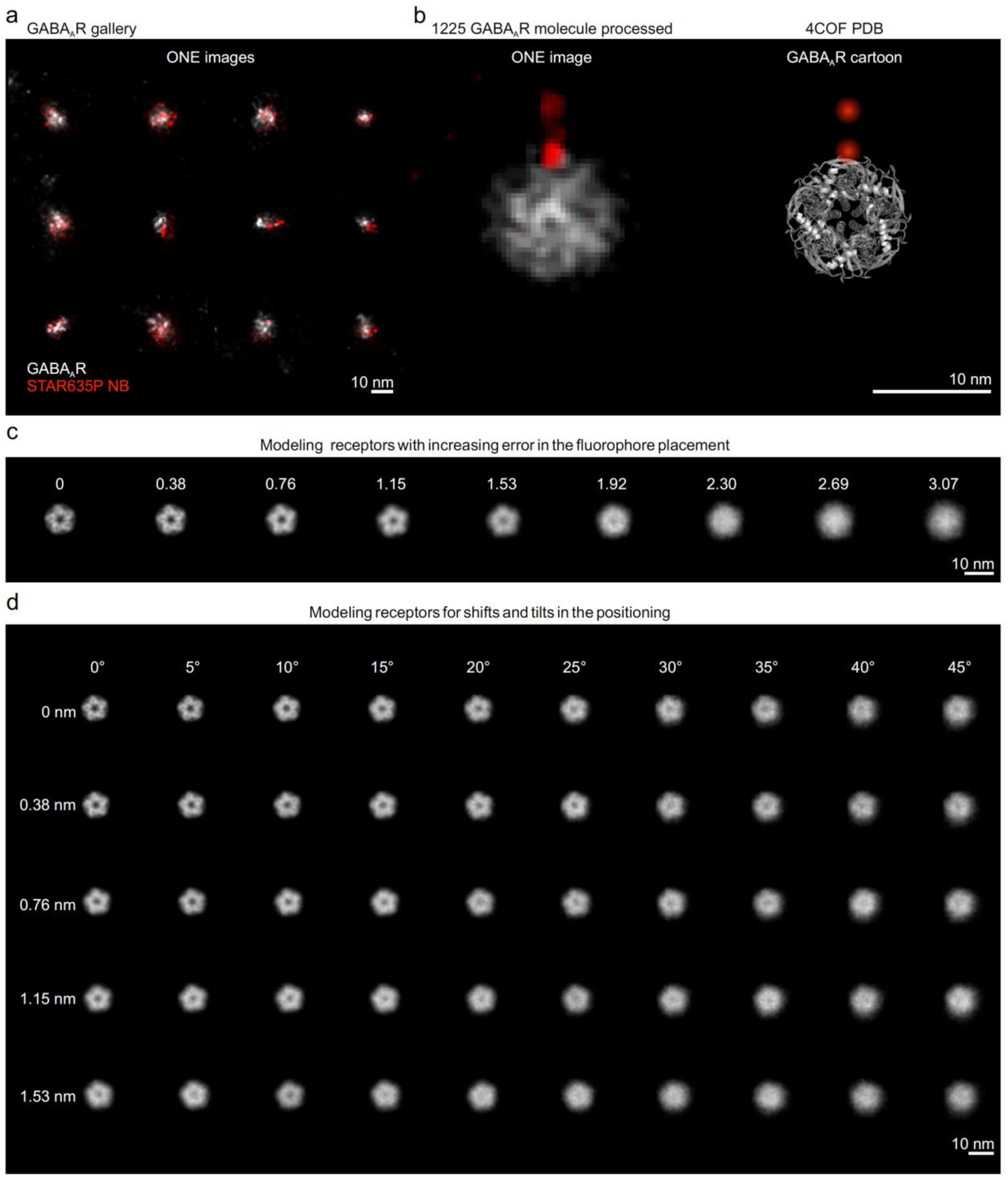
Averaging of GABAAR ONE images. **a,** A gallery of GABAAR/NB ONE images. **b,** An average view generated by processing 1225 receptor images. **c,** Receptors were modeled with increasing errors in the fluorophore placement. The error is indicated below above each receptor, in nanometers. The pore of the receptor is no longer visible when the fluorophores are displaced by 2 nm from their original positions. This analysis assumes that all receptors we analyzed are placed in a perfectly vertical position, and that there is no error in the identification of the receptor center. Neither of these assumptions is reasonable. **d,** The receptors were modeled with a variable shift in their lateral positioning and in their vertical tilt angles. The angles and placement errors are indicated above and to the left, respectively. Here we assume that fluorophore placement is perfect. This analysis reveals the fact that minor errors in the receptor orientation would lead to the same type of image as we obtained in panel b.

The results are nonetheless satisfactory. The average receptor obtained has a pore in the middle, and has the nanobody at a reasonable position. Two main fluorophore positions appear on the nanobody, at the expected distance from each other, exactly as predicted by the double labeling of the nanobody (two fluorophores on each nanobody). To determine whether this average view is “good enough”, we modeled receptor images, starting from the Cryo-EM coordinates of the receptors. Our models indicate that, in a perfect world, the blur observed in our receptor would come from a placement error, for each fluorophore, of about 2 nm (**Fig. 3c**). However, taking into account the fact that our 1225 particles have slightly different positions and tilt angles (**Fig. 3d**), the average receptor image is as good as it could be expected (**Fig. 3b**).

Overall, this approach indicates that single-particle averaging is indeed possible, albeit it would benefit greatly from increased resolution in the axial dimension, which can be achieved, for example, by using ExM protocols with expansion factors beyond 10x^49, 50^. This would enable us to determine better the position and tilt of each particle, thereby improving the results.

In principle, a similar procedure could be performed using purified tubulin assembled into microtubules *in vitro*. Unfortunately, this is an incredibly difficult experiment, since the microtubules are unstable *in vitro*. If they are thoroughly fixed, using glutaraldehyde, then they barely link to the gel, since majority of amines are modified by the fixative. If they are poorly fixed, with PFA, they link into the gel, but only in part, since a degree of microtubule depolymerization is still observed, leading to a destruction of the structure, quite visible both before and after expansion (**Supplementary Fig. 15**). However, we did succeed in averaging the free tubulin dimers that result from the depolymerization of the microtubules, and found that they averaged to an object that is similar to the known structure of this dimer (**Supplementary Fig. 15**).

We also extended this analysis to actin. To use a natural, rather than *in vitro*, system, we turned to cell cultures subjected to detergent extraction during fixation. This procedure results in the preservation of actin filaments^51^, which could then be analyzed in ONE microscopy. The images reproduce the known size of the actin filaments and the distance between the actin subunits, as well as providing views of the filament pitch (**Fig. 4**).

**Figure 4.**
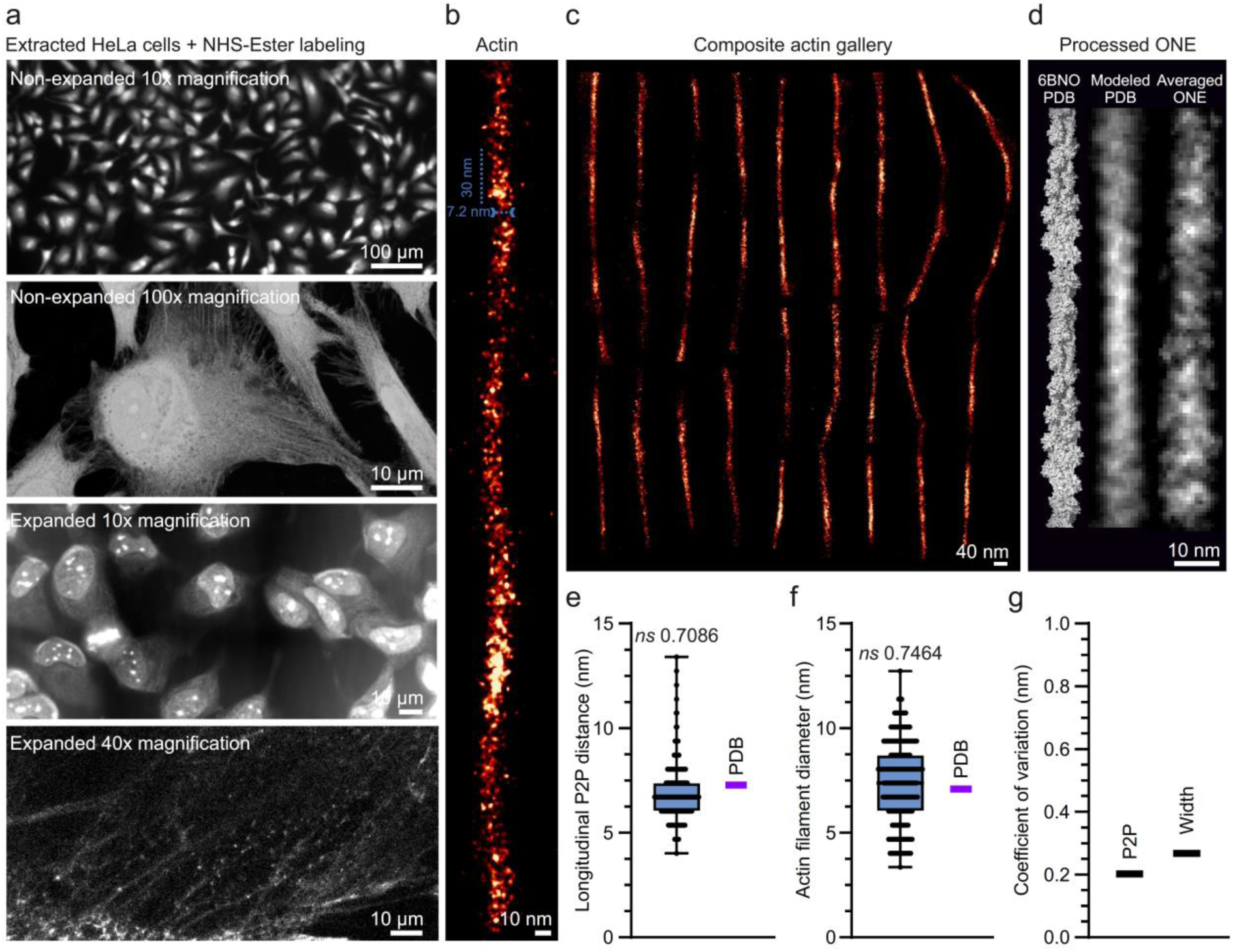
ONE imaging of cytoskeletal components, in NHS-ester labelled cells. **a,** HeLa cells were subjected to extraction according to a previously published protocol^51^. Panels from top to bottom show an overview of non-expanded, extracted HeLa cells labelled with NHS-ester chemistry (10x objective), followed by a higher magnification view and by two views of cells expanded after extraction. **b,** An exemplary image of a cytoskeletal filament. **c,** Composite actin gallery. **d,** A PDB cartoon, followed by a modeled averaged PDB, depicting a best-case scenario for ONE microscopy of actin fibrils under ideal conditions, and the achieved ONE image reconstructed from 47 different actin regions. **e, f,** & **g,** A set of measurements that were applied over actin filaments, quantifying the longitudinal peak-to-peak distance along the filament (distance between actin monomers), the actin filament width, and the coefficient of variance of these two measurements, respectively. These values resemble well the known dimensions from actin PDBs (magenta lines). N = 198 and 130 for the first graph and second graph, respectively; 2 independent experiments.

### Visualization of synaptic proteins

We next tested the performance of ONE microscopy in cultured neurons, focusing on the synaptic transmission machinery. To test whether a simple epifluorescence microscope could be obtained for ONE microscopy in samples where molecular resolution is not absolutely needed, we turned to an Olympus iX83 TIRF microscope, equipped with an Andor iXon Ultra EMCCD camera. This microscope should provide a maximal resolution of ∼3 nm, since the obtained resolution is limited by the pixel size, which reaches ∼1.5 nm, when the radiality and expansion factor are taken into consideration (the expected resolution cannot be better than 2-fold the pixel size^52^).

Synaptic vesicles fuse to the plasma membrane to release their neurotransmitter contents, and their molecules are afterwards endocytosed, following different pathways^53^. A significant fraction of the vesicle proteins are found on the plasma membrane, forming the so-called “readily retrievable pool”^54^. It is unclear whether these proteins are grouped in vesicle-sized patches, or whether they are dispersed on the presynaptic membrane. We investigated this here, using a fluorescently-conjugated antibody directed against the intravesicular domain of the Ca^2+^ sensor synaptotagmin 1 (Syt1), an essential component of the vesicles. The labeling density was sufficiently high to reveal that endocytosed vesicles have the expected circular shape (**Fig. 5a**). Turning to the readily retrievable pool, we found that the molecules were grouped in areas of size consistent with theoretical expectations for single fused vesicles, with copy numbers similar to the values expected for this molecule (7-15 per vesicle^55, 56^), taking into account the fact that one antibody can bind up to two Syt1 molecules (**Fig. 5b,c**). Removing cholesterol from the plasma membrane, using methyl-β-cyclodextrin (MβCD), forced the dispersion of synaptotagmin molecules (**Fig. 5b,c**), albeit it left the overall synapse organization unaffected (**Supplementary Fig. 16**). The numbers of antibodies were not estimated by counting the individual antibody spots, which is a difficult procedure under the lower resolution provided by the epifluorescence microscope, but by analyzing the fluorescence intensity of single vesicles, compared to the intensity of individual antibodies (**Supplementary Fig. 17**). One important consideration is whether the intensity of specific spots can be used reliably in ONE images. While the current analysis suggests this, we also performed a more formal analysis of spot fluorescence in single-molecule images, which are the most challenging for ONE analyses. As shown in **Supplementary Fig. 18**, ONE does result in signal variations, but they do not impede the differentiation of real signals from background values.

**Figure 5.**
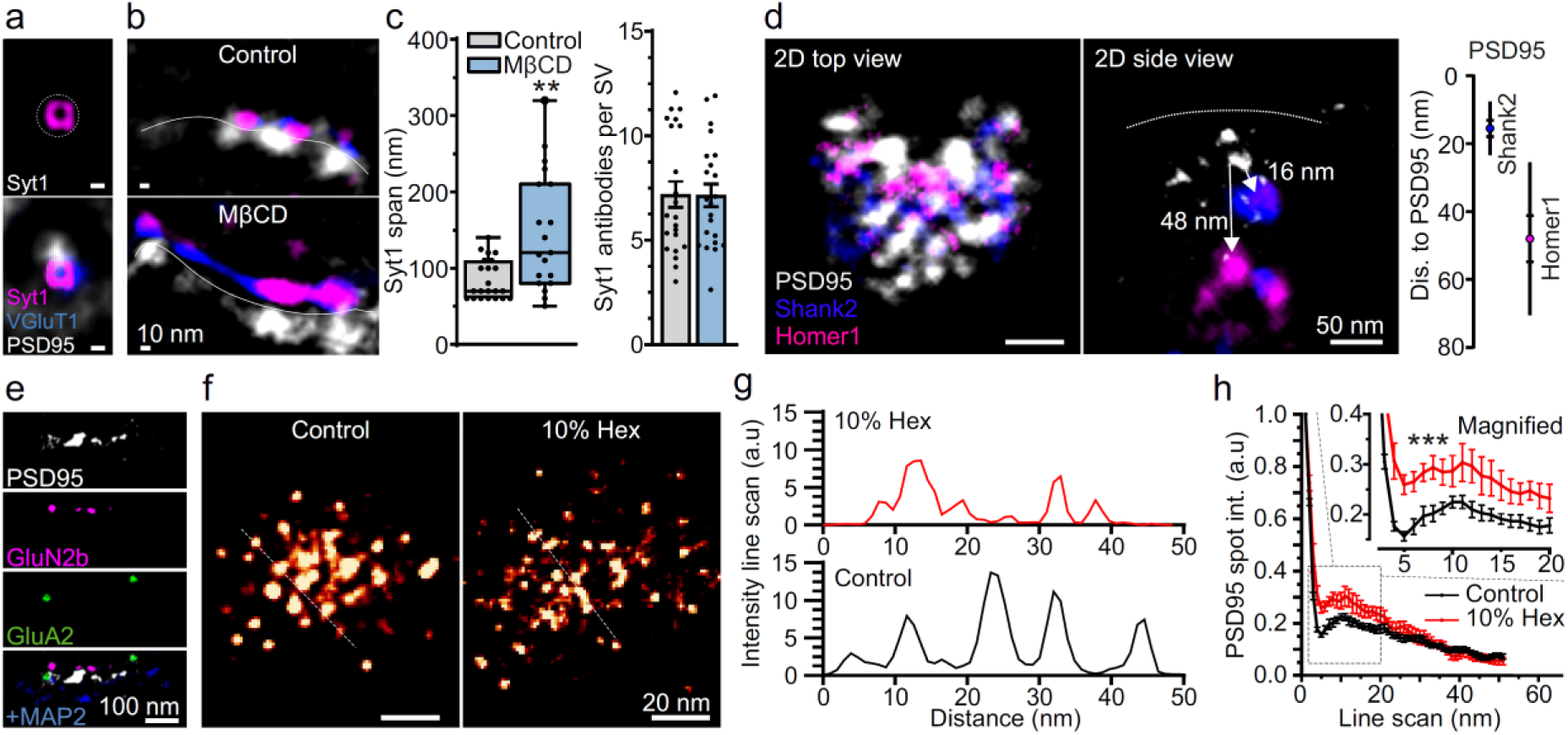
ONE reveals pre-and postsynaptic structures. **a-c**, Synaptic vesicles were labelled live using an antibody against a luminal epitope of synaptotagmin 1 (Syt1, magenta). The vesicular glutamate transporter (vGluT1, blue) and PSD95 (gray) were immunostained using an antibody and a nanobody, respectively. **a**, Recently endocytosed vesicle exhibiting circular morphology. **b**, Readily retrievable pool molecules form patches containing Syt1/vGluT1 (top), which are dispersed by cholesterol extraction using MβCD (bottom). **c**, MβCD causes molecules to spread across larger areas (left: N = 22-19, 2 independent experiments, *p* < 0.0044, Mann-Whitney test; right: N = 22-22, 2 independent experiments, *p* = 0.8937), although the signal per vesicle (the Syt1 copy number) remains unchanged. **d**, A visualization of PSDs (top and side views), after immunostaining PSD95 with the same nanobody used in a-c, and Shank2 and Homer1 with specific antibodies. The graph indicates the axial positioning, which agrees well with the literature^57^. N = 11 measurements for each protein, 2 independent experiments; symbols show the medians, SEM and SD. **e**, Side view of a postsynapse displaying PSD95, MAP2 and two glutamate receptors (GluA2, AMPA type, and GluN2b, NMDA type). **f**, ONE images of PSD95 (top views), with, or without, the addition of 10% 1,6-hexanediol (Hex). **g**, Line scans through the PSD95 staining shown in panel f. **h**, An analysis of PSD95 spot profiles; N = 10-7 synapses, Friedman test followed by Dunn-Sidak testing, *p* = 0.0027; the error bars show the SEM. For details on the analysis, see Supplementary Fig. 19.

In the post-synaptic compartment, we could confirm known organization principles, including the layered aspect of the postsynaptic density (PSD), in which molecules like PSD95, Shank and Homer occupy different positions in the axial direction (**Fig. 5d**), or the clustered distribution of postsynaptic receptors, with NMDA receptors typically observed in more central locations than AMPA receptors (**Fig. 5e**; see also ^58^; see **Supplementary Fig. 20** for a quantification of the receptor positions). These experiments confirm the ease with which ONE microscopy provides multicolor super-resolution in crowded cellular compartments.

### PSD95 has a retiolum-like organization

The fine structure of the PSD, or even its very existence, is a matter of considerable debate. The current prevalent view is that the PSD is maintained by liquid-liquid phase separation (LLPS)^59^, which, intuitively, implies an amorphous organization. To test this hypothesis, we immunostained PSD95 in cultured hippocampal neurons, using a specific nanobody (**Fig. 5f**), and we switched back to confocal microscope equipped with resonance scanning, to improve image resolution. PSD95 appears to be organized in a quasiregular lattice, a conclusion that was strengthened by overlaying PSD images to obtain average views (**Extended Data Fig. 6**). An analysis of the distances between PSD95 spots revealed that they have a preferred spacing of ∼8-9 nm, which is significantly different from a random distribution (**Extended Data Fig. 7**). A similar result was obtained when using a Ripley curve-like analysis (**Fig. 5g,h**; see **Supplementary Fig. 19** for details). To test the stability of the supramolecular PSD95 arrangements observed, we incubated the cells with 1,6-hexanediol, an alcohol that has been often used to cause the dispersion of liquid phases^60^. This treatment readily dispersed other components of the PSD, as Homer1 and Shank2, but did not affect PSD95, which appeared to remain unchanged at the confocal imaging level (**Extended Data Fig. 7d**). This was no longer the case when samples imaged in ONE microscopy, as the 1,6-hexanediol treatment caused the PSD95 arrangements to lose much of their regularity (**Fig. 5f-h**). Importantly, this organization of PSD95 is not an artefact due to steric hindrance induced by the nanobodies, since post-expansion labeling results in virtually identical PSD images (**Supplementary Fig. 21**).

These results suggest that the PSD95 positioning may only be partially, but not fully, controlled by LLPS mechanisms. We propose the term “*retiolum*” for this nanoscale PSD patterning, a Latin term describing small string nets with knots at regular intervals^61^. These fine details of PSD95 organization are fundamentally different from the PSD95 nanodomains observed in the past by super-resolution imaging of antibody-based immunostaining^58, 62^. The nanodomains observed in the past are most likely a result of the limited resolution of the respective technologies, as demonstrated in **Extended Data Fig. 8**.

While all of the results presented above on synaptic proteins were derived from neuronal cell cultures, we would like to point out that ONE microscopy can also be applied to tissue samples to investigate such protein arrangements, as we performed for brain slices of more than 200 µm in thickness (**Extended Data Fig. 9**).

### Towards Parkinson’s Disease diagnostics

While ONE microscopy provides substantially more details than more established super-resolution methods X10 ExM and STED microscopy, or even their combination (**Supplementary Fig. 22**), its precise resolution remains relatively difficult to estimate. To provide a measurement for different structures, we performed a number of FRC analyses, as indicated in **Supplementary Fig. 23**. As shown in the last panels of this figure, the average FRC value is below 1 nm when suitably small pixels are used in the ONE analysis. This observation is confirmed by our ability to measure intra-molecular distances as low as 0.5 nm, within single molecules (**Supplementary Fig. 24**).

Overall, while precise claims of resolution are difficult to make for this imaging scale, these observations suggest that ONE microscopy should be able to tackle real-world questions. Another pre-condition for such analyses is that the expansion is sufficiently precise (isotropic) to not modify the protein shapes drastically. This is indeed the case, as demonstrated by a large number of measurements, performed across a range of different proteins (**Supplementary Fig. 25**).

Therefore, we next sought to address a pathology-relevant imaging challenge by ONE microscopy. Parkinson’s disease (PD) is a neurodegenerative disease characterized by the accumulation of aggregates composed of several proteins, of which alpha-synuclein (ASYN) is the most prominent^63^. In the cell, ASYN can exist as a monomer, or assemble into species of different sizes, in a process believed to be relevant in the context of PD. These species include soluble oligomers and fibrils (e.g.^64^). In patient-derived samples, such as cerebrospinal fluid (CSF), detecting disease-relevant ASYN species is thought to be a relevant marker for PD diagnostic purposes and for monitoring disease progression, since the measurement of ASYN levels alone has proven to lack diagnostic relevance^65–67^.

We explored the diagnostic potential of imaging ASYN assemblies in the CSF of PD patients *versus* controls (**Supp. Table 1**). A nanobody able to bind ASYN^68^ was used because full-length immunoglobulins only provide relatively poor labeling due to their large size (**Fig. 6a, Supplementary Fig. 26**). Different types of ASYN assemblies could be detected (**Fig 4b**) and PD patients had higher levels of oligomer-like structures (**Fig. 6c,d**). We classified the structures observed and noticed that the “very large” assemblies (>200 nm in length, >50 nm in width) were found at similar frequency in PD patients and controls (**Fig. 6e**). The same was observed for assemblies in the 50 to 200 nm length range (**Fig. 6e**). However, this was not the case for the smaller, oligomer-like assemblies. Some resembled strikingly polymorphic ASYN assemblies that have been recently described by Cryo-EM^69, 70^, while others had an annular organization as observed in the past by negative stain transmission EM or Cryo-EM^71, 72^. All oligomer-like species were significantly more abundant in PD CSF than in control samples (**Fig. 6f**), and their cumulative analysis, which alleviates ambiguities due to imperfect classification, resulted in a good discrimination of PD patients and age-matched controls (**Fig. 6g,h**; for overviews of more ASYN objects see **Extended Data Fig. 10**). We conclude that the analysis of ASYN aggregates by ONE microscopy is a promising procedure for PD diagnosis.

**Figure 6.**
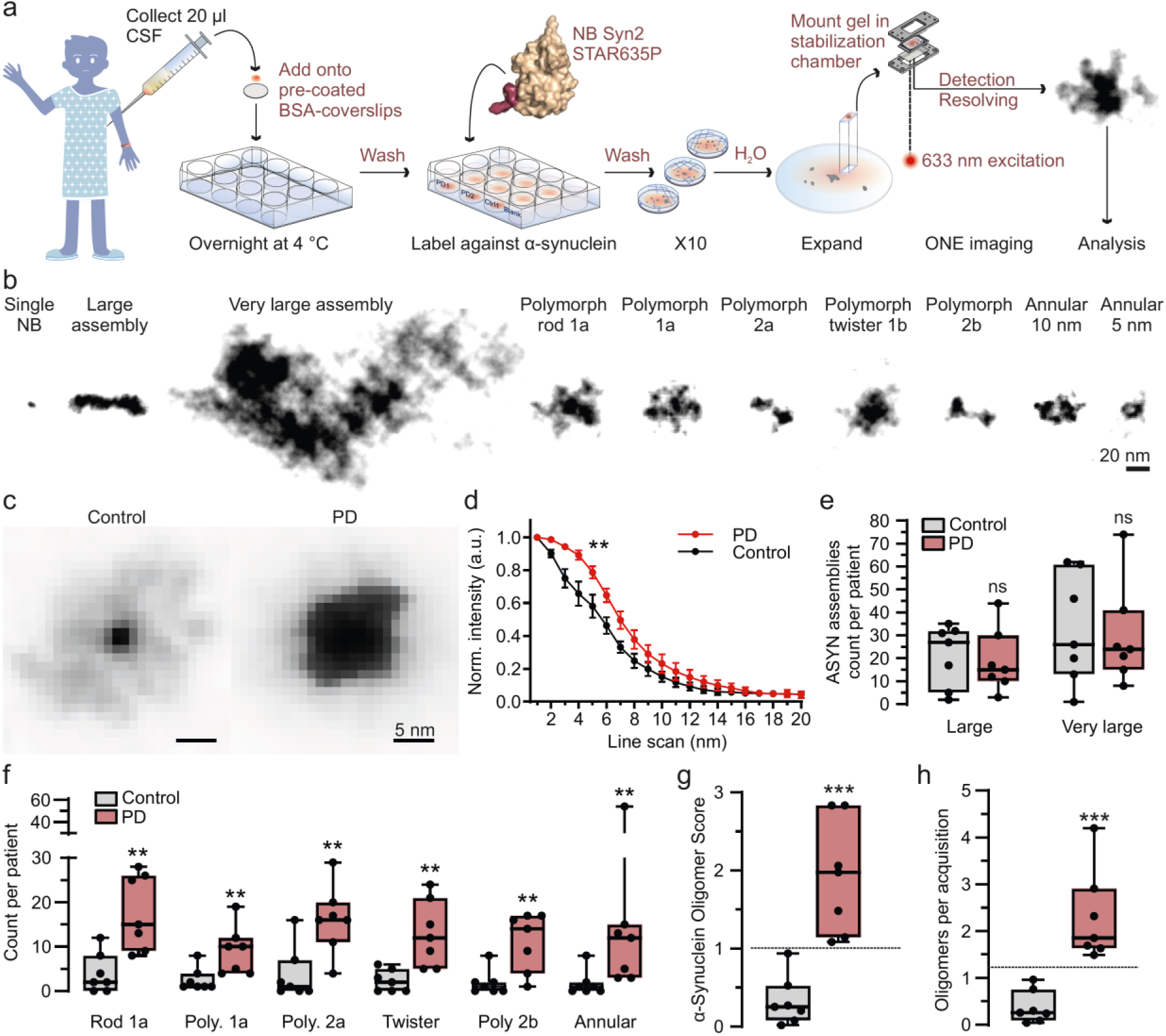
A promising avenue for Parkinson’s Disease (PD) diagnostics. **a**, Cerebro-spinal fluid probes were obtained from PD patients and controls, and 20 µl amounts were placed on BSA-coated coverslips, followed by ONE imaging, after immunolabeling ASYN using a specific nanobody^68^. **b**, A gallery of typical ASYN species observed in the CSF samples. **c**, Average ASYN assemblies from a PD patient and a control. **d**, An analysis of the spot profiles detects significant differences, with the average control object being smaller than the average PD object. All ASYN assemblies for the control and PD patients were averaged, from 3 independent experiments, Friedman test followed by Dunn-Sidak, *p* = 0.0237; errors show SEM. **e**, An analysis of the numbers of the larger assemblies in CSF samples. No significant differences, Mann-Whitney tests, *p =* 1, *p* = 0.7104. **f**, An analysis of the numbers of oligomers in CSF samples. All comparisons indicated significant differences, Mann-Whitney tests followed by a Benjamini-Hochberg multiple testing correction with FDR of 2.5%; *p* values = 0.0105, 0.0023, 0.0111, 0.0012, and 0.0012, in the respective order of data sets. **g**, Analyses of the numbers of oligomers, as a proportion of all ASYN assemblies analyzed (left), or as numbers per acquisition in (**h**). Both procedures discriminate fully between the PD patients and the controls. For the second procedure, the lowest PD value is 50% larger than the highest control. N = 7 PD patients and 7 controls, Mann-Whitney test, *p*<0.0001 for both **h** and **g**.

Finally, ONE can also analyze medical samples that have been fixed for prolonged (and uncontrolled) time periods, as observed for chemically fixed blood sera from COVID-19 patients, obtained commercially (**Supplementary Fig. 27**). Here we observed that the patient IgGs clustered in specific regions of the viral particles, whose detailed composition could be targeted in the future.

For completeness, we performed an FRC analysis of the ASYN and virus samples, presented in **Supplementary Fig. 28**.

### Multi-laboratory applications of ONE microscopy

An important issue for any new technology is its wide applicability, in multiple laboratories. To test this issue, we collaborated with academic laboratories in Homburg (Germany), Würzburg (Germany) and at MIT (USA), and with the industrial laboratory of microscope developer Leica Microsystems (Mannheim, Germany). We focused on GABA_A_ receptors and tubulin, samples that were well described in the rest of the work (**Supplementary Figures 29** to **33**). We were able to show that ONE can be applied in different laboratories, with some of the experiments even surpassing our original applications by either using larger expansion factors (MIT laboratory, 20x expanded immunostained microtubules in cultured cells; **Supplementary Fig. 31**; post-expansion stained bassoon in x20 expanded mouse brain tissue, **Supplementary Fig. 33**) or by using faster scanning to allow volumetric zONE imaging in two color channels (Leica Microsystems laboratory; **Supplementary Fig. 29**).

## Discussion

Here we show that a fluorescence microscopy procedure based on a combination of X10 ExM and radial and temporal fluorescence fluctuation analysis (SRRF) can provide sufficient spatial resolution to analyze single protein shapes. ONE was implemented in five laboratories on different microscopes. Therefore, ONE microscopy makes super-resolution imaging broadly available, in a fashion that has always been a primary goal of ExM^73^. Moreover, no special handling, unusual fluorophores or reagents are necessary. The ONE data processing is relatively fast because the SRRF procedure is performed in minutes (see Supplementary File 1 for details). The initial immunostaining and expansion procedures take, combined, 3-4 days, while imaging individual regions of interest only takes between 35 seconds and 2 minutes, depending on the number of color channels.

The ONE axial resolution surpasses that of confocal microscopy by one order of magnitude, owing to the 10x expansion factor. Further improvements of axial resolution could be introduced in the future, through total internal reflection fluorescence (TIRF), lattice light-sheet microscopy^74^, or multi-focus microscopy^7^. The only major limitation we see is that ONE cannot be applied to live samples, due to the ExM procedures. Therefore, the MINFLUX concept^8, 11, 75, 76^ is currently the only solution for live imaging in the very high-resolution domain. Nevertheless, future developments in ONE microscopy are likely to enable 3D structural analysis of proteins, either purified or in cells and tissue samples, at resolutions approaching electron cryo-microscopy and tomography techniques, at room temperature and at a fraction of the cost. Developments envisaged include refined anchoring chemistries, gels that are homogeneous to sub-nanometer levels, as well as imaging automation, to enable the analysis of tens of thousands of particles in a time-efficient manner.

Overall, we conclude that the ONE technology provides a simple, robust and easily applied technique, bridging the gap to X-ray crystallography and electron microscopy-based technologies.

## Extended Data Figure legends

**Extended Data Fig. 1.**
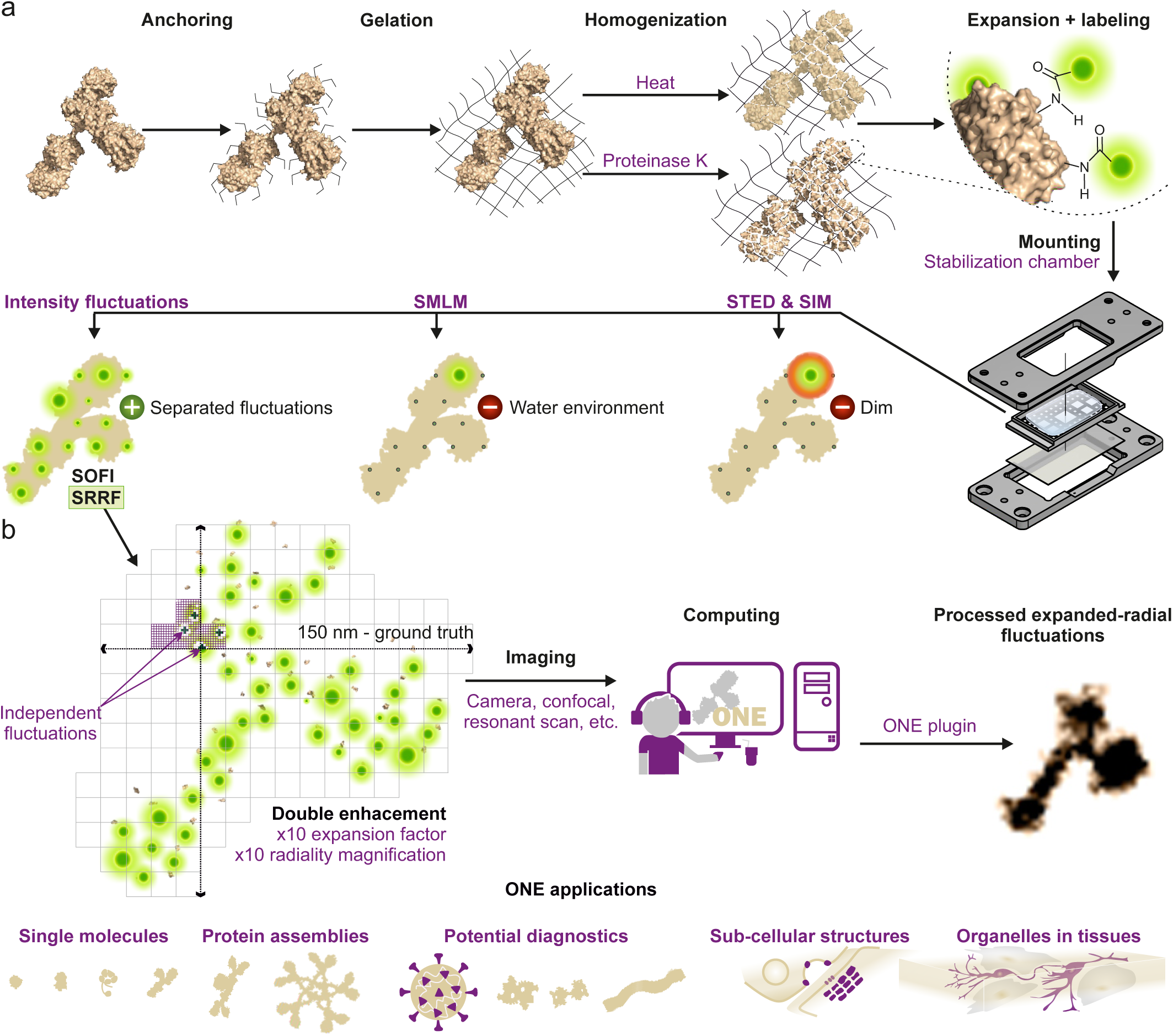
A general overview of the ONE microscopy approach. **a**, Biological samples are linked to gel anchors, relying on Acryloyl-X, followed by X10 gel formation and homogenization, either by proteinase K additions or by autoclaving in alkaline buffers. Full expansion is achieved by repeated washes, and is followed by mounting gel portions in a specially designed chamber. In principle, one could image the samples using different super-resolution procedures. Techniques benefitting from bright samples, as STED or SIM, suffer due to the fluorophore dilution induced by the expansion procedure. Techniques requiring special buffers (*e.g.* SMLM) are negatively affected by the water environment. In contrast, technologies relying on fluorophore fluctuations profit from the expansion, as the fluorophores are spatially separated and can fluctuate independently^13^. **b**, Repeated imaging is performed (up to 3000 images), in any desired imaging system (confocal, epifluorescence, etc.), to detect signal fluctuations, which are then computed using through a plugin (ONE platform) based on the SRRF algorithm, before assembling the final super-resolved images.

**Extended Data Fig. 2.**
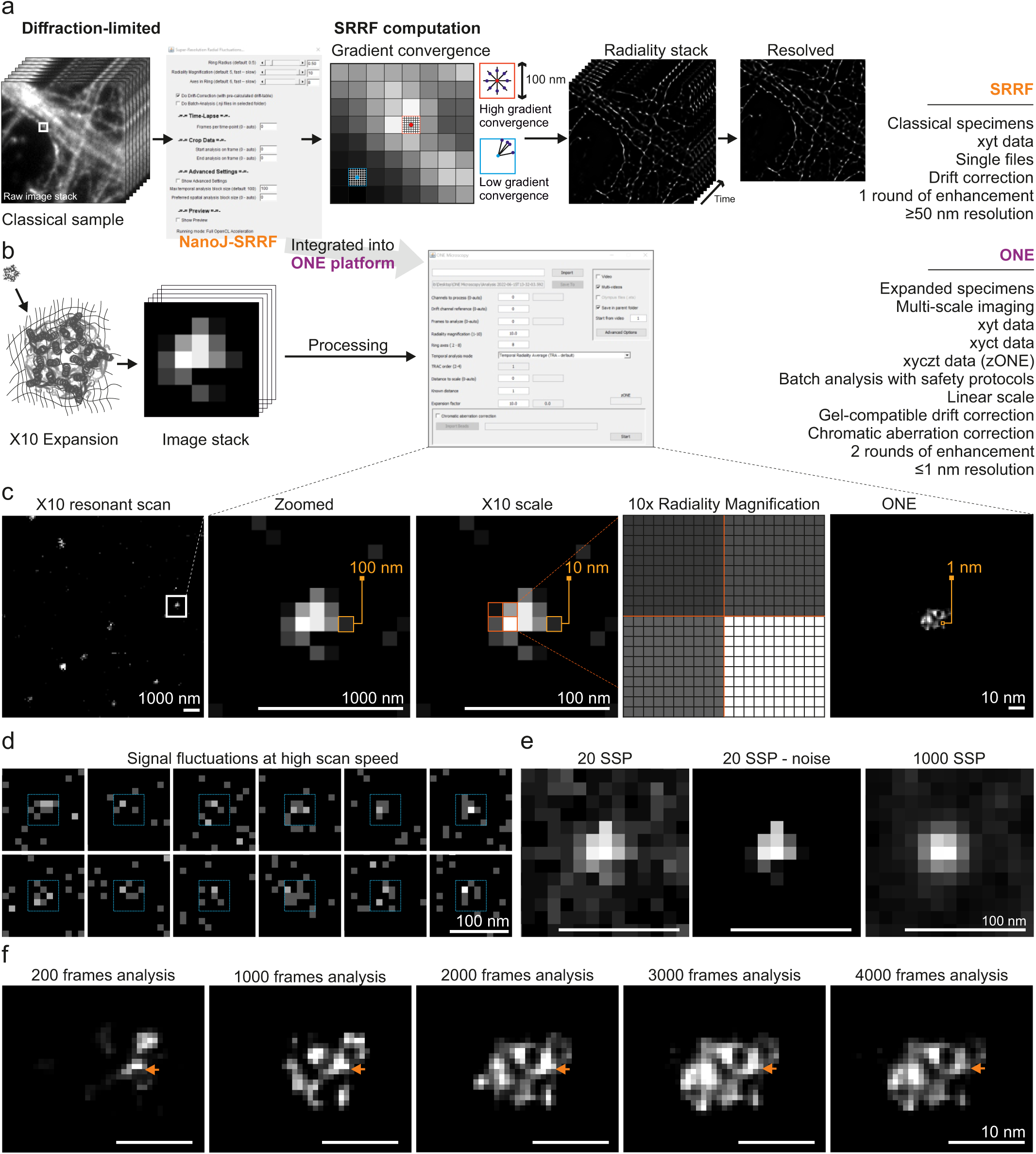
A detailed view of the ONE procedure. **a**, Processing a stack of diffraction-limited images with SRRF, based on the analysis of a gradient of convergence of sub-pixels over a radiality stack, results in super-resolved images with resolutions varying between 50-70 nm. **b**, The ONE procedure adapts the SRRF algorithm to expanded gels. **c-f**, A detailed explanation of the analysis procedure. **c**, A sample was fixed and expanded using a 10-fold expansion protocol (X10). The sample was then imaged using a resonant scanner on a confocal microscope. The zoomed-in view indicates one bright spot, whose size in real space is limited by diffraction to ∼200-300 nm, but represents a 10-fold smaller size in the pre-expansion space (see scale bars in the middle panels). Every pixel is then subjected to a 10-fold radiality magnification and is then subjected to the procedure explained in panels d-f, which provides the final, high-resolution image (right-most panel). **d**, Signal fluctuations are measured by imaging the sample repeatedly, using the resonant scanner (here at 8 kHz). **e**, A view of the overall signals, obtained by summing 20 of the fluctuating images (raw in the left-most panel, background-subtracted in the middle panel), or by summing 1000 images. **f**, Each image from series obtained as in panel b is subjected to a temporal analysis of fluctuating fluorophores, based on radiality magnification^6^, thereby providing a super-resolved image whose level of detail becomes optimal after ∼1500 frames.

**Extended Data Fig. 3.**
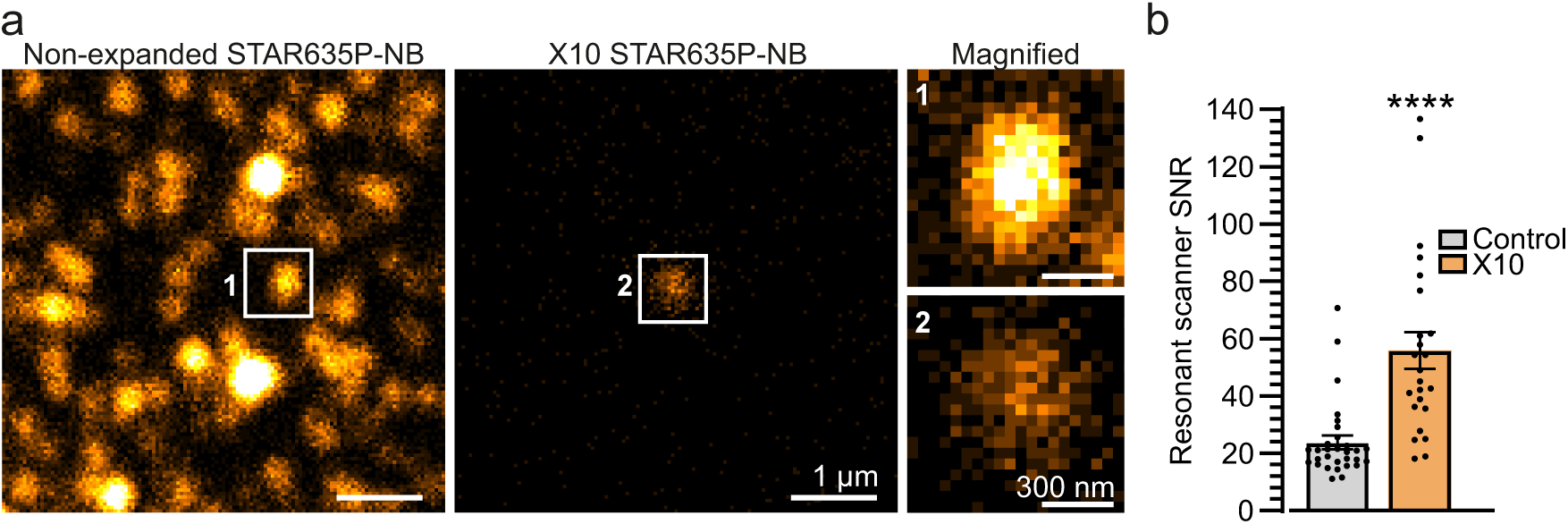
Expansion microscopy results in a higher signal-to-noise ratio. Expansion microscopy, which separates proteins of interest and removes much of the other cellular components (*e.g.* lipids, metabolites) should result in a higher signal-to-noise ratio (SNR). **a**, To test this, we analyzed here the simplest possible sample, consisting of Star635P-conjugated nanobodies on glass coverslips, or in expanded gels, using confocal microscopy, relying on analysis using a resonant scanner. **b**, The SNR of these samples increases by 2-fold, on average, after expansion. N = 30-24, *P* = 0.000001, Mann-Whitney Ranksum test.

**Extended Data Fig. 4.**
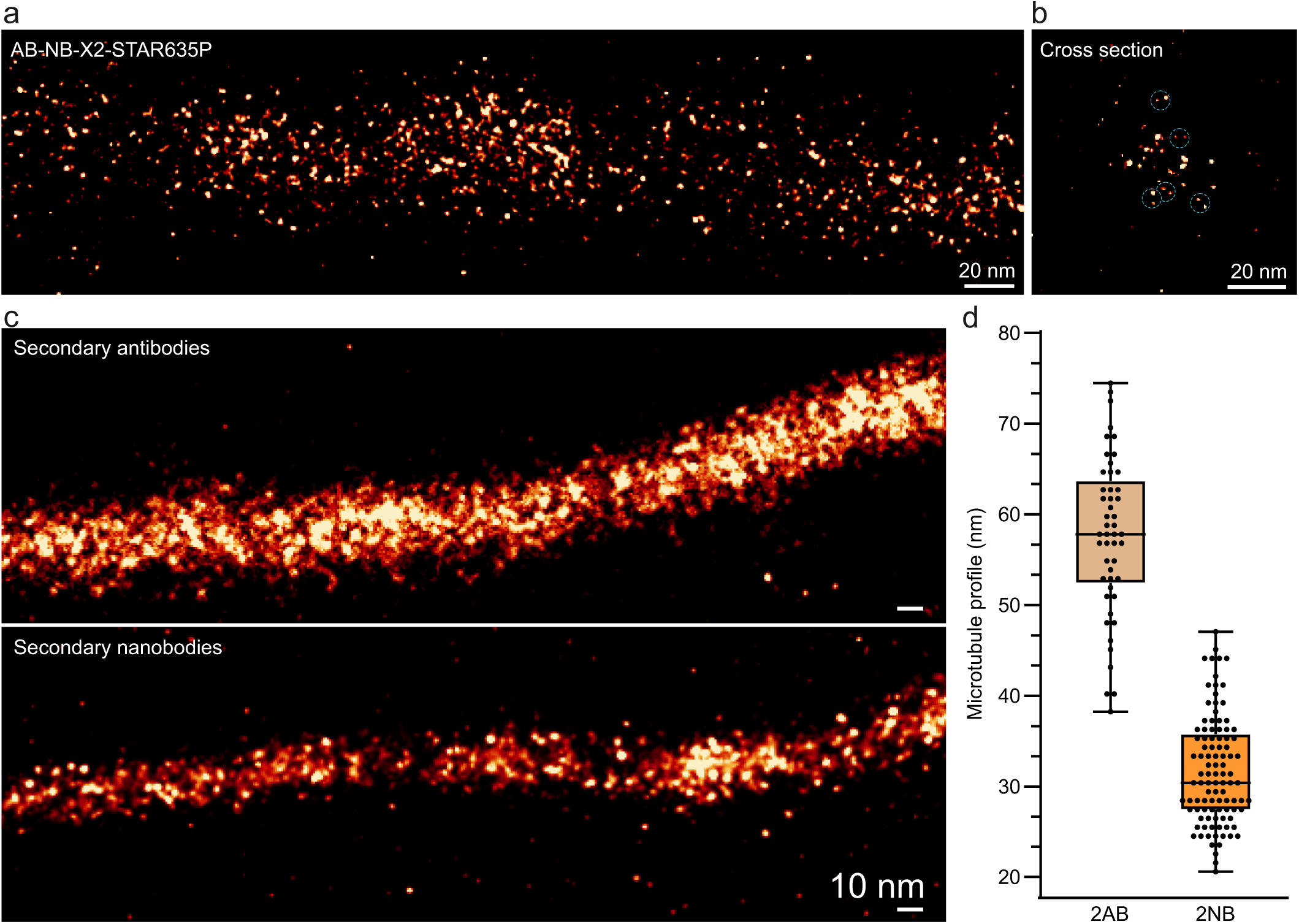
An analysis of tubulin immunostainings. **a** & **b**, An analysis of tubulin, following immunostainings relying on primary antibodies detected using Star635P-conjugated secondary nanobodies. While the overall signal distribution is similar to that obtained with secondary antibodies (Fig. 1), one can observe often pairs of fluorescent spots in very close vicinity (marked by dotted circles in the cross section), which probably represent the two fluorophores on each nanobody. For a formal analysis of this issue on different nanobodies, see Fig. 1 and Extended Data Fig. 5. **c**, Immunostainings relying on primary antibodies followed by secondary antibodies (upper panel) or by secondary nanobodies (lower panel). **d**, The graph shows the diameter of microtubules in when using secondary antibodies (left; N = 49 microtubule profiles) or secondary nanobodies (right; N = 101).

**Extended Data Fig. 5.**
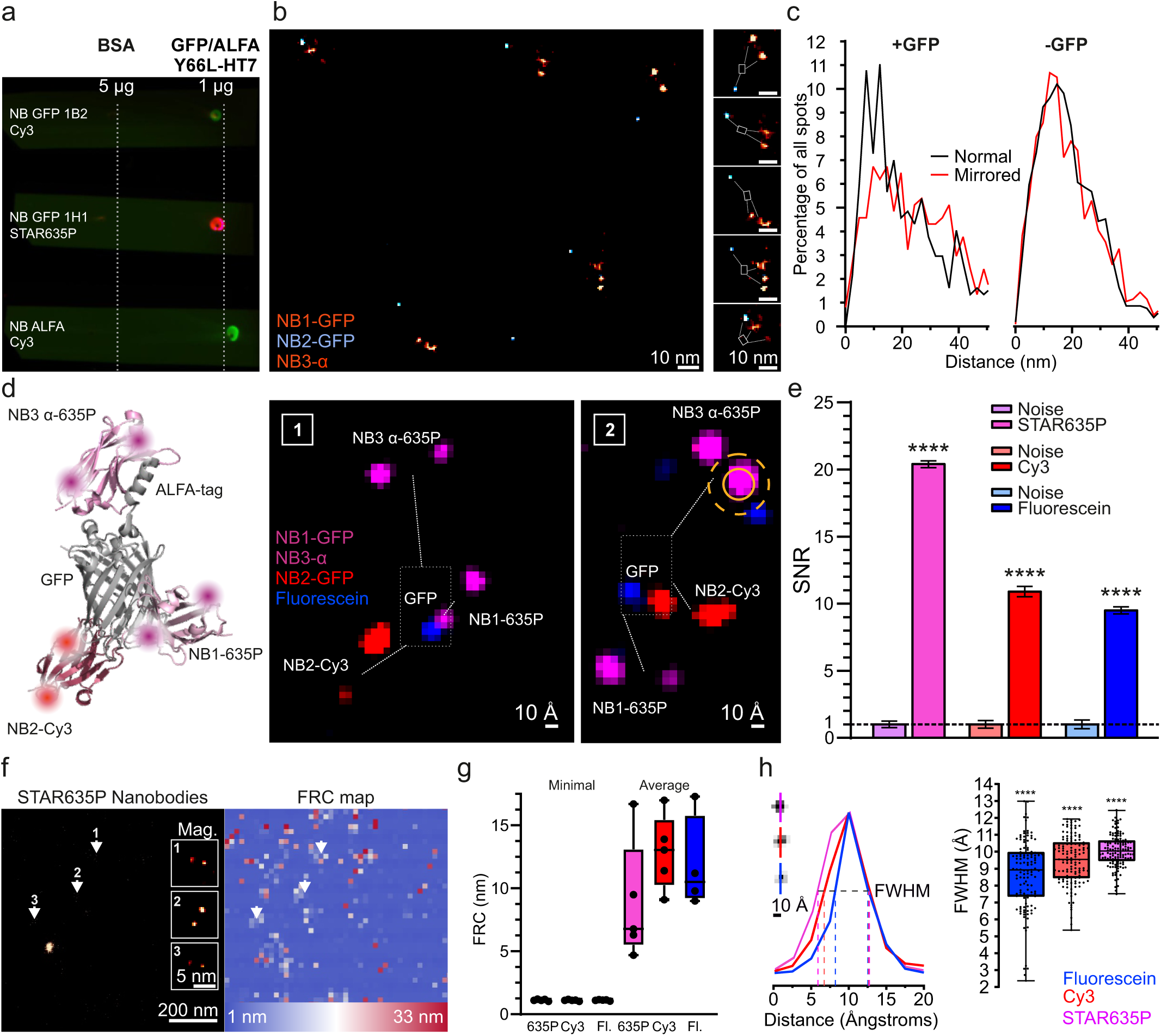
In-depth analysis of GFP-nanobody complexes. **a**, Dot blots to validate that each nanobody was binding specifically the TSR individually. Nitrocellulose membranes were spotted with TSRs and bovine serum albumin, as control, and the spots were revealed with the respective nanobodies, using a fluorescence scanner (GE-Healthcare AI 600). **b**, An overview of an image showcasing nanobodies bound to their GFP target. **c**, An analysis of distances from STAR635P to Cy3 nanobodies, in normal images or after mirroring one of the fluorescence channels, as a negative controls. The close-distance interval is largely removed by mirroring. N = 40-40 TSRs. Performing this in samples lacking the GFP, in which the nanobodies are randomly distributed, results in no differences between the normal and mirrored distributions. N= 40/40 images. **d**, Overview of the TSR using only two-color nanobody labeling (same as the one used in Fig. 5c,d), along with two different examples. The sample is also labelled using NHS-ester fluorescein, and a small pixel size (0.48 nm) is used, to enable the optimal visualization of the TSRs. **e**, An analysis of the signal-to-noise ratio of the TSRs, obtained by measuring the noise levels in the vicinity of the nanobodies. The noise levels are normalized to 1, implying that the normalized signal of the respective nanobodies now provides directly the signal-to-noise ratio. N = 20-18, 12-14, and 17-11 measurements, *P* < 0.0001, Mann-Whitney test. **f**, A Fourier Ring Correlation (FRC) analysis of nanobody images. **g**, The best and average resolutions obtained per image, in the different color channels (N = 4 to 5 analyses for each). **h**, To approximate the apparent resolution of the system, we drew line scans across spots and measured the full width at half maximum (FWHM) in curve fits executed on the line scans. The graph plots the FWHM of 129, 135, and 132 fluorescein, Cy3 and STAR635P line scans. The values are significantly different between the color channels. *p* < 0.0001, Kruskal-Wallis test. The box plot shows the median, 25^th^ percentile and the range of values.

**Extended Data Fig. 6.**
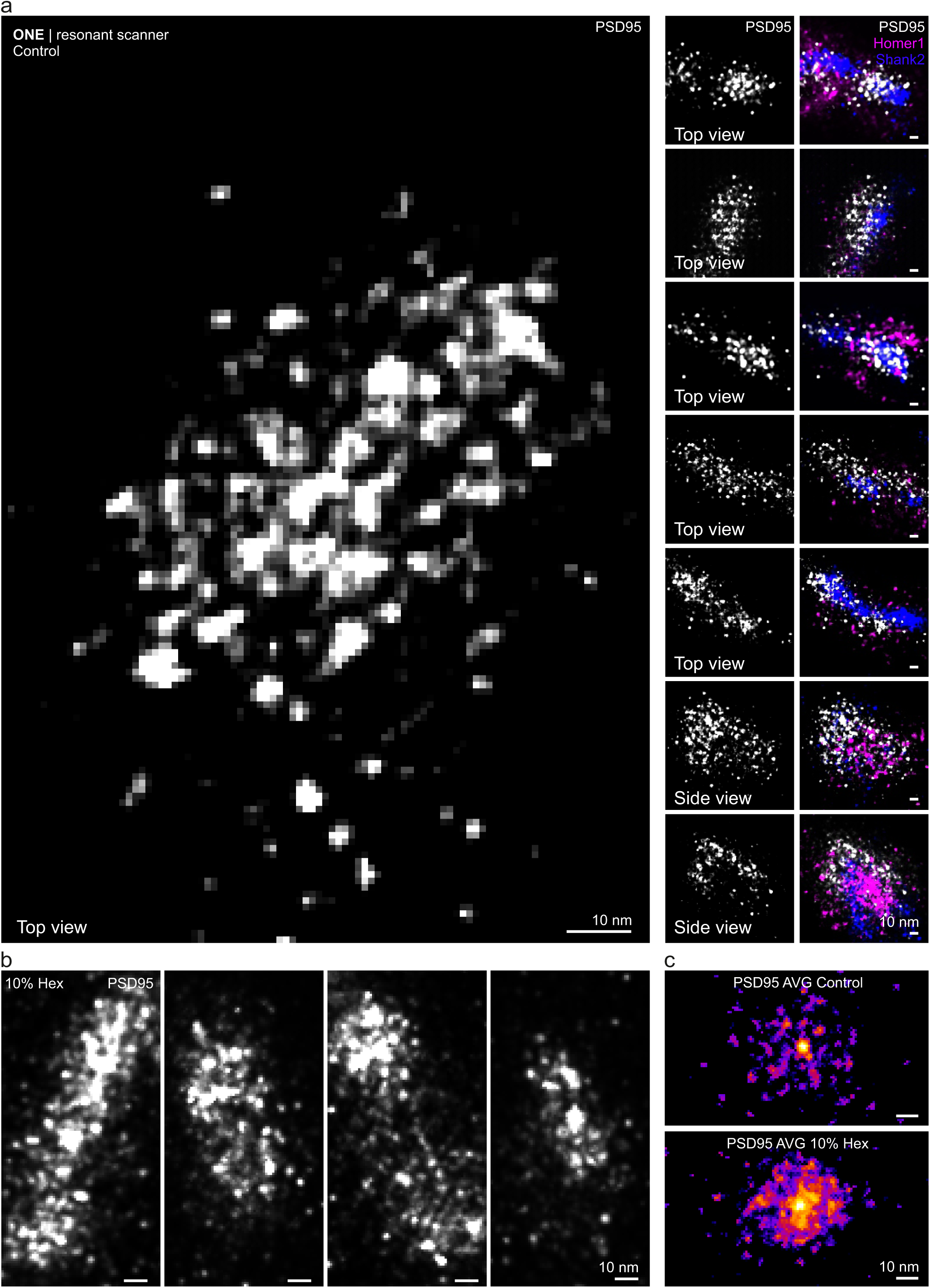
Further PSD examples. **a**, ONE imaging of PSDs, employing a resonant scanner and a final pixel size of 1 nm, to achieve a high resolution (same procedure and resolution as in Fig. 3f). **b**, Examples of PSD95 stainings, after treatment with 1,6-hexanediol (Hex), as in Fig. 3f. **c**, We averaged the PSD95 signals for both control and Hex-treated synapses (8 PSDs imaged in top views, for each treatment). The control shows a somewhat regular pattern, while the Hex treatment seems to perturb this.

**Extended Data Fig. 7.**
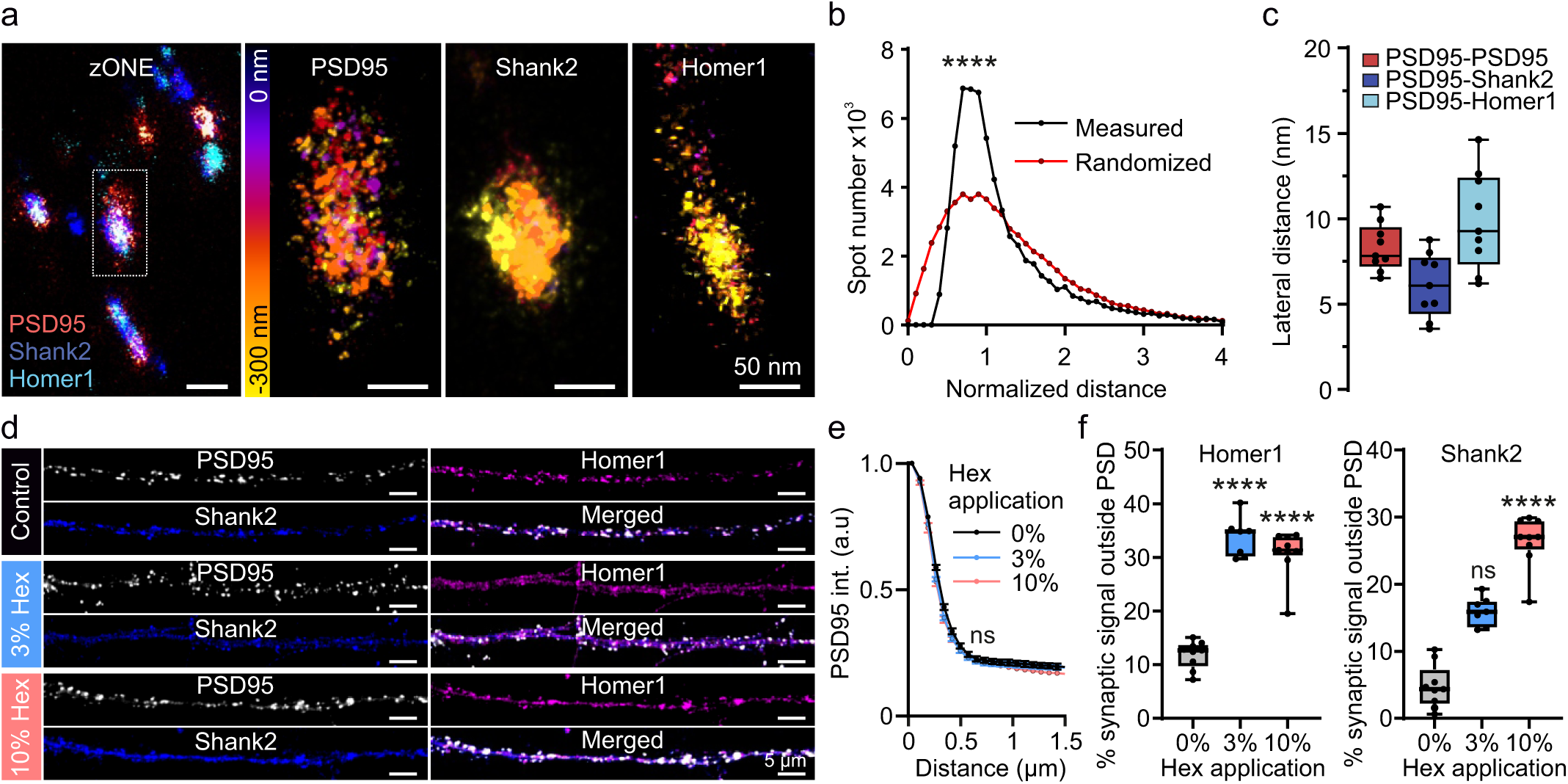
A detailed analysis of the PSD. **a**, The PSD was immunostained for PSD95, Homer1 and Shank2, as in Fig. 3, and images were taken at different heights along the Z-axis (zONE imaging). An overlay (summed image) is shown in the left panel, along with an analysis of the proteins at different Z levels, using a colormap that describes the positions along the Z axis (right panel). **b**, The distance between PSD95 spots was computed from images as in panel a, and was compared to that obtained from positioning the molecules randomly within the PSD95, N = 10 synapses, Friedman test followed by Dunn-Sidak, *p*=0.0001. **c**, The lateral distance between PSD95 spots and between PSD95 and Homer1 or Shank2. The minimal distance between each PSD95 spot and a Homer1/Shank2 spot is shown (measured in the lateral plane, in 2D projections of the PSD). N = 10 synapses, from 2 independent experiments. While the distance between PSD95 spots has a non-random character, as indicated in panel b, the distances to Homer1 or Shank2 spots are not different from randomized distributions (Dunn-Sidak tests, p>0.1), possibly also because these two molecules are immunostained using antibodies, which causes the fluorescence signals to scatter broadly. **d**, Confocal microscopy analysis of the PSDs, in non-expanded samples. In control conditions all three components analyzed here (PSD95, Homer1, Shank2) are well colocalized. The addition of 3% 1,6-hexanediol (Hex) causes the dispersion of Homer1 (magenta), while 10% Hex also disperses Shank2 (blue). PSD95 remains largely unaffected by Hex. **e**, An analysis of the average PSD95 spot profile confirms this impression, N = 10-7-10 neurons, a set from 3 independent experiments. **f**, We analyzed the dispersion of Homer1 (left) and Shank2 (right) away from the PSD95 spots. The signal present in synapses (near the PSD95 labeling, but not within the PSD) was analyzed, to determine the % that is not correlating to the PSD structure. The same samples were analyzed as in panel e.

**Extended Data Fig. 8.**
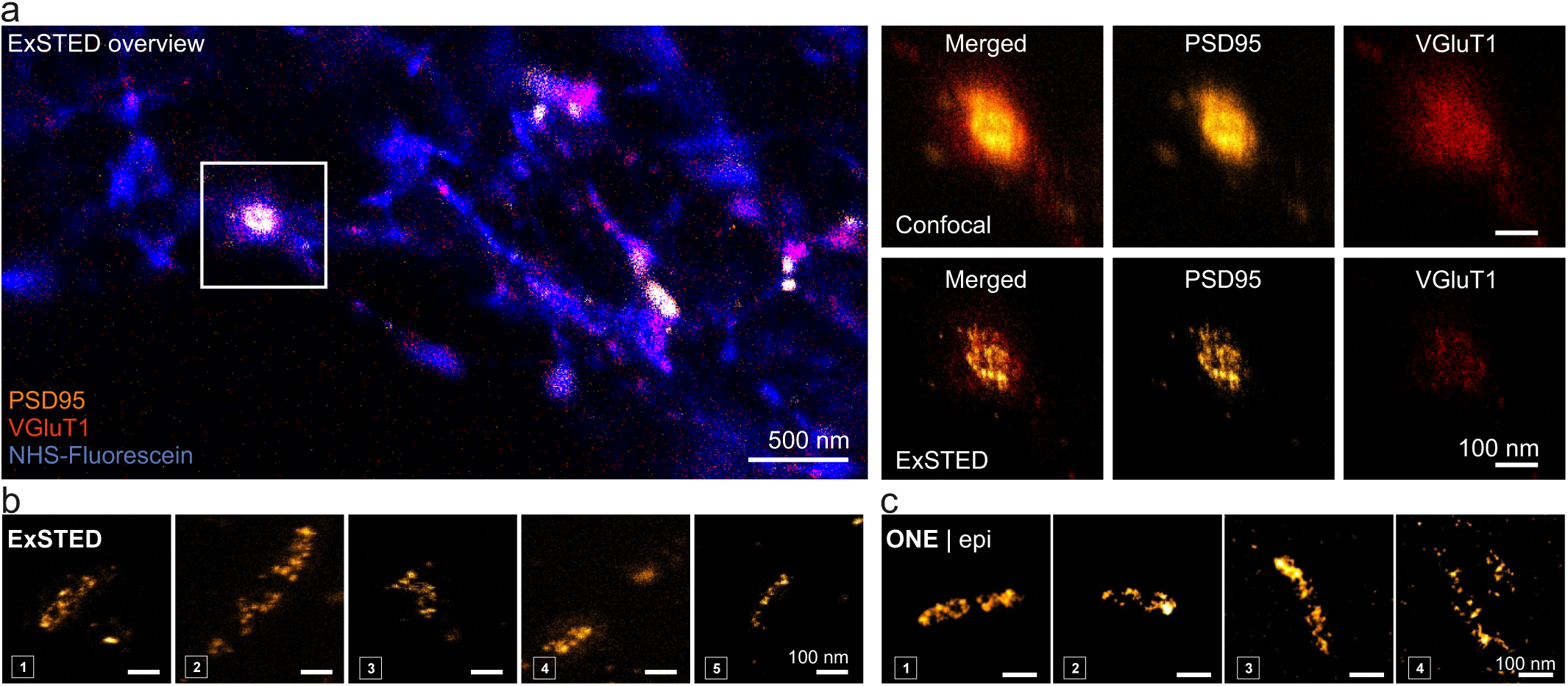
ExM-STED (ExSTED) imaging of PSDs. **a**, Hippocampal cultures were immunostained for PSD95 and VGlut1, and were additionally labelled with NHS-ester fluorescein, after homogenization. **b**, A gallery of high-zoom ExM-STED views of synapses, with a focus on PSD95. Relatively large PSD domains are visible, as in most previous works in the literature, and unlike most of our ONE images. **c**, To determine if this is simply an issue of resolution, we aimed to generate ExM-STED-like images with ONE microscopy, by reducing its resolution. We employed an epifluorescence microscope (as opposed to a rapidly scanning confocal in the panels dealing with PSD95 in Fig. 5), and we used the temporal radiality pairwise product mean (TRPPM) option of analysis, which broadens the resulting spots. The results are very similar to ExM-STED images, demonstrating that the modular/domain appearance of the PSD95 stainings is a result of insufficient resolution, with a *retiolum* being evident only at very high resolution (under optimal ONE imaging).

**Extended Data Fig. 9.**
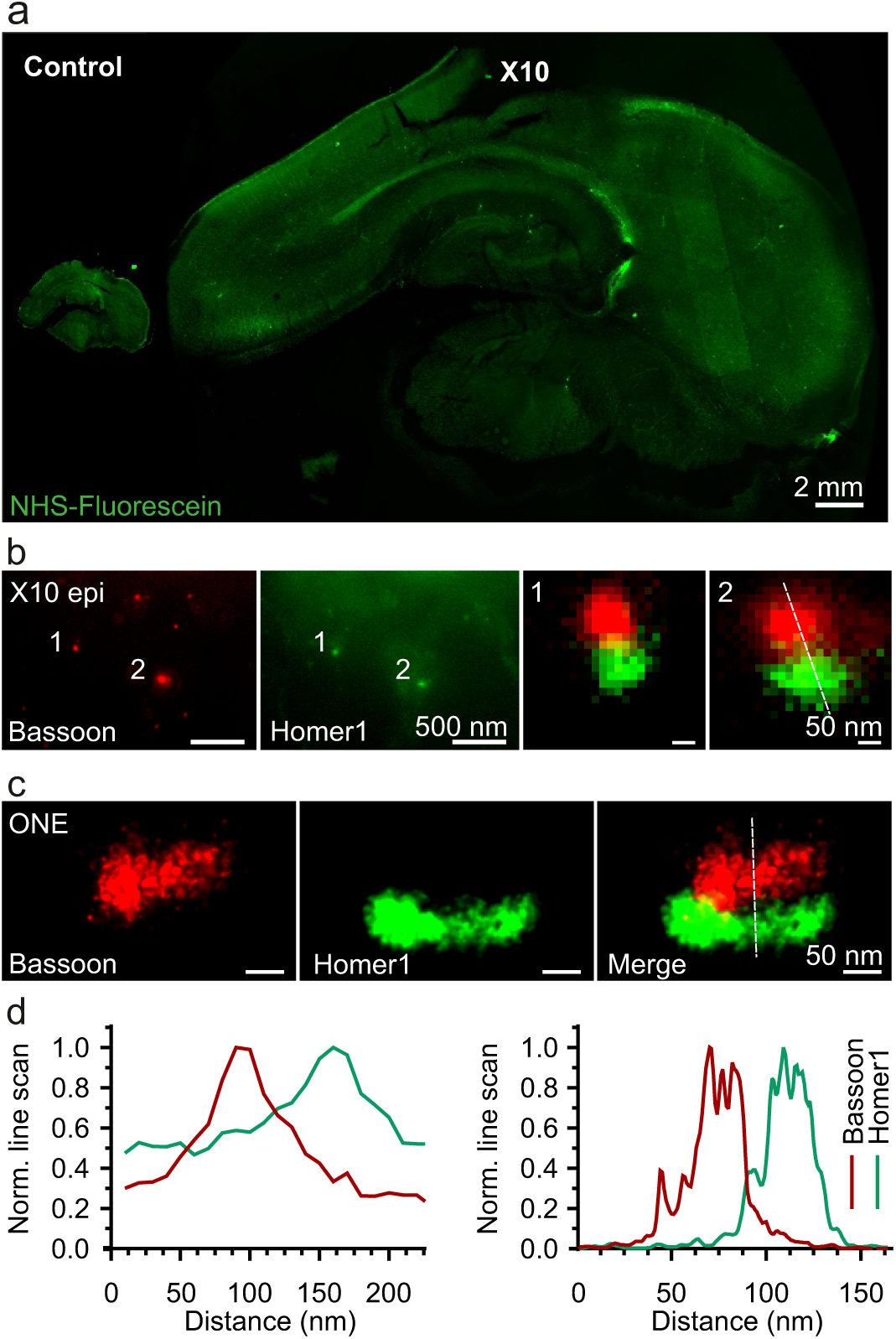
ONE analysis of brain slices. **a**, Images of a 200 µm-thick rat brain section before (left) and after (right) expansion, relying on autoclaving for homogenization^47^. The scale bar does not take the expansion factor into consideration. The sections were labelled by using NHS-ester fluorescein incubations. **b**, Epifluorescence images of expanded brain slices, focusing on Bassoon and Homer1 as pre-and postsynaptic markers, respectively. **c**, Similar images, taken using the ONE procedure. **d**, Line scans executed over the areas indicated in panels b and c. As expected, far more details can be observed in ONE than in simple epifluorescence microscopy.

**Extended Data Fig. 10.**
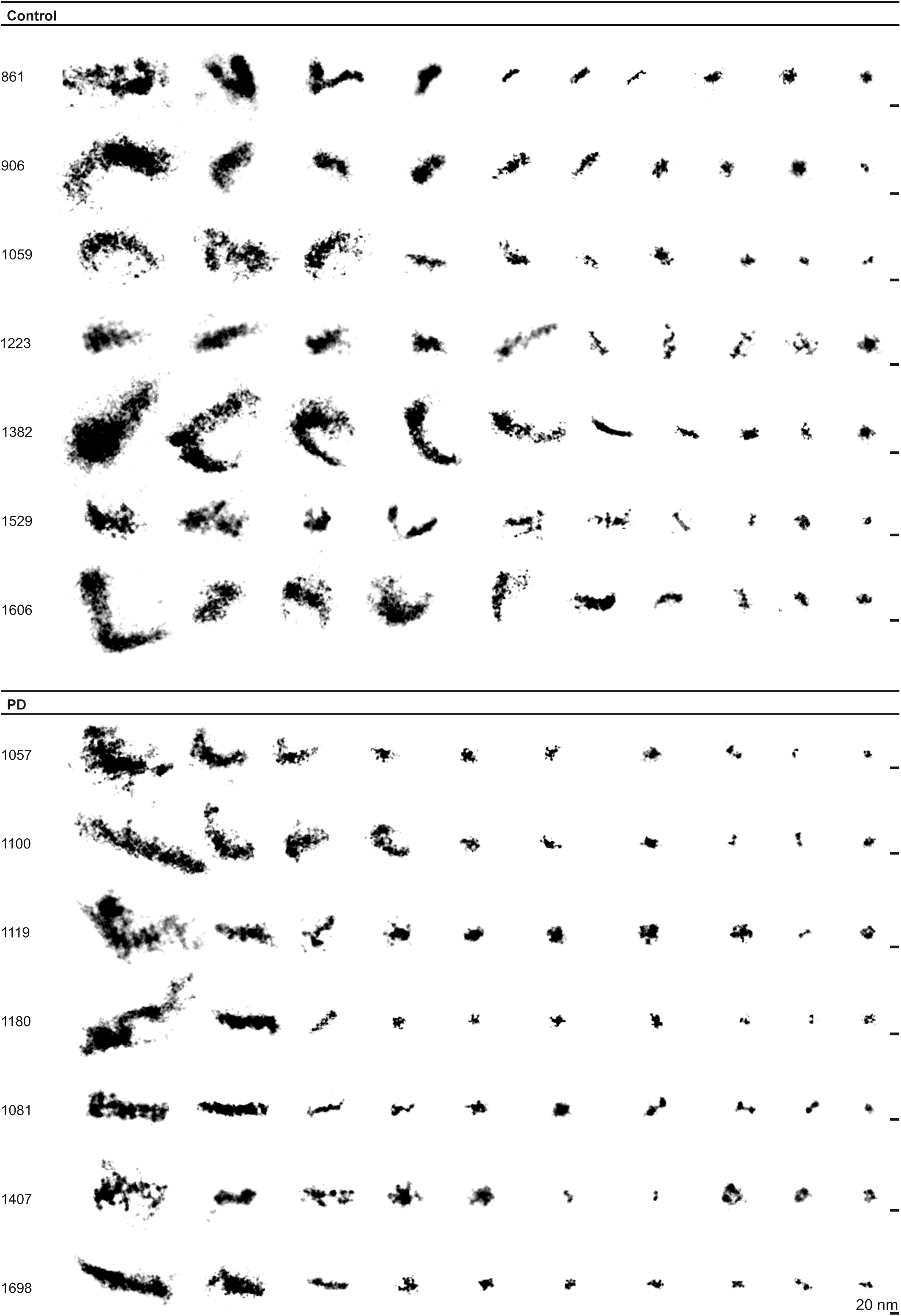
A gallery of ASYN object images from 7 PD patients and 7 controls. The images were obtained following the procedure indicated in Fig. 4a. See Supp. Table 1 for details on the respective patients.

## Supplementary Figure legends

**Supplementary Fig. 1.**
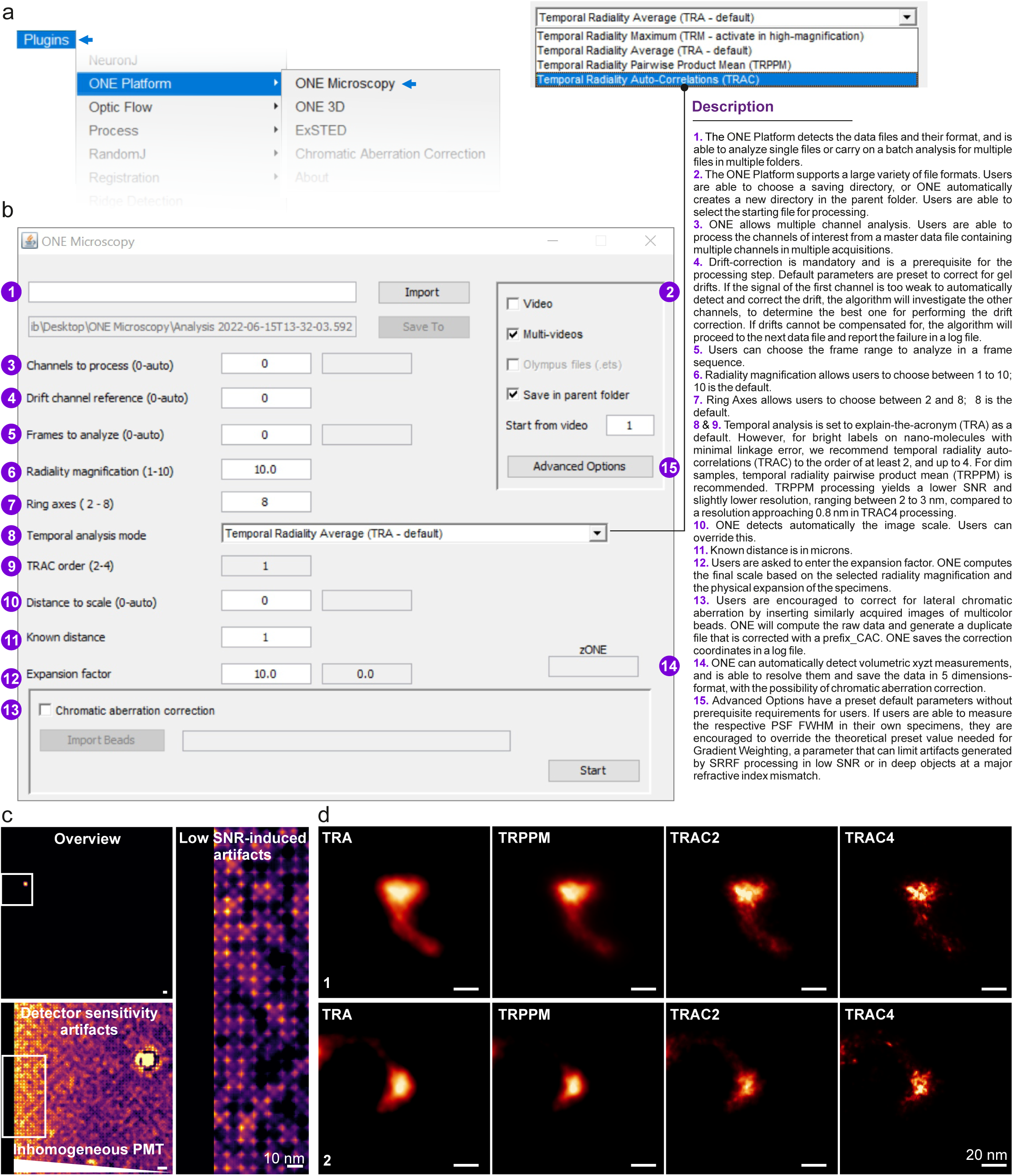
ONE analysis and examples. **a** & **b**, Several views of the starting interface of the ONE software package. The examples show the intuitive software choices. See also the “Readme/Help” file of the software package. **c**, Examples of different potential artifacts that should be avoided in ONE imaging. **d**, Different potential choices in how to resolve ONE images. We suggest using the temporal radiality pairwise product mean (TRPPM) procedure for dim samples. This reduces the obtainable resolution, but follows much better the potential sample shape. For brightly labelled samples with direct labeling, the temporal radiality auto-cumulant (TRAC4) procedure provides the best resolution and SNR, indicating the positions of the individual fluorophores.

**Supplementary Fig. 2.**
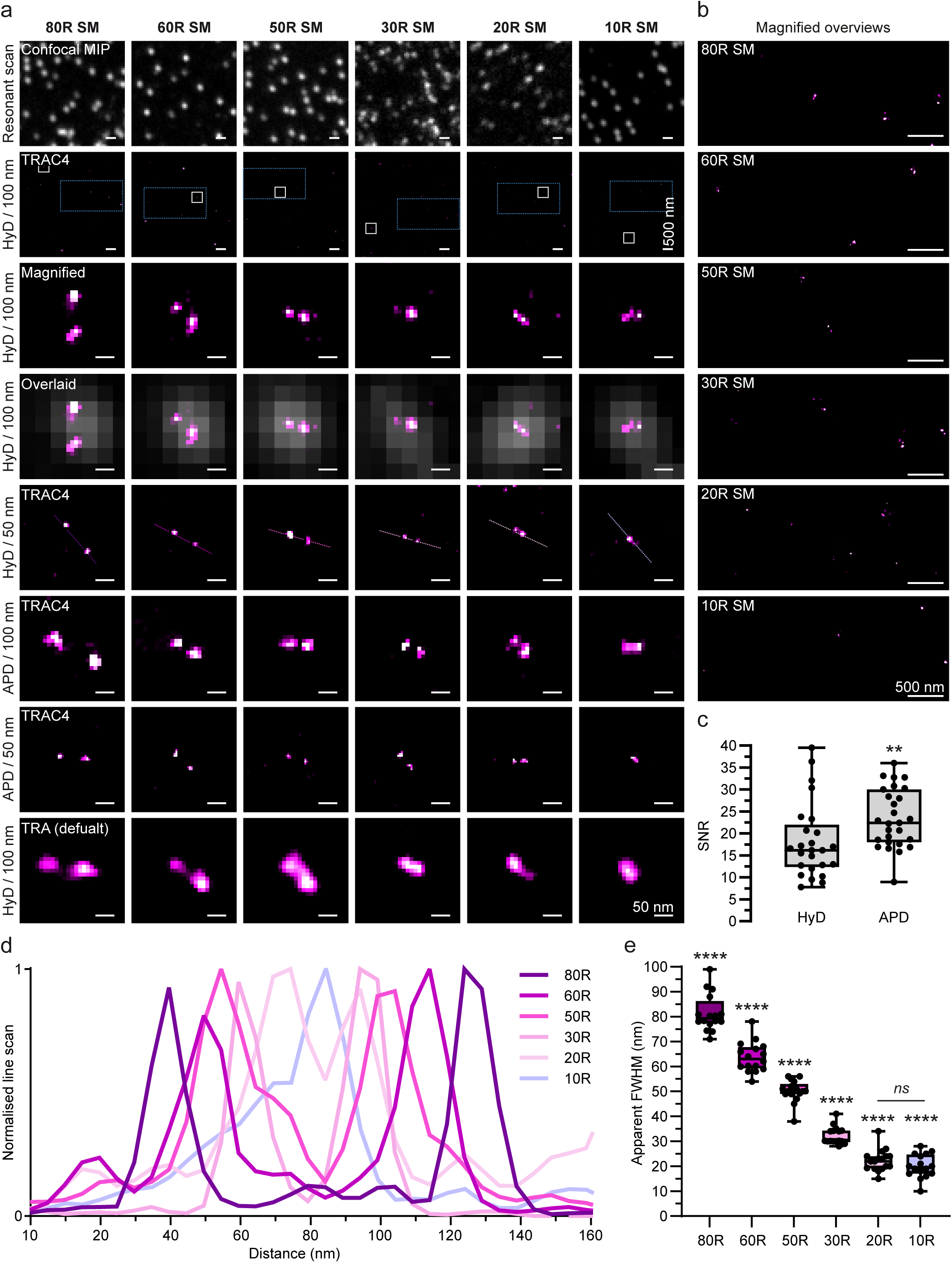
Evaluating SRRF analysis performance using DNA origami nanorulers, in non-expanded samples. **a**, Nanorulers with <u>s</u>ingle Atto647N <u>m</u>olecules (R SM) were generated by GATTAquant^30^ carrying fluorophores on each end of DNA structures of 80, 60, 50, 30, 20 and 10 nm in length. They were then imaged using a confocal resonant scanner, without expansion procedures. The first panel shows confocal maximal intensity projections (MIPs) for each of the rulers. The second panel shows temporal radiality averaging (TRA) analysis overviews. White boxes indicate the magnified regions displayed in the third panel. The fourth panel shows a temporal radiality auto-correlations of fourth order (TRAC4) analysis, overlaid with the respective confocal MIPs. The remaining panels show different ruler examples, acquired at different starting pixel sizes, using either a hybrid detector (HyD) or an avalanche photodiode detector (APD), and analyzed in different SRRF modalities. This analysis is shown in the fifth panel for 50 nm pixel size, using a HyD and analyzed using TRAC4. The sixth and seventh panels show rulers acquired at 100 and 50 nm pixel sizes, using an APD and analyzed using TRAC4. The eighth panel shows rulers that were acquired at 100 nm pixel size and were analyzed using default SRRF settings (TRA). **b**, Magnified overviews of selected regions (indicated by blue rectangles) from each of the ruler exemplary images to the left. **c**, Signal-to-noise ratio (SNR) analysis of HyD and APD detectors, N = 25 and 30 for HyD and APD, respectively. Mann Whitney test, *p* = 0.004. **d**, Normalized line scans across the different ruler images, as indicated in the respective panels in (**a**). **e**, Apparent FWHM of the different rulers. N = 17, 17, 18, 17, 18 and 17 for 80, 60, 50, 30, 20, and 10 R SM, respectively. A Kruskal-Wallis test was applied, followed by Dunn’s *post hoc* test; *p* < 0.0001.

**Supplementary Fig. 3.**
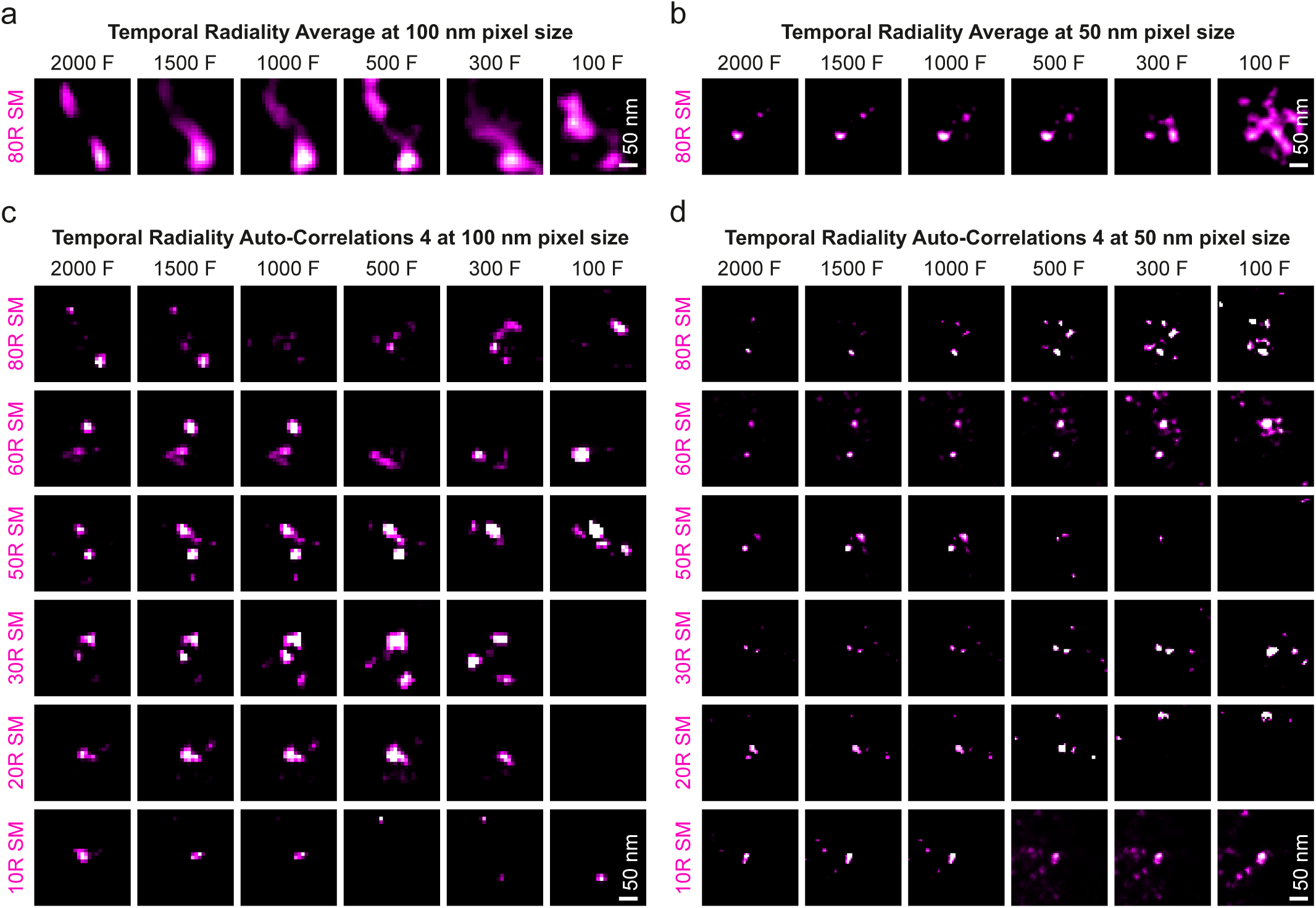
The effect of frame number on SRRF analysis. **a** & **b**, 80 nm rulers were imaged at 100 and 50 nm pixel sizes, and were then analyzed with the default SRRF parameter (temporal radiality average, TRA), using varying frame counts (termed F in the figure), from 100 to 2000. **c** & **d**, The same procedure was repeated using temporal radiality auto-correlations (TRAC4) for rulers of 10 to 80 nm. The frame count does not affect the TRA analysis as much as it affects TRAC4. The TRA performance, which is the parameter reported in most publications, is far poorer than that of TRAC4, when sufficient frames are analyzed.

**Supplementary Fig. 4.**
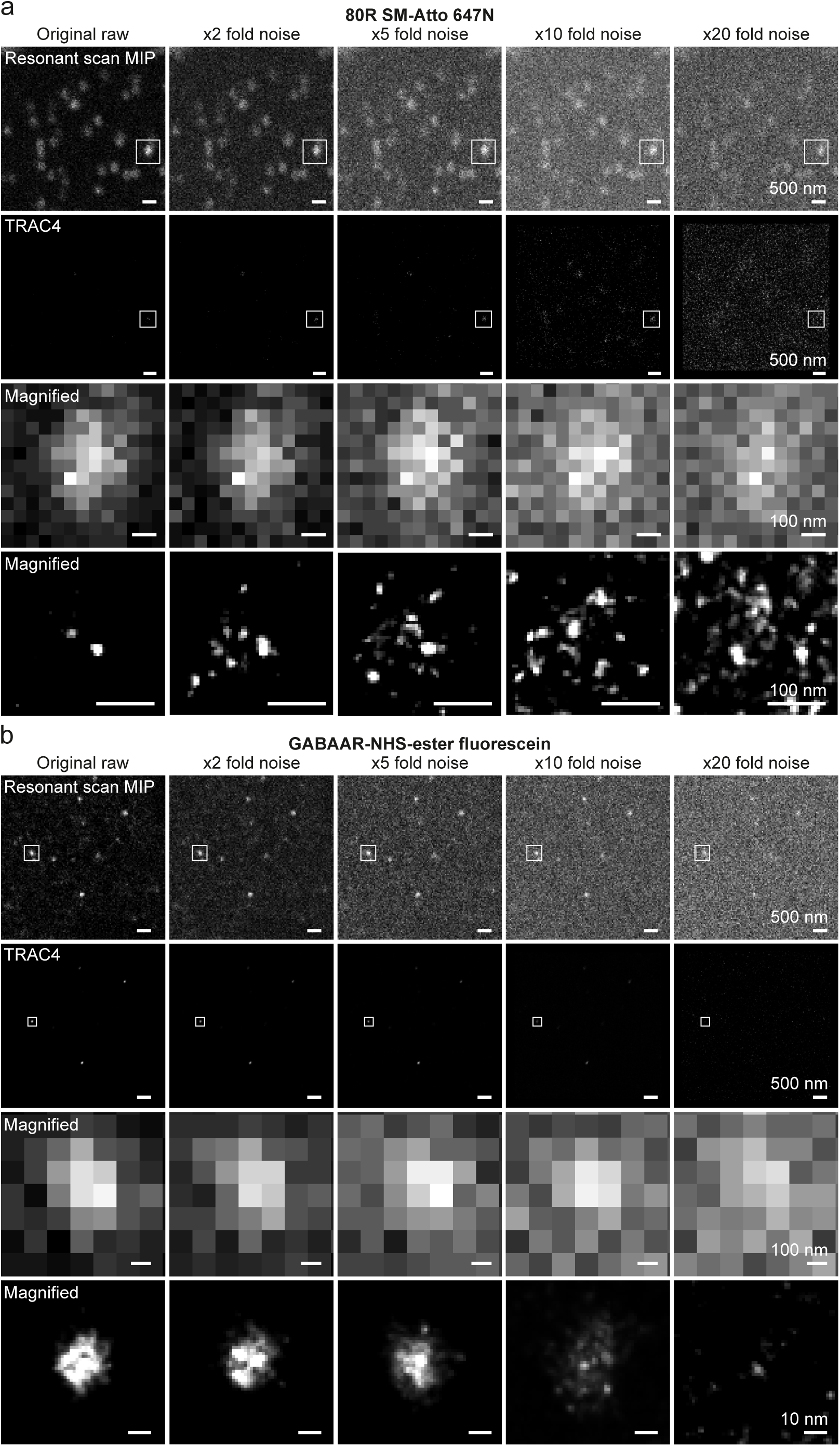
SNR effect on SRRF performance. **a**, The top panel shows an overview of 30 frame-MIPs of an 80 nm ruler, followed by MIPs of the same ruler that were subjected to 2-fold, 5-fold, 10-fold and 20-fold increase in noise. Noise was added artificially, using a Matlab routine. The initial SNR was 27.84 .The second panel shows TRAC4 analyses of the data. The third and fourth panels show a magnified region from the resonant scan MIPs, and their respective TRAC4 analysis results. **b**, The same analysis was performed on expanded GABA_A_R. Note that the receptor pore disappearing at x5 fold noise in TRAC4 resolved image. The nanoruler image is corrupted far more strongly by a 2-fold increase in noise than that of the GABA_A_R, owing to the substantially higher original SNR of the receptor image (76.72).

**Supplementary Fig. 5.**
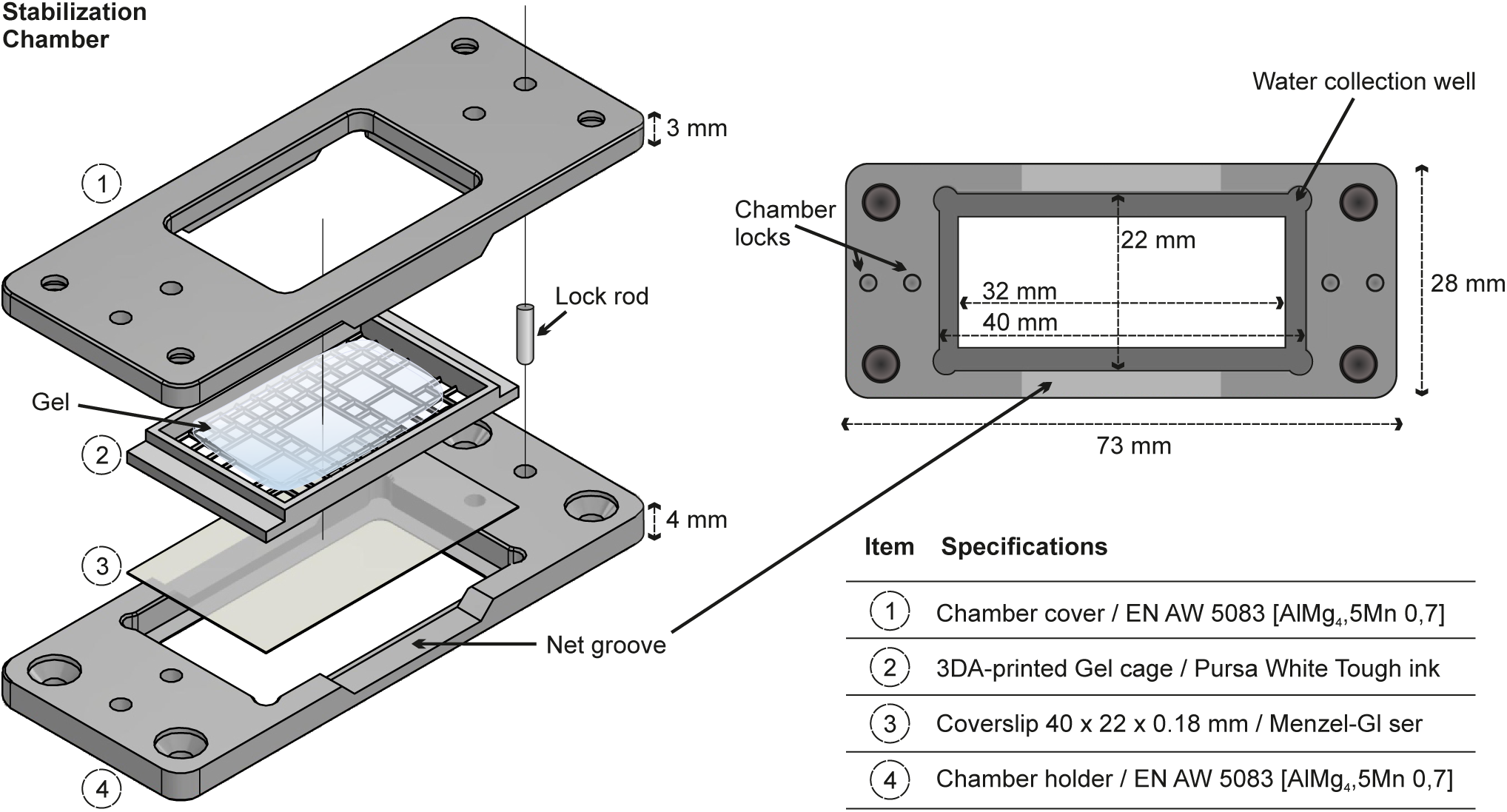
Technical scheme of the stabilization chamber used in this work. The exact measurements and materials for the stabilization chamber are included in the figure text. The 3D-printed gel cage patterning can be organized according to the user’s preferred design. Only a suggested design is included here (many others work equally well). The exact design files can be obtained from the corresponding authors, to produce this chamber in any facility.

**Supplementary Fig. 6.**
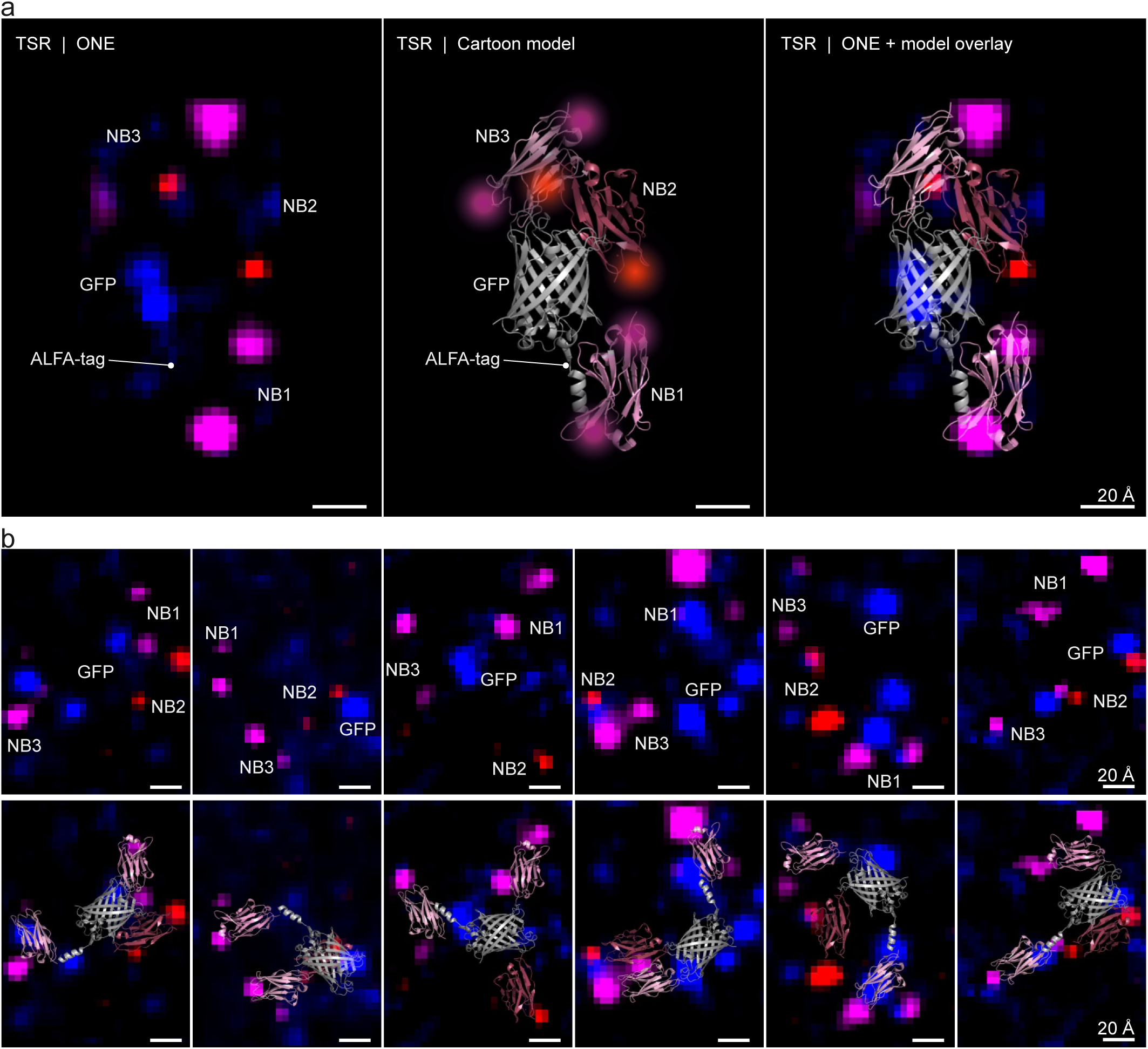
TSR gallery. **a**, An example of a TSR. The first panel shows a ONE image of a TSR, the middle panel shows a cartoon model that fits the imaged TSR, and the third panel shows an overlay of the ONE image and the model. **b**, A gallery of TSRs (upper panels) and a best guess of cartoon models overlaid over the TSR images (lower panels).

**Supplementary Fig. 7.**
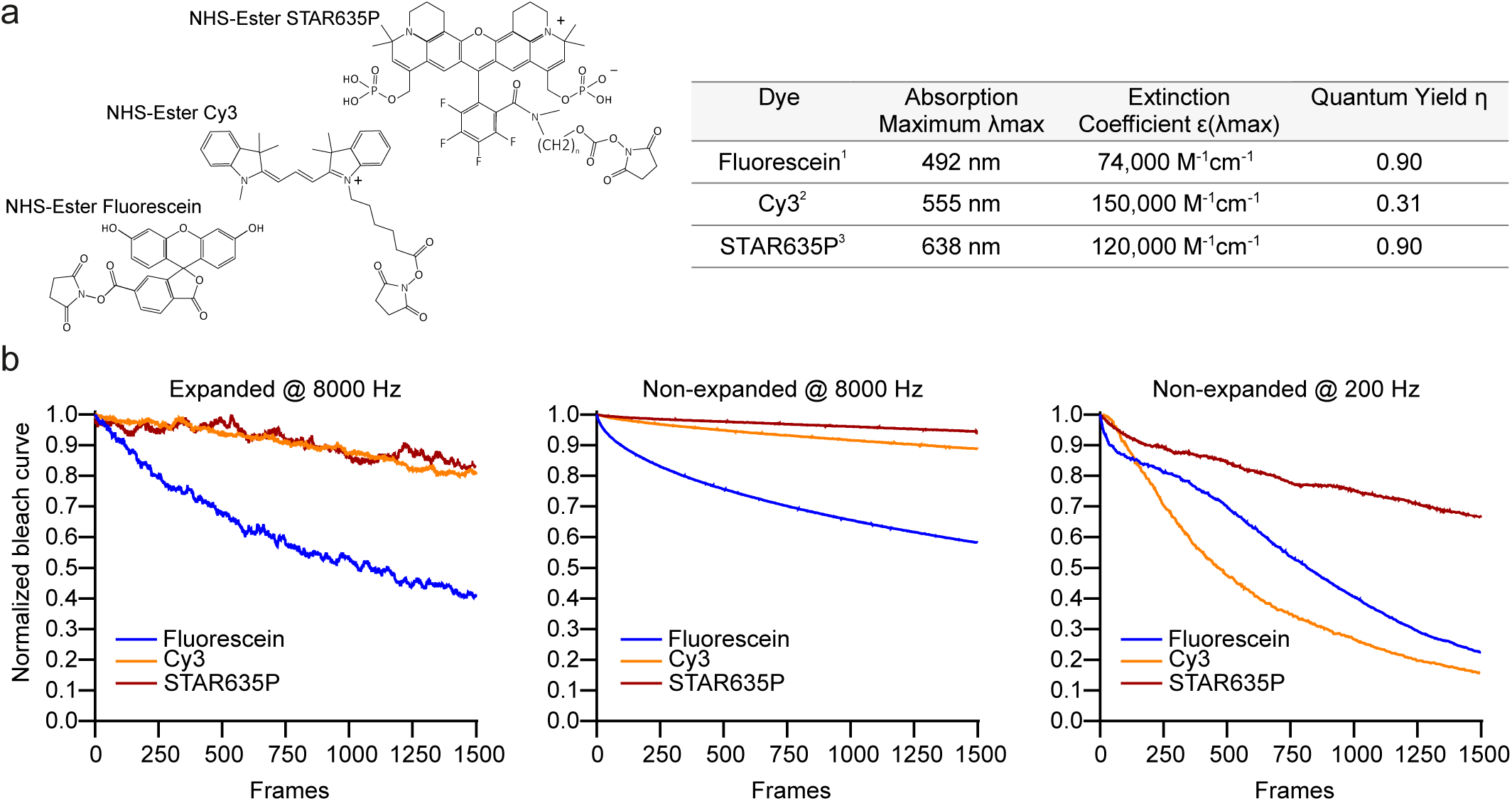
Bleaching properties of fluorescein, Cy3, and STAR635P. **a**, A representation of the structures of each of the used dyes, followed by a table of their properties. The molecule structures and properties were reproduced from measurements of commercial providers: ^1^https://broadpharm.com/product/bp-23900, ^2^https://broadpharm.com/product/bp-22535, and ^3^https://abberior.shop/abberior-STAR-635P. **b**, Normalized bleach curves from expanded specimens at 8000 Hz, and non-expanded specimens at 8000 Hz and 200 Hz.

**Supplementary Fig. 8.**
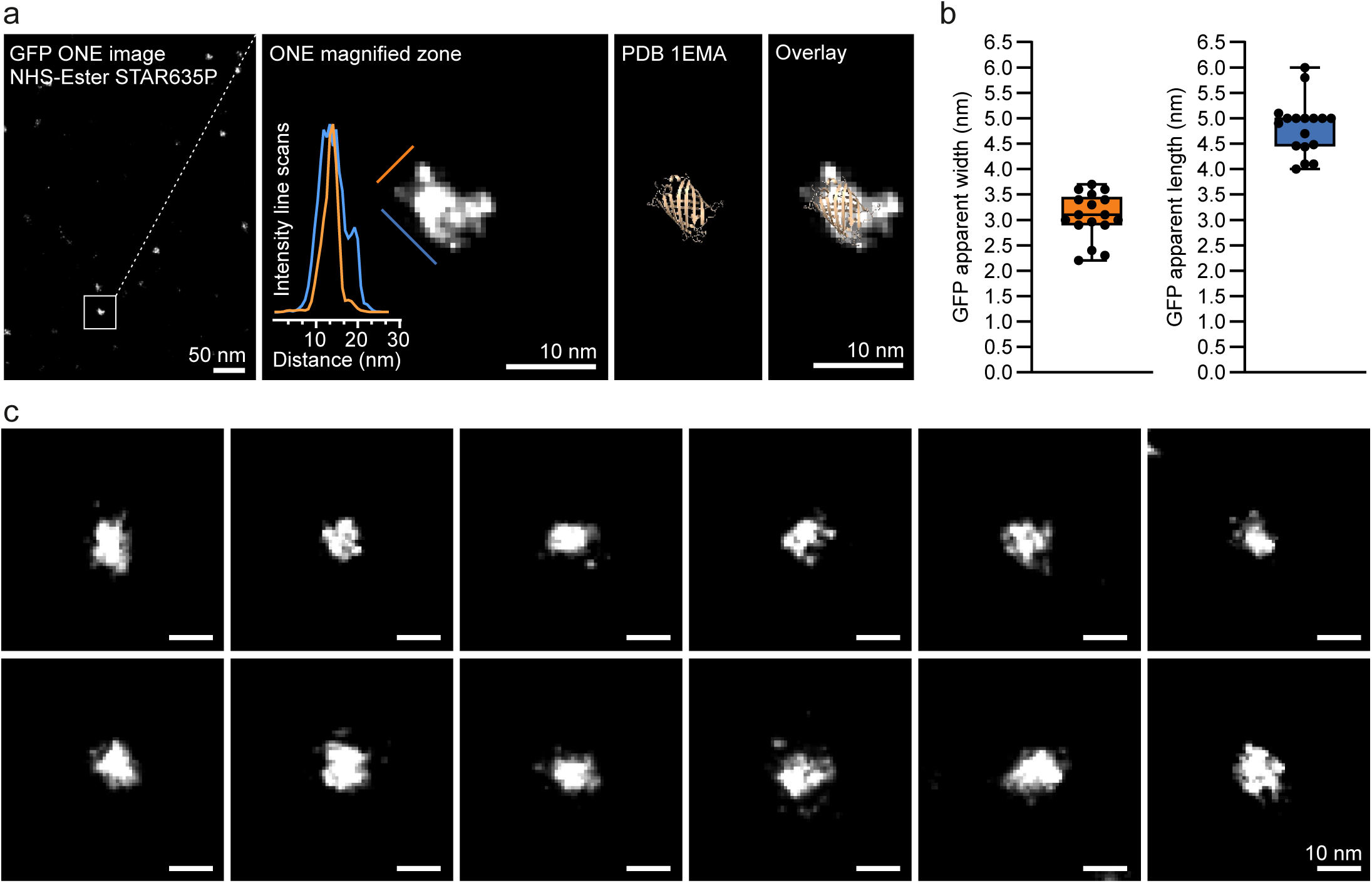
ONE imaging of purified eGFP molecules. **a**, The first panel shows a ONE overview of eGFP molecules labelled with NHS-Ester STAR635P. The second panel shows a magnified area. The third panel shows the eGFP 1EMA PDB structure. The fourth panel shows a PDB/fluorescence overlay. **b**, A measurement of the apparent width and length of the molecules, from line scans as the examples shown in panel a, in blue and orange. A total of 17 single molecules were measured. **c**, A gallery of eGFP molecules.

**Supplementary Fig. 9.**
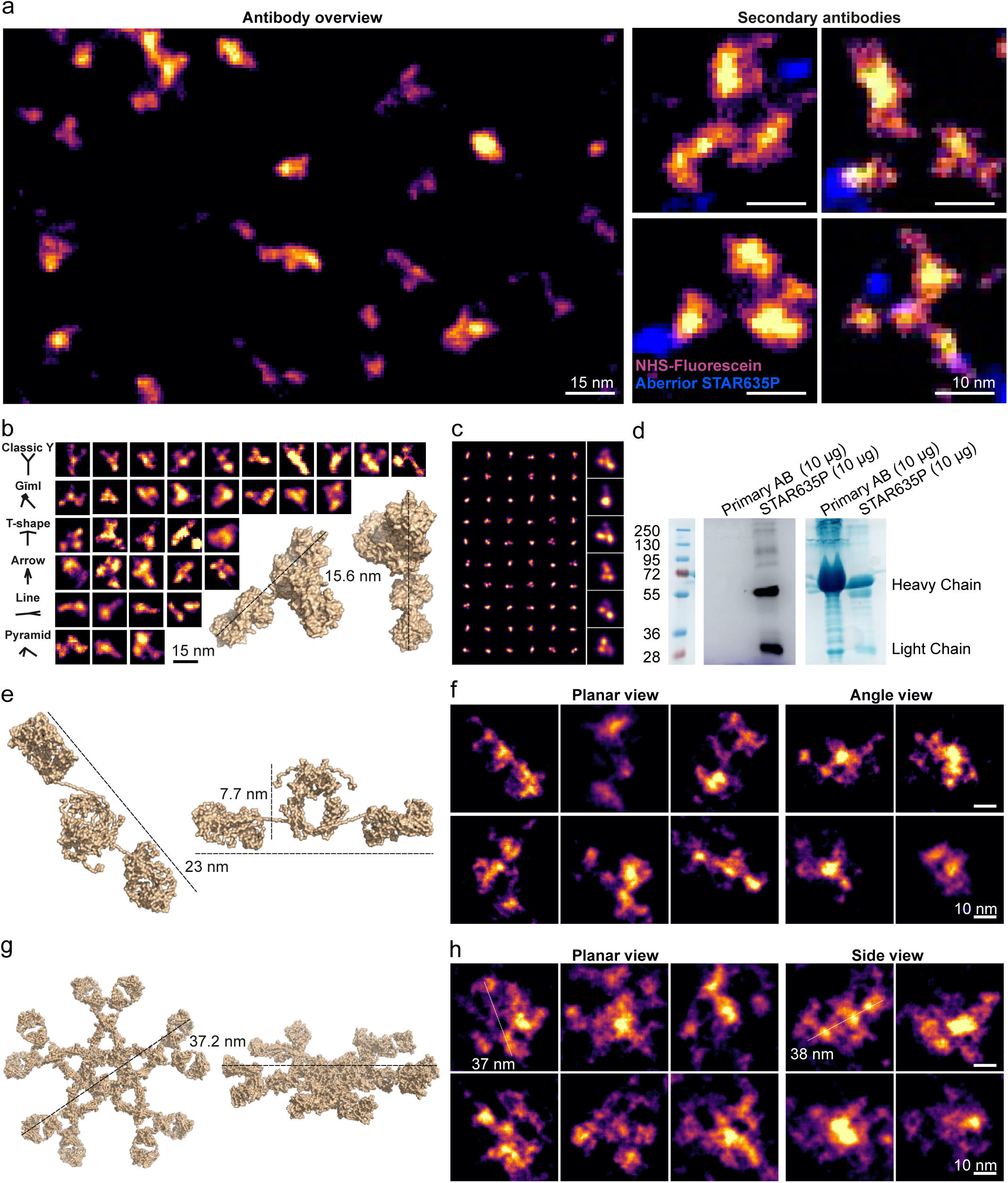
Further ONE examples of immunoglobulin imaging. **a**, An overview of a field showing IgG antibodies labelled using NHS-fluorescein (left), along with a few zoom-in images of fluorescently-conjugated secondary IgG antibodies (right; Abberior Star635P conjugation shown in blue). **b**, Several examples of IgG antibodies imaged in different positions and perspectives. **c**, A gallery of the expected antibody shapes, obtained by convoluting a PDB IgG structure with a ONE point-spread-function, after revolving the IgG molecules in 3D space randomly. A few enlarged views are shown, along with a multitude of small-sized views, to explain how IgG molecules should appear when they are visualized in fluorescence in random orientations. The typical IgG views are similar to the modeled ones. **d**, Fluorescence (Abberior Star635P) and Coomassie SDS-PAGE gels indicating the size distribution of antibody fragments. A mouse monoclonal primary antibody was run on the gels, along the secondary antibody imaged in panel a. The gel was first imaged under a fluorescence (Cy5 channel) and then total proteins were revealed with Coomassie brilliant blue staining. The results suggest that numerous small fragments are expected for both primary and secondary antibodies in the ONE images, not only full antibodies, due to impurities being present in the commercial antibody samples. **e-f**, An overview of IgA molecules. **g-h**, A similar overview of IgM molecules. The antibody structures are shown using Pymol representations from PDB structures 1HZH, 1IGA, and 2RCJ.

**Supplementary Fig. 10.**
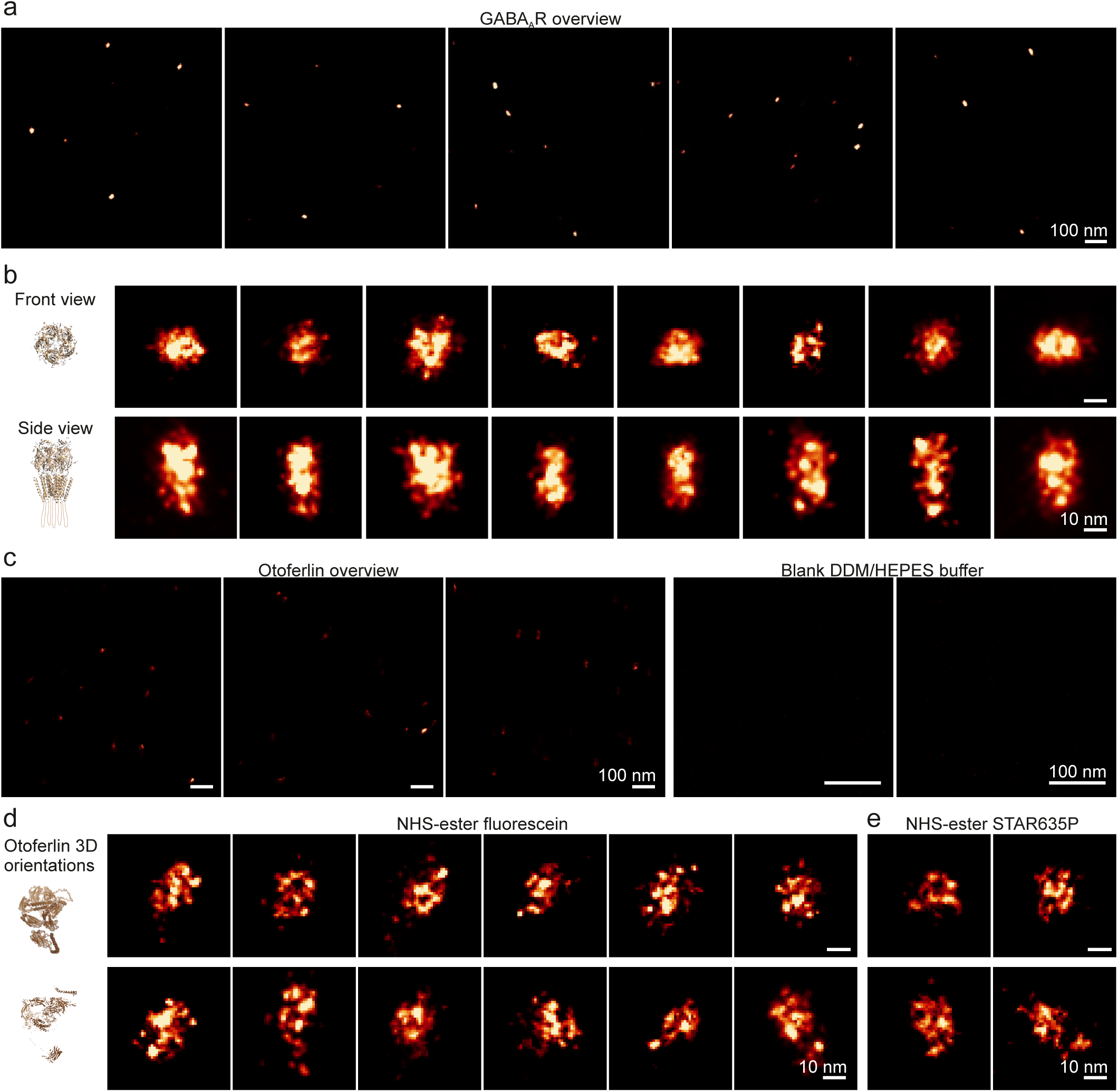
GABA_A_ receptor and otoferlin galleries. **a**, An overview of images of GABA_A_ receptors. **b**, The images display GABA_A_ receptors in different 3D positions. The positional indications are best guesses performed by an experienced investigator. **c**, Overview images of otoferlin (right panel), and blank buffer as a control (left panel). **d**, Otoferlin labelled with NHS-ester fluorescein ONE images in different 3D positions. **e**, Otoferlin labelled with NHS-ester STAR635P ONE images.

**Supplementary Fig. 11.**
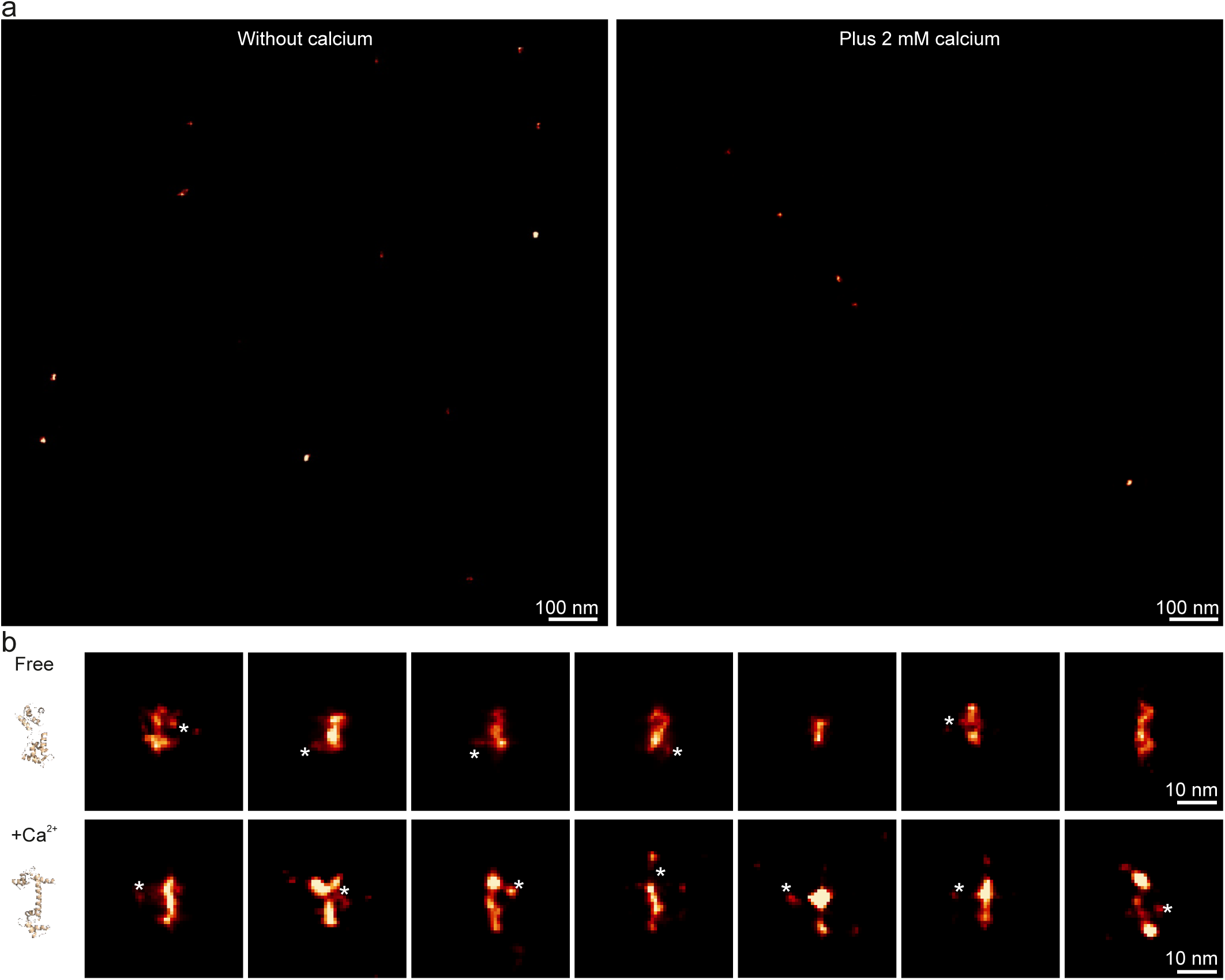
Calmodulin gallery. **a**, An overview of calmodulin ONE acquisitions in the presence and absence of calcium. This molecule was expressed and purified as a chimera containing mEGFP. The compact signal associated to the GFP molecule, as observed already in the TSR images in Fig. 1, has a limited contribution to the overall size of the molecule. **b**, Exemplary zoomed calmodulin ONE images. The asterisk denotes the best guess of GFP molecule bound to calmodulin.

**Supplementary Fig. 12.**
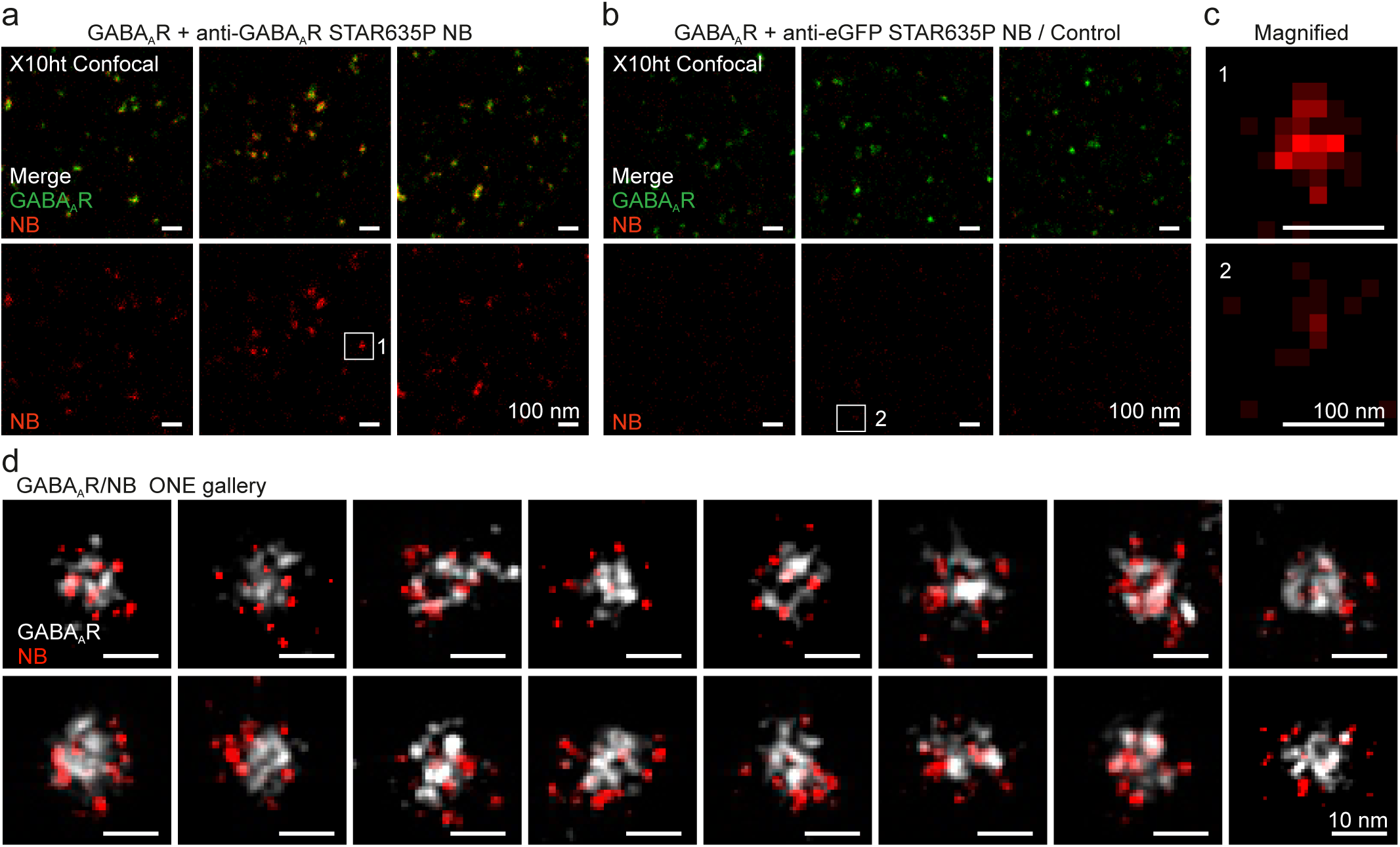
GABA_A_R nanobody labeling. **a**, Confocal images of expanded GABA_A_R labelled with anti-GABA_A_R nanobodies (NBs) conjugated to STAR635P. **b**, Confocal images of expanded GABA_A_R mixed with anti-eGFP nanobodies, which only induce little non-specific background. **c**, Magnified regions of single receptor either labelled with anti-GABA_A_R or anti-eGFP NBs. **d**, A gallery of ONE images showing GABA_A_R in white and anti-GABA_A_R NBs in red.

**Supplementary Fig. 13.**
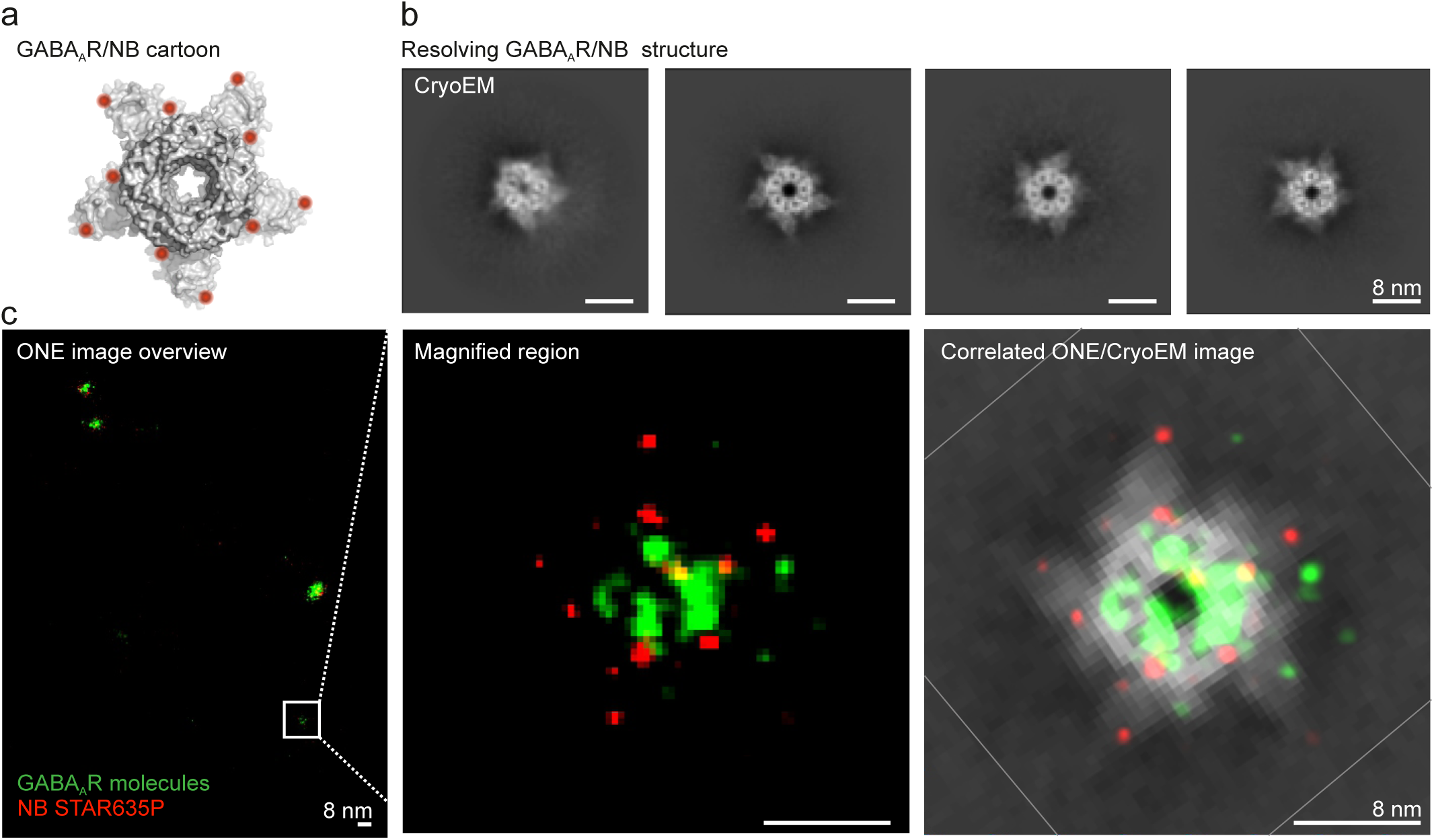
Super-imposition of ONE microscopy images and Cryo-EM data. **a**, A cartoon view of the 5OJM GABA_A_R/NB PDB. The red dots represent the 2 fluorophores on each nanobody. **b**, Cryo-EM images of representative 2D-classes of the GABA_A_R/NB complexes, derived from the same samples as used for ExM. **c**, The first panel shows a ONE image of GABA_A_R/NB. The second panel shows a magnified region of a single receptor. The third panel shows a Cryo-EM/ONE overlay.

**Supplementary Fig. 14.**
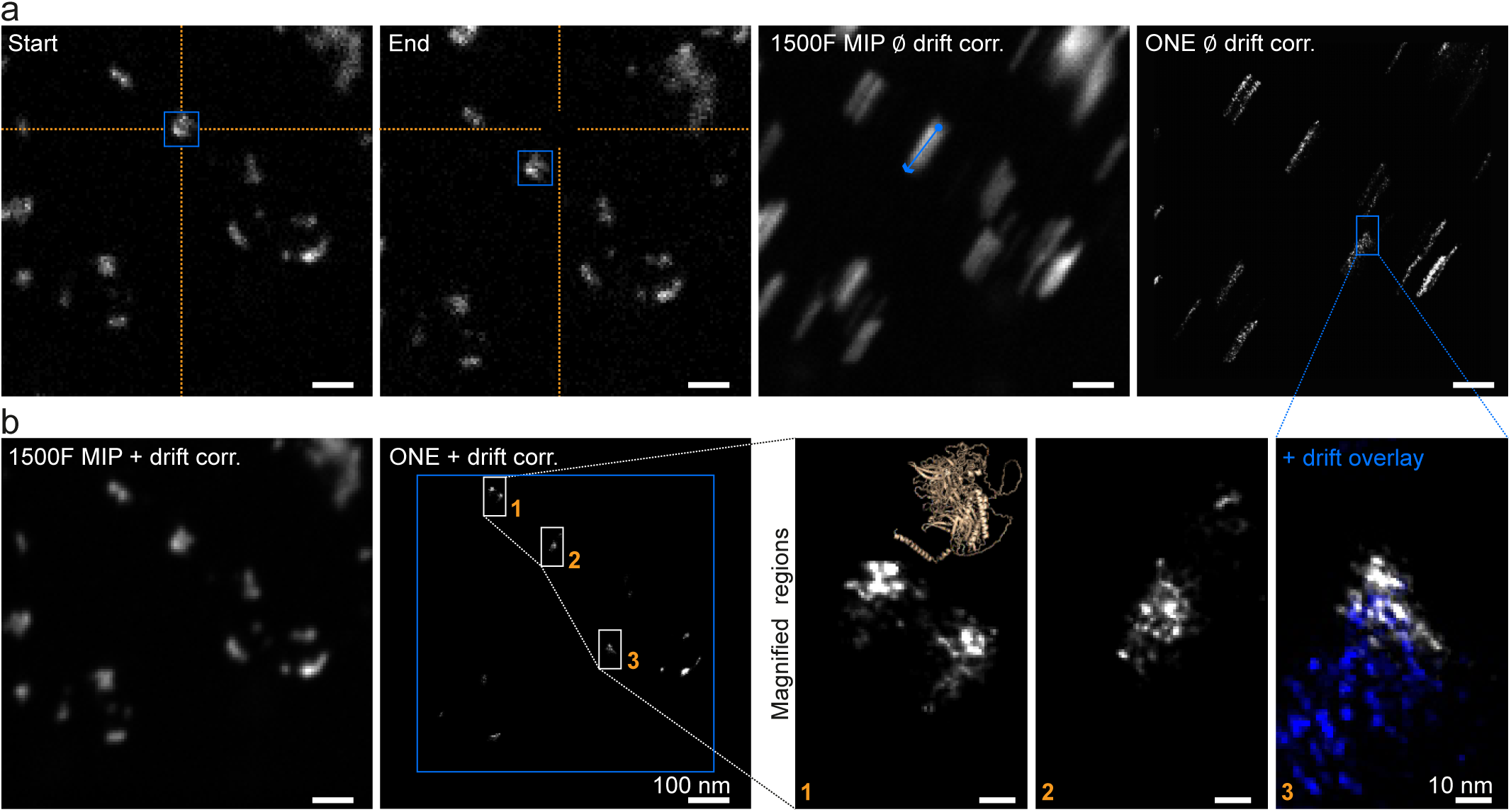
Drift compensation. **a**, A resonant confocal X10 image of otoferlin molecules at the start of a 1500-frame time series recording (first panel) and at its end (second panel). The third panel shows a maximum intensity projection of the resonant confocal scan. The blue arrow indicates the direction of drift. The fourth panel shows ONE processing without drift correction. A streak artefact is evident as a result. **b**, Applying drift correction, using the SRRF software, to the same acquisition and maximum intensity projection yields an image (first panel) similar to the first image in (**a**). The second panel shows the result of the ONE processing with drift correction application. The last set of panels show magnified regions of otoferlin molecules. An otoferlin AlphaFold cartoon is presented for comparison (not drawn to scale). In panel 3, the ONE image is overlaid with its counterpart from the same dataset, processed without drift correction (blue).

**Supplementary Fig. 15.**
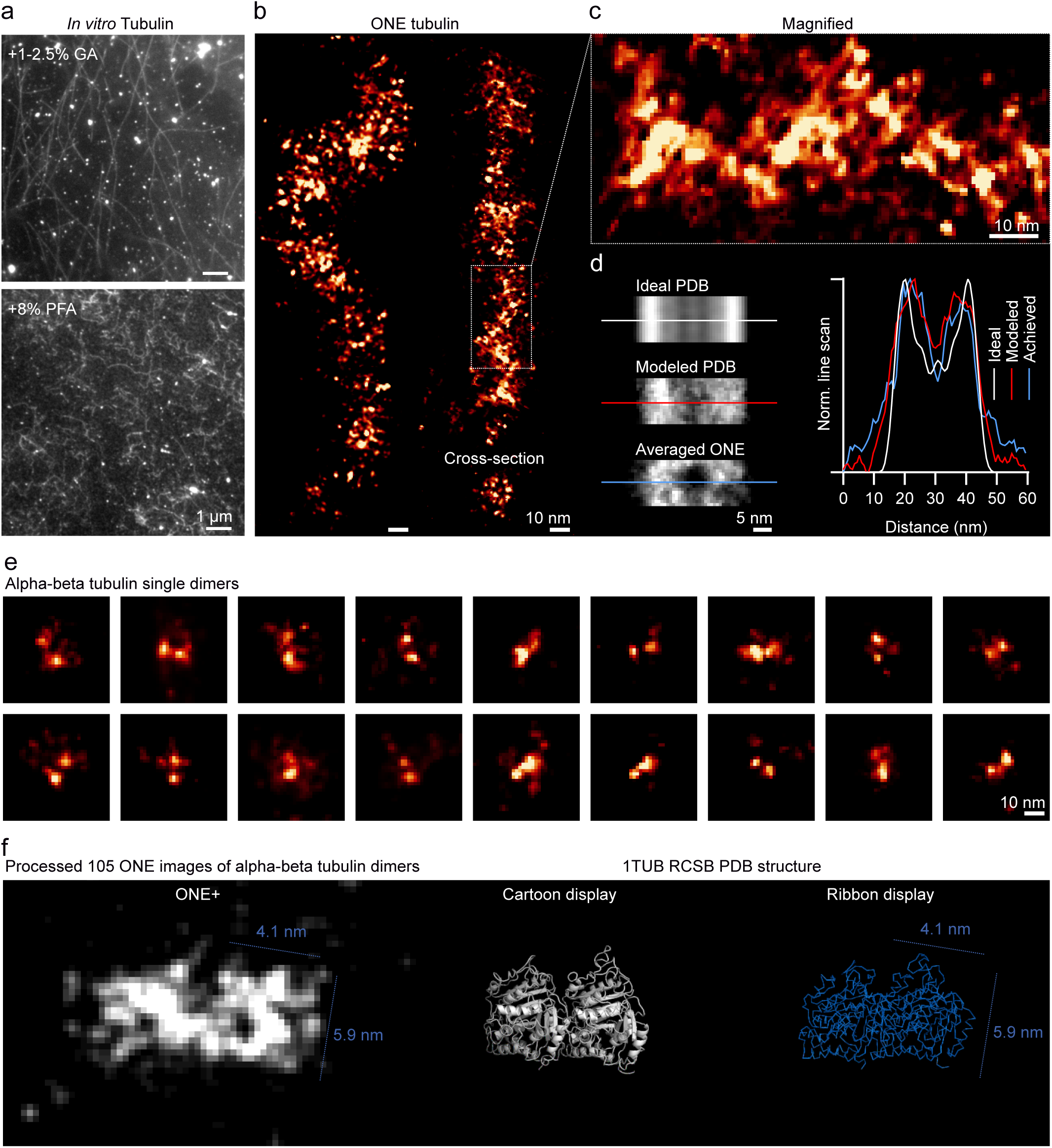
ONE imaging of *in vitro* assembled microtubules. **a**, The upper panel shows *in vitro*-assembled microtubules that are stably fixed with 1-2.5% glutaraldehyde (GA). GA above 0.2% interfered with anchoring into the gels. The lower panel shows less stable microtubules, fixed with 8% PFA. These microtubules are deformed and tend to depolymerize, but this fixation does allow a reasonable degree of anchoring to the gels. **b**, ONE images of microtubules fixed with 8% PFA and expanded and labelled using NHS-ester fluorescein**. c,** A magnified region. **d,** An averaged analysis of side views of microtubule segments. The top panel shows an ideal image, obtained by convoluting the PDB structure of a microtubule segment (3J2U) with a fluorescent PSF, in which every amino acid is labelled fluorescently. The second panel shows a realistic model, in which sparser labelling is considered, and in which different microtubule segments are overlaid with slight tilt angle differences (up to 5°). The third panel shows an averaged processed ONE image of 175 partially depolymerized tubulin segments. The graph shows the respective line scans across each of the panels. **e**, A gallery of alpha-beta tubulin dimers that were left unpolymerized. **f**, The first panel shows a tubulin dimer reconstructed from 105 dimers, following the same procedure as for the GABA_A_Rs, shown in Fig. 3. The second panel shows a 1TUB PDB cartoon structure, and the third panel shows a ribbon display of the molecules, for comparison.

**Supplementary Fig. 16.**
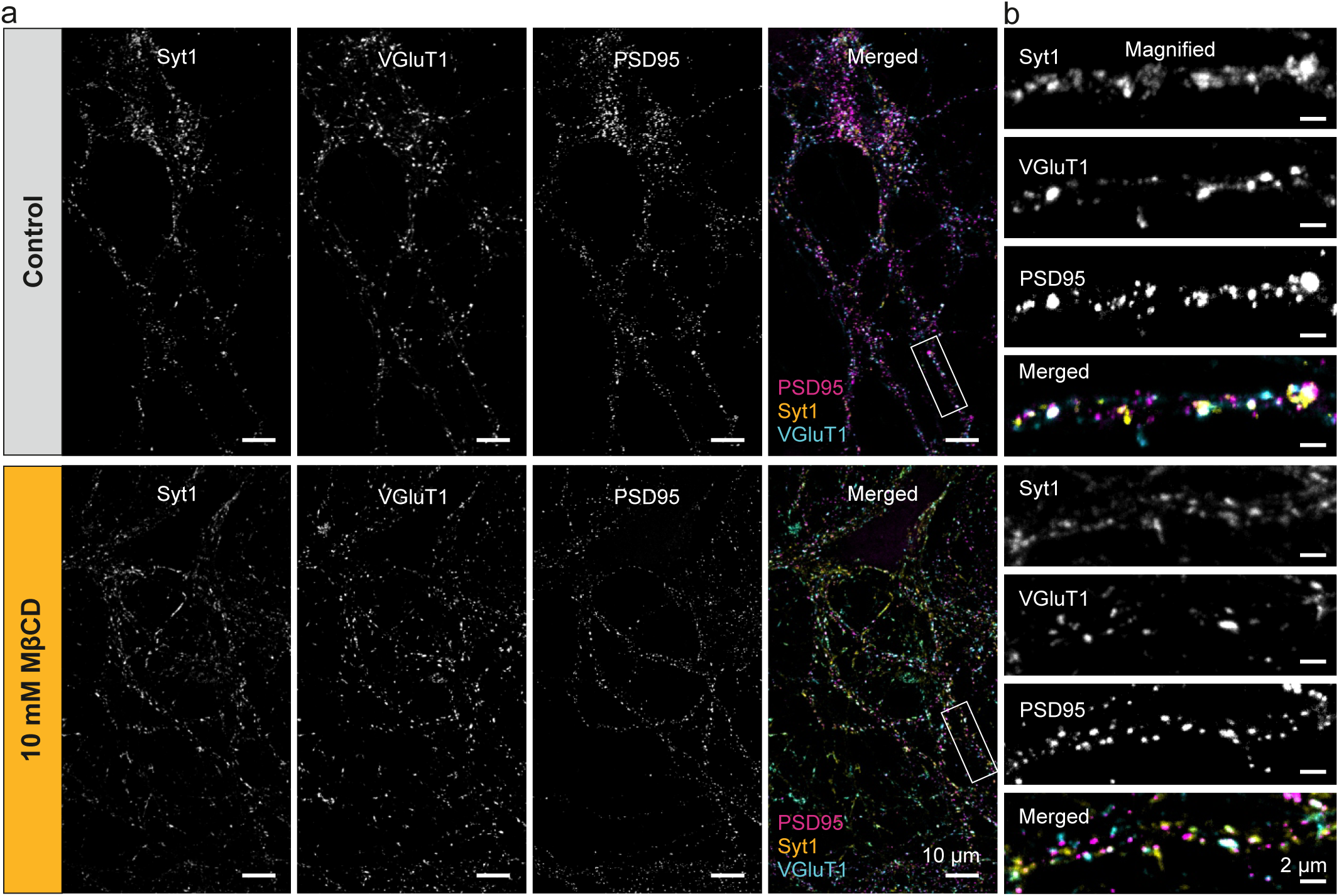
A confocal analysis of synapses after MβCD treatments. **a**, Confocal images of hippocampal cultures immunostained for the three synaptic markers employed in Fig. 3a-c (Syt1, vGlut1 and PSD95), relying on the same staining protocol as in Fig. 3a-c. **b**, The panels show a magnified region. The culture morphology and synapse distribution are similar with or without MβCD treatments.

**Supplementary Fig. 17.**
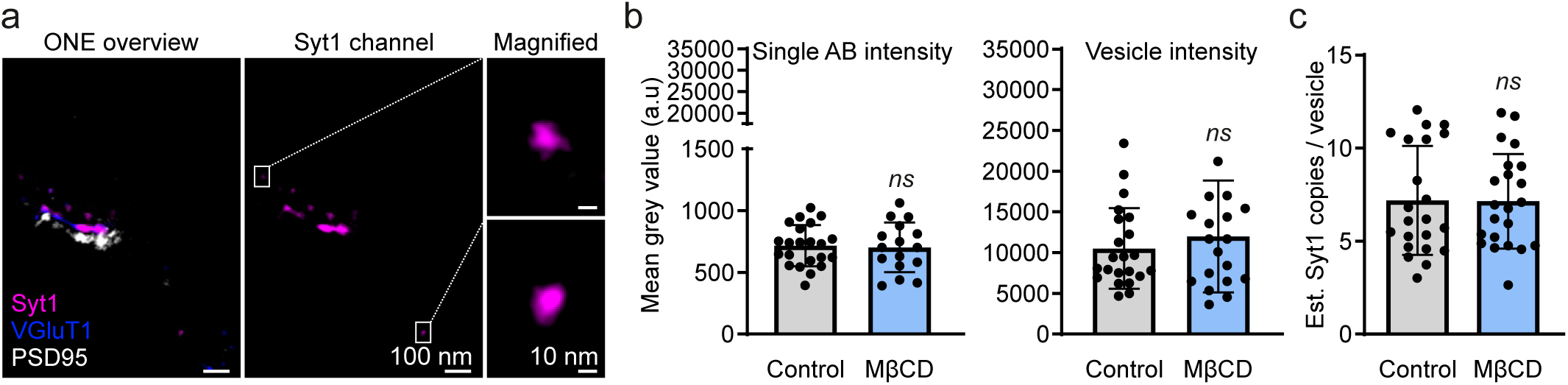
Syt1 count analysis. **a**, The first panel shows a ONE overview image of Syt1, VGluT1, and PSD95 channels. The middle panel shows the Syt1 channel alone, with two selected regions indicating signal from isolated antibodies. The two regions are magnified and displayed on top of each other in the third panel. **b**, The first graph shows the mean grey value of isolated spots in control and MβCD treated neurons. The second graph shows the mean grey value of vesicles. N = 22-19, 2 independent experiments, Mann-Whitney test, *p =* 0.6507 for the first graph and *p* = 0.8494 for the second graph. **c**, A graph showing the number of Syt1 antibodies per vesicle. Syt1 antibody numbers were estimated by dividing the vesicle intensity value by single AB intensity value. Mann-Whitney test; *p* = 0.8937.

**Supplementary Fig. 18.**
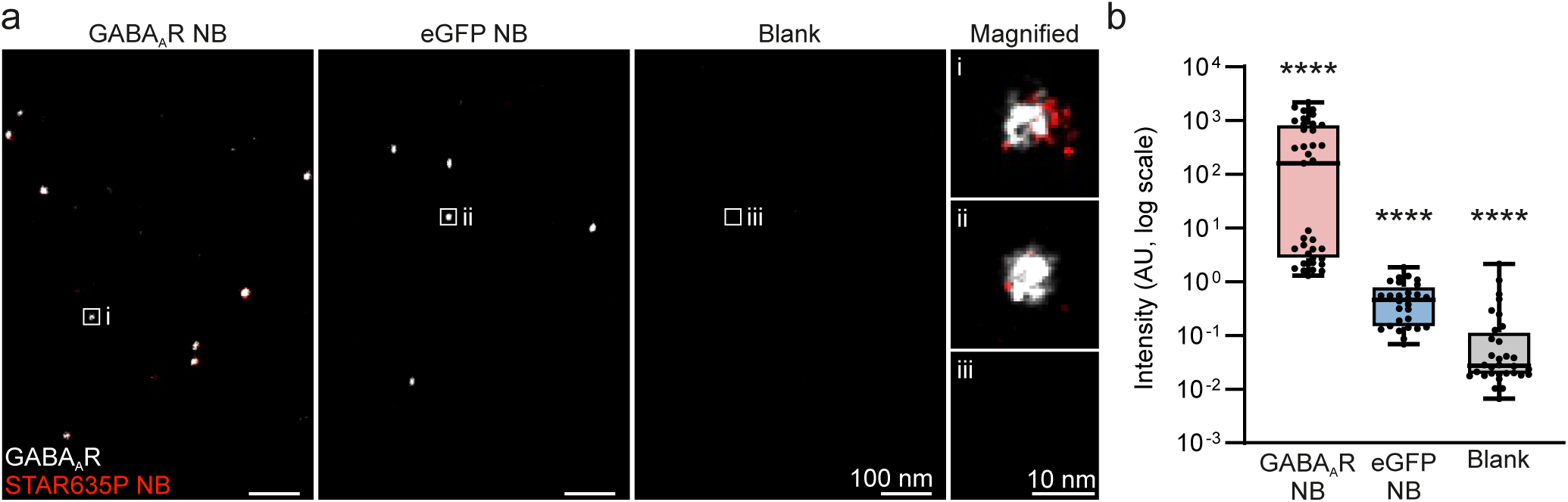
Intensity analysis. Specific nanobodies, which detect GABA_A_Rs, are compared to non-specifically bound eGFP nanobodies and to background noise. **a**, A set of images that shows GABA_A_Rs bound to their respective nanobodies, GABA_A_Rs + eGFP NBs, and a blank control. **b**, Fluorescence intensities were analyzed across the different conditions. N = 37, 28, and 33 images for GABAAR NB, eGFP NB and blank, respectively, from 2 independent experiments.

**Supplementary Fig. 19.**
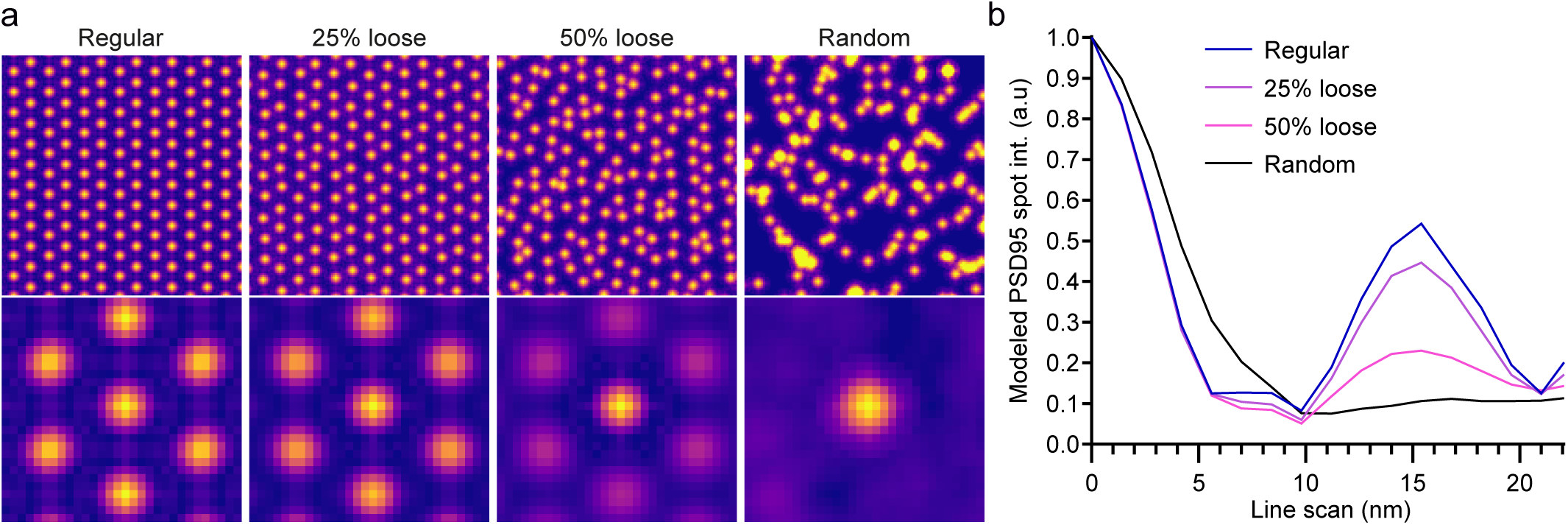
PSD95 model. **a**, To complement the distance analysis presented in Extended Data Fig. 7b, we analyzed the PSD95 distribution using a spot averaging procedure similar to a Ripley curve profile. To explain this analysis in more detail, we modeled it here. The top row of panels shows PSD-like spots, placed in a perfectly regular arrangement (left), with positions varying by 20 or 50% from perfect regularity (middle), or placed randomly (right). The bottom rows of panels show average spots, obtained by overlaying the areas surrounding each of the individual spots in the model arrangements from the top panels. This procedure results in arrangements in which the central spot is surrounded by increasingly weak spots, with virtually no regular spots around it in the right-most panel. **b**, Lines were drawn from the center of each spot in the bottom panels in panel a, in all directions, and were then averaged. The average line going from the center of a spot to the periphery shows a prominent peak if the arrangement is regular, since the neighboring spots are always present at a set distance, and thus provide a visible intensity peak. The less regular the arrangement is, the less clear the second peak becomes. It disappears completely when the spot positions are fully random.

**Supplementary Fig. 20.**
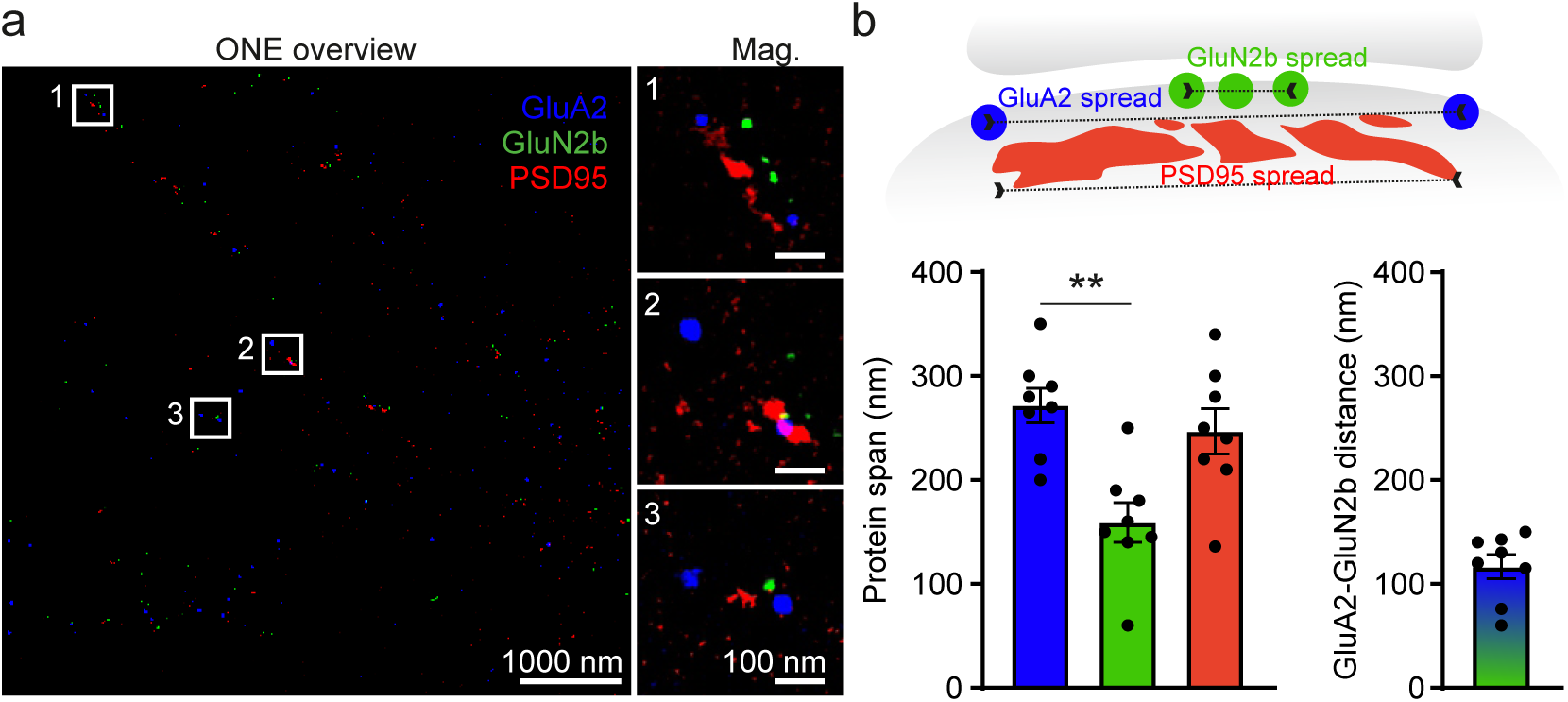
Distribution of post-synaptic elements. **a**, A ONE overview of a hippocampal culture immunostained for GluA2, GluN2b and PSD95. **b**, A cartoon that color-codes the analysis presented in the lower two graphs. The lateral span of the GluN2b, GluA2 and PSD95 signals is presented in green, blue and red. The second graph shows the distance between the center of the GluN2b cluster and the GluA2 periphery. This implies that the GluN2b cluster is positioned relatively close to the center of the GluA2 distribution, since the value here is very close to the half of the GluA2 span. N = 8 post-synapses analyzed, 2 experimental replicates. Kruskal-Wallis test was applied, *p =* 0.0049 **.

**Supplementary Fig. 21.**
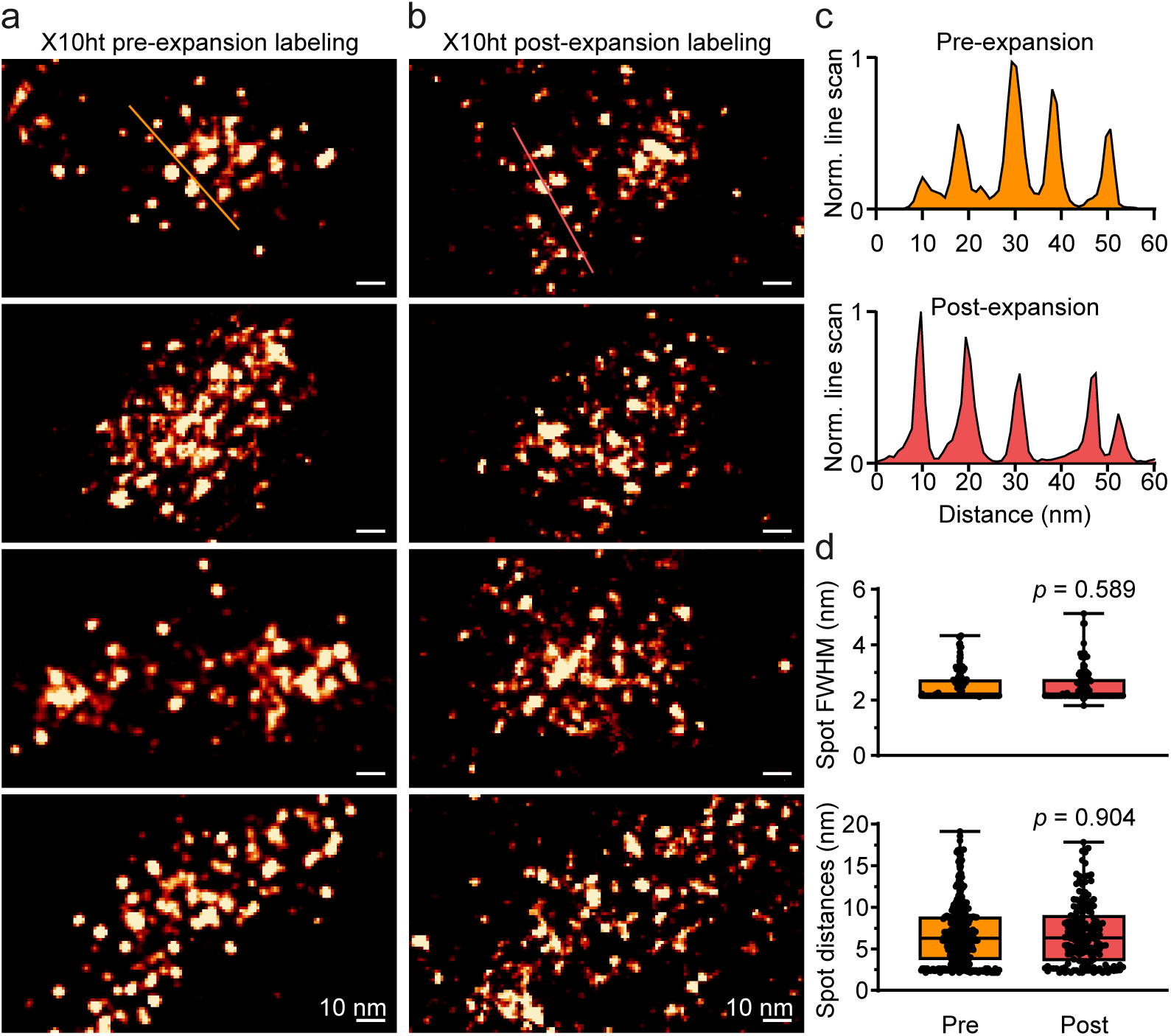
ONE analysis of PSD95 labelled post-expansion. Neuronal cultures were fixed, expanded using the X10ht protocol and labelled after expansion using a PSD95 NB conjugated to Atto488. **a**, For comparison purposes, pre-expansion labelled PSD95 images are reproduced from Fig. 5f (top panel) and **Extended Data Fig. 6a** (the other 3 panels) of the original manuscript. **b**, Post-expansion labelled PSD95 examples. **c**, Exemplary line scans over pre-and post-labelled PSD95 show similar cluster spacing. **d**, Spot FWHM and spot distances measurements showed similar values. N = 140, and 113 measurements spot FWHM, and 402, and 172 for spot distances, for pre-and post-expansion, respectively. Mann Whitney test, non-significant *p* > 0.05.

**Supplementary Fig. 22.**
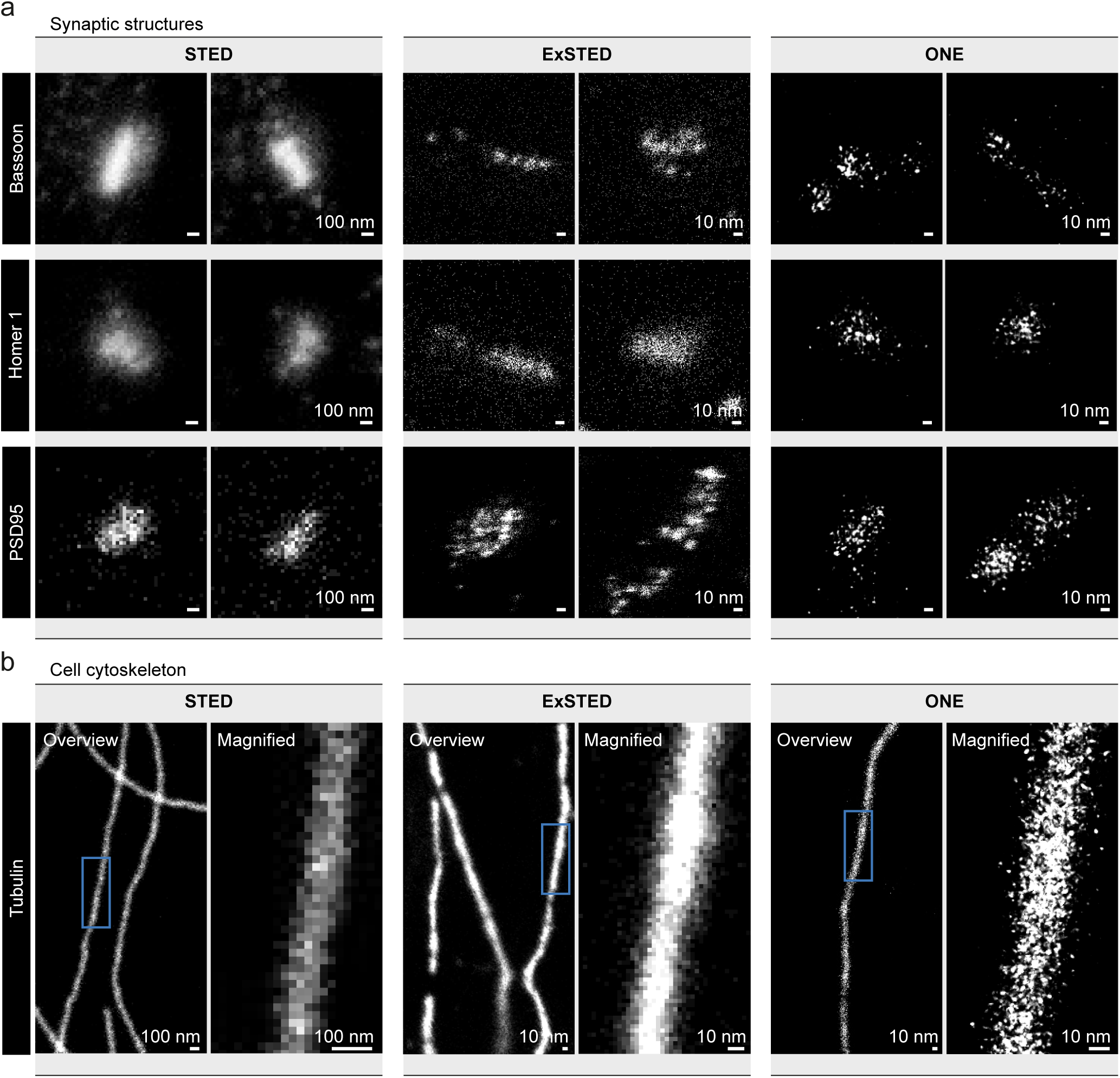
STED, ExSTED and ONE comparison. **a**, The synaptic proteins Bassoon, Homer1 and PSD95 were imaged using STED, ExSTED (X10 expansion combined with STED), and ONE microscopy. **b**, The same procedure was applied to tubulin.

**Supplementary Fig. 23.**
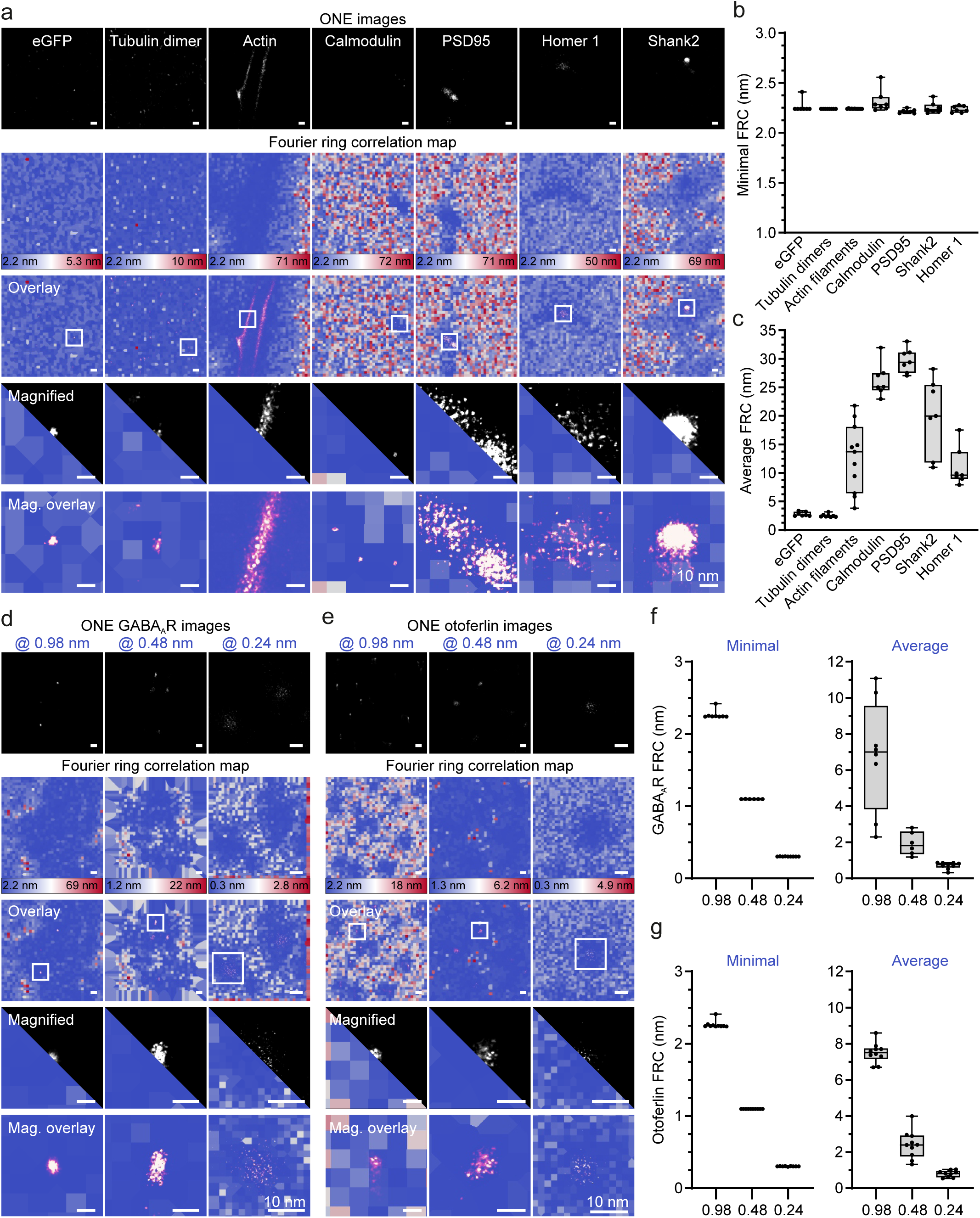
Detailed analysis of Fourier ring correlation. **a**, FRC analysis of ONE images collected with a pixel size of 0.98 nm. The first panel row shows ONE images of the different specimens. The second row shows the corresponding FRC maps. The third row shows ONE images overlaid over FRC maps, using a screen-blend mode. The fourth and fifth rows show magnified views. **b**, A graph plotting the minimal FRCs in nm. **c**, A graph plotting the average FRCs in nm. Please note that all the labelled targets reside in the “bluest” regions of the map, indicating minimal FRCs that correspond to high resolution. N = 7, 8, 12, 7, 7, 7, and 7 for eGFP, tubulin dimer, actin, calmodulin, PSD95, Homer 1, and Shank2, respectively. **d** & **e**, FRC analysis of ONE images achieved with a pixel size at 0.98, 0.48, and 0,24 nm for GABA_A_R and otoferlin, respectively. **f**, The graphs shows minimal and average FRCs in nm for GABA_A_R ONE images. N = 8, 6, and 9 for 0.98, 0.48, and 0.24 nm images, respectively. **g**, The graphs shows minimal and average FRCs in nm for otoferlin ONE images N = 10, 10, and 9 for 0.98, 0.48, and 0.24 nm images, respectively. All experimental sets were performed with at least 2 replicates.

**Supplementary Fig. 24.**
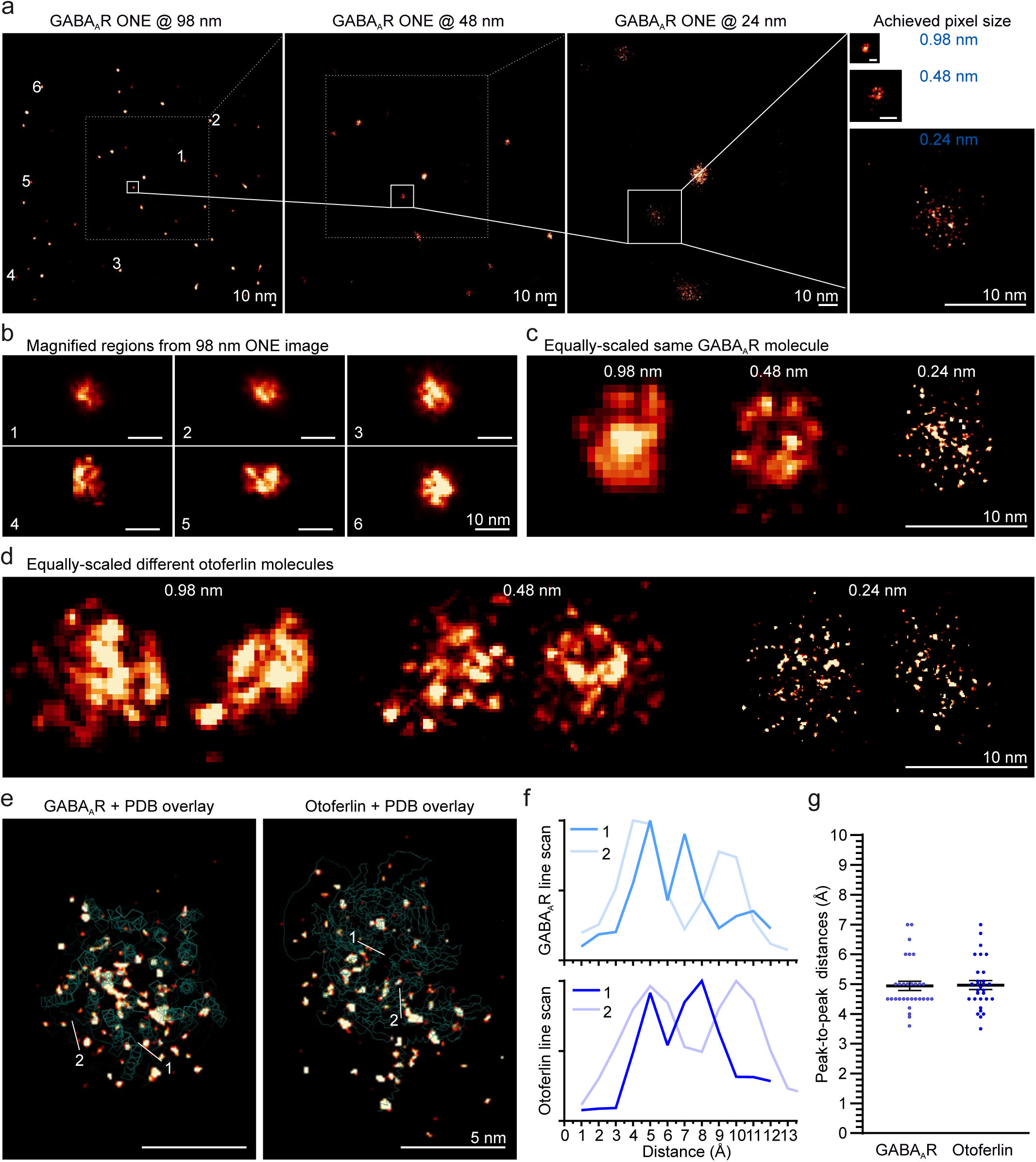
Intra-molecular measurements. **a**, GABA_A_R ONE images acquired with 0.98, 0.48, and 0.24 nm pixel sizes, for the same region. **b**, GABA_A_R magnified examples from the first image in the panel above. **c,** One particular GABA_A_R molecule displayed at different resolutions. **d,** Equally-scaled otoferlin molecules acquired at different resolutions. **e**, ONE images of GABA_A_R and otoferlin at 0.24 nm overlaid with their respective PDBs. **f**, The graphs show 2 exemplary line scans for peptide segments in GABA_A_R and otoferlin. **g**, A graph showing peak-to-peak distances in Ångström. N = 30 for GABA_A_R, and 30 for otoferlin, 10 independent experiments for GABA_A_Rs and 4 independent experiments for otoferlin.

**Supplementary Fig. 25.**
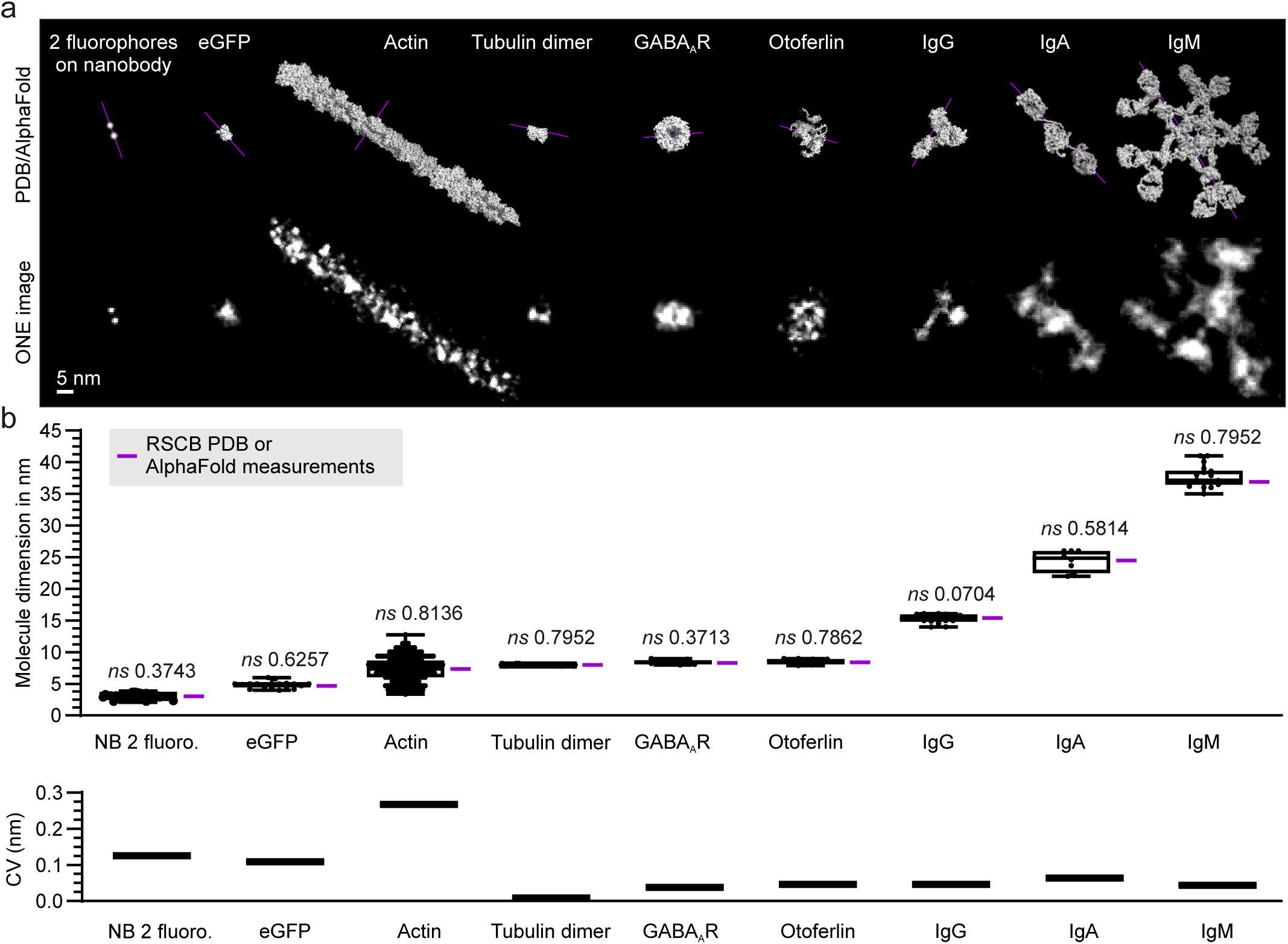
Expansion precision evaluation. **a**, A direct compassion between single-molecule ONE images and their respective PDB/AlphaFold models. The purple line indicate the line scan used to measure the molecule dimension indicated in the first graph. **b**, The upper graph shows measurements of molecule dimensions, in nm. The horizontal purple line indicates the expected value, obtained from measurements of PDB structures (for all molecules except otoferlin), or AlphaFold predictions (for otoferlin). The lower graph shows the variability of these measurements, in the form of the coefficient of variance. N = 34, 17, 192, 75, 10, 10, 14, 8, and 18 for NB, eGFP, actin, tubulin dimer, GABA_A_R, otoferlin, IgG, IgA, and IgM; at least 2 experimental replicates were carried for all experiments. Paired t tests were carried out to determine whether the measured values are different from the values predicted by the PDBs; the respective *p* values are reported above the plots.

**Supplementary Fig. 26.**
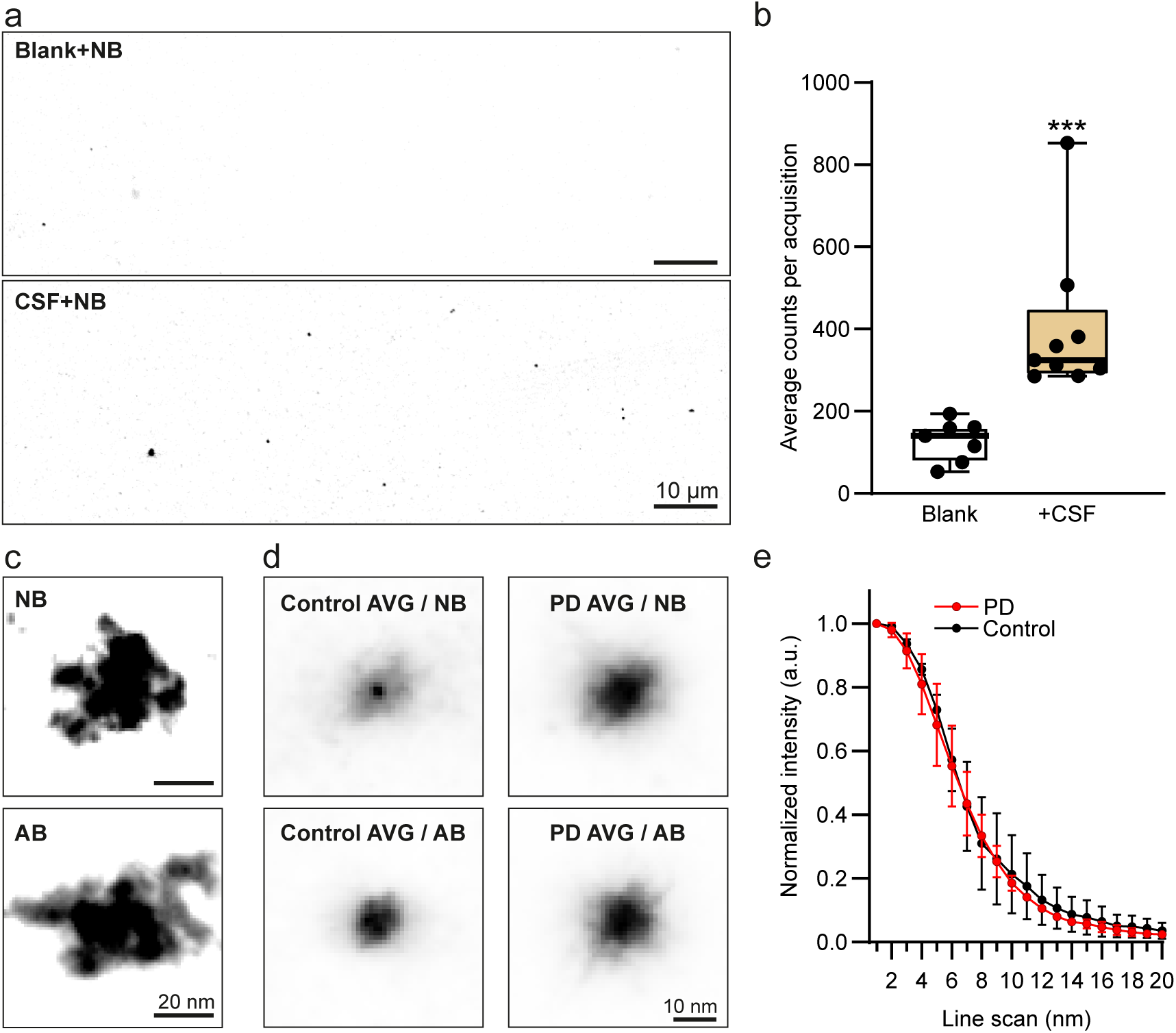
The nanobody imaging of ASYN objects is specific and is not easily reproduced by antibodies. **a**, Low-resolution images of CSF-containing samples, or blanks (clean, BSA-coated coverslips). Only a few dim spots, presumably representing single nanobodies, are seen in the blanks. **b**, Quantification of the signal intensity, as a sum across all image pixels. N = 7-9; Mann-Whitney test, *P* = 0.0002. **c**, Individual examples of oligomers immunolabelled with nanobodies (top) or antibodies (bottom). **d**, Averages of ASYN objects from individual patients, immunolabeled with nanobodies or antibodies. **e**, An analysis of the average object size in antibody-labelled samples, as in Fig. 4. N = 2 patients for each condition; the graph shows mean ± range of values. Nanobodies reveal differences between patients, at object sizes of only a few nm. Antibodies have difficulties in this direction, as their large size causes a lower-fidelity labelling, and as their sizes obscure the actual sizes of small objects.

**Supplementary Fig. 27.**
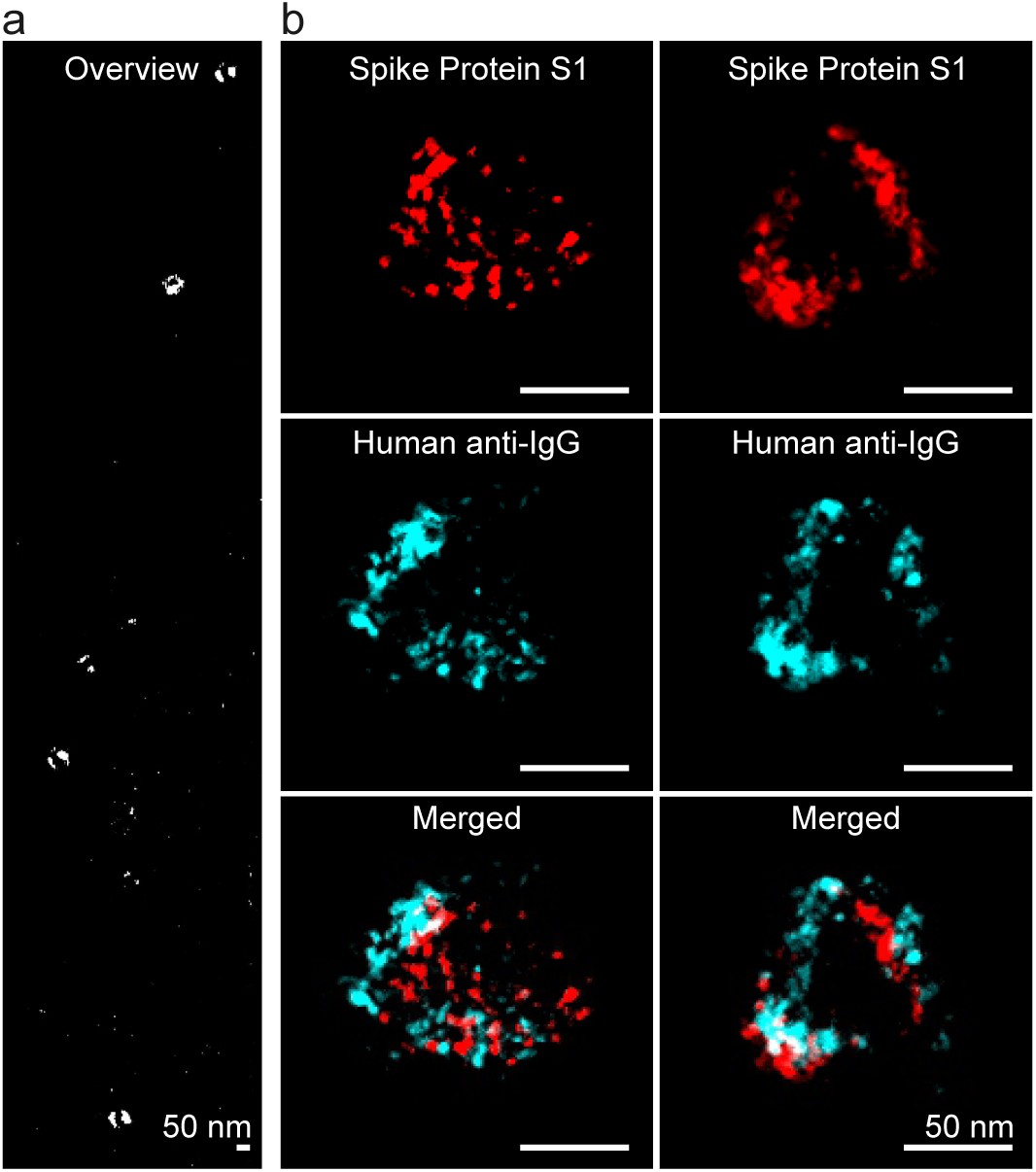
ONE analysis of SARS-CoV-2 viral particles. **a**, ONE overview of a sample containing SARS-CoV-2 viral particles immunostained against Spike Protein S1. **b**, More detailed views of two particles, indicating the Spike Protein S1 and the native IgG molecules from the serum of the patients. Interestingly, a domain-like structure is observed, which is presumably induced by the native IgGs gathering the spike proteins together, by the dual binding capacity of the IgG molecules.

**Supplementary Fig. 28.**
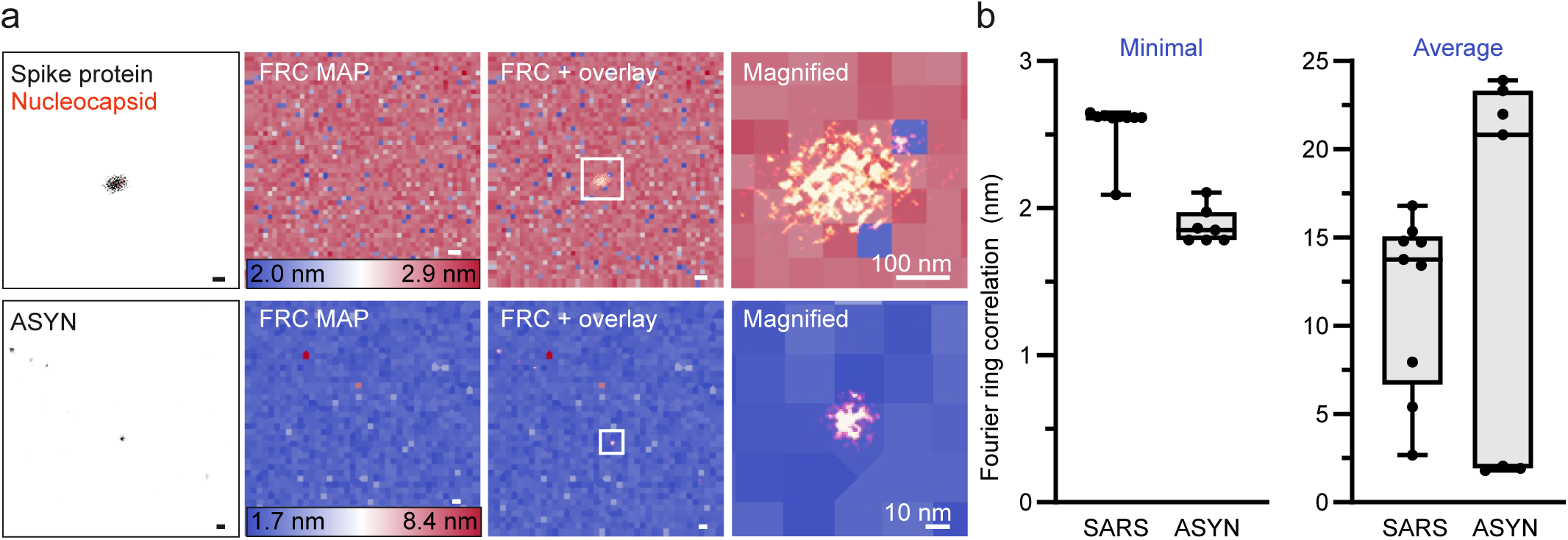
FRC Analysis of SARS-CoV-2 and ASYN aggregates. **a**, FRC analysis of ONE images achieved at 0.98 and 0.78 nm per pixel for SARS-CoV-2 and ASYN, respectively. The first panel shows ONE images of the two specimens. The second panel shows the corresponding FRC maps. The third panel shows ONE images overlaid with FRC maps, using screen-blend mode. The fourth panel shows magnified overlays. Please note that the FRC scale for SARS-CoV-2 ranges between 2.0 and 2.9 nm, with red color in this dataset still indicating very good FRC values. **b**, The graphs plot the minimal and average FRCs in nm. N = 10, and 8 for SARS-CoV-2 and ASYN, respectively, 2 experimental replicates.

**Supplementary Figures 29.**
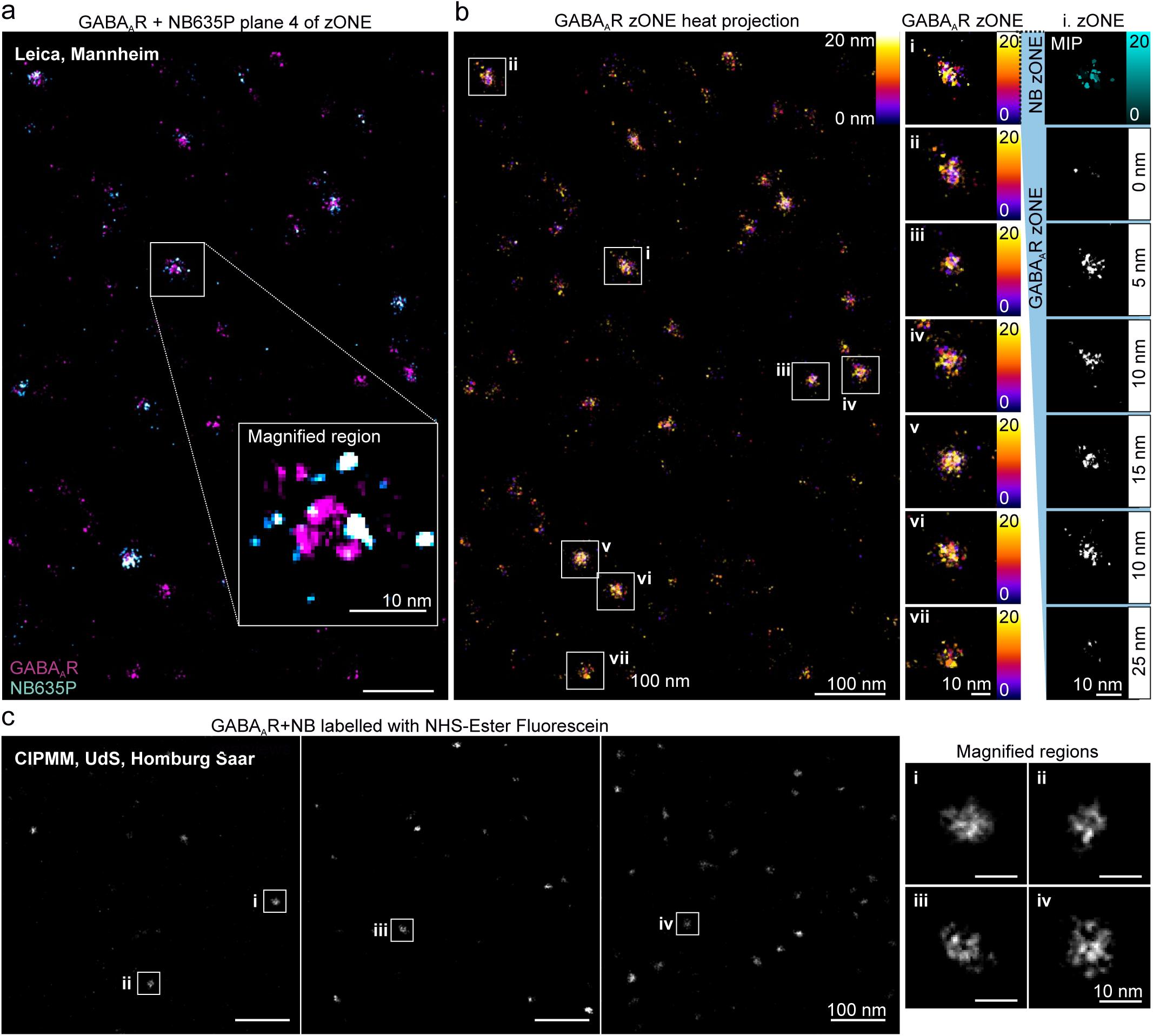
ONE microscopy applied at the confocal headquarters of Leica Microsystems and at the Center for Integrative Physiology and Molecular Medicine (CIPMM) of the Saarland University (UdS). As GABA_A_R were systematically investigated in this study, we chose them as a reference to evaluate the applicability of ONE technique at different laboratories using different systems. **a,** Using a STELLARIS 8 microscope at Leica Microsystems, we present a snapshot of a plane from a 5-dimension x,y,z,c,t image of GABA_A_R+NB. **b**, The first panel shows a depth projection of a zONE stack. The second panel shows a set of GABA_A_Rs that were magnified. The optical sectioning of the first example is displayed in the rightmost panel. **c**, GABA_A_Rs were also successfully imaged at CIPMM, Saarland University (UdS), as shown in full 3 full-scale overviews and in their respective magnified regions. Several microscopes were used at the CIPMM, which are presented in the next figure.

**Supplementary Figures 30.**
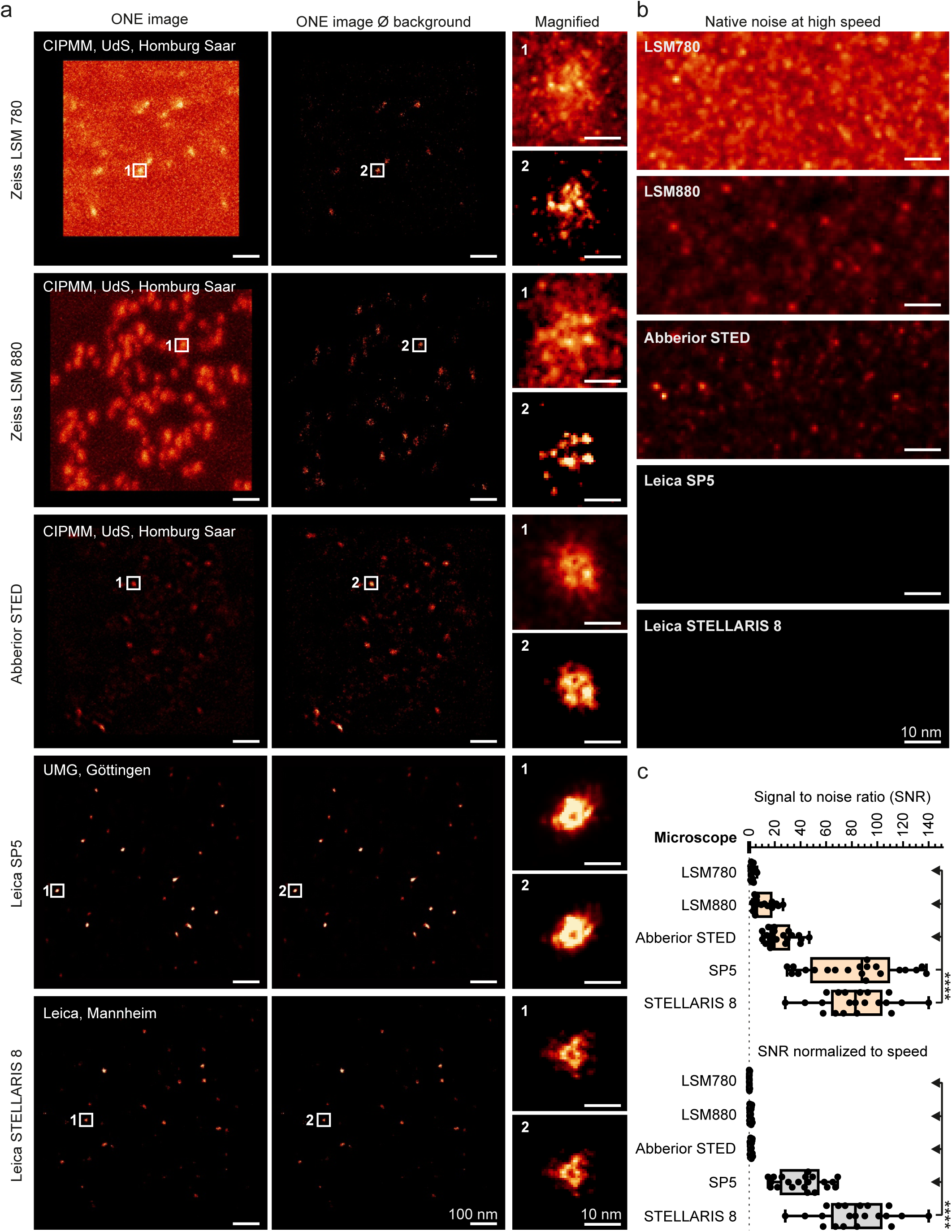
GABA_A_Rs could be imaged with different microscopes. Acquisition settings were matched among different systems to the level that each system allowed. The highest achievable speeds were used for each system. This was systematically characterized (data not shown, but can be presented upon request). **a**, Images from the first panel show GABA_A_R ONE images acquired from different microscopes. As the imaging systems were pushed to their speed limit, background noise was substantially higher on older models. The second panel shows ONE images with background subtraction. The third panel shows a magnified receptor example. **b**, Magnified regions showing the noise readings of each of the used microscopes. **c**, A graph showing the achievable signal-to-noise ratio (SNR), as well as the SNR normalized to acquisition speed. Not surprisingly, higher SNRs yielded better ONE images. N = 22, 22, 22, 23, and 23 for LSM780, LSM880, Abberior STED, SP5, and STELLAIRS 8, respectively, Kruskal-Wallis test was applied *p* < 0.0001 ****.

**Supplementary Figures 31.**
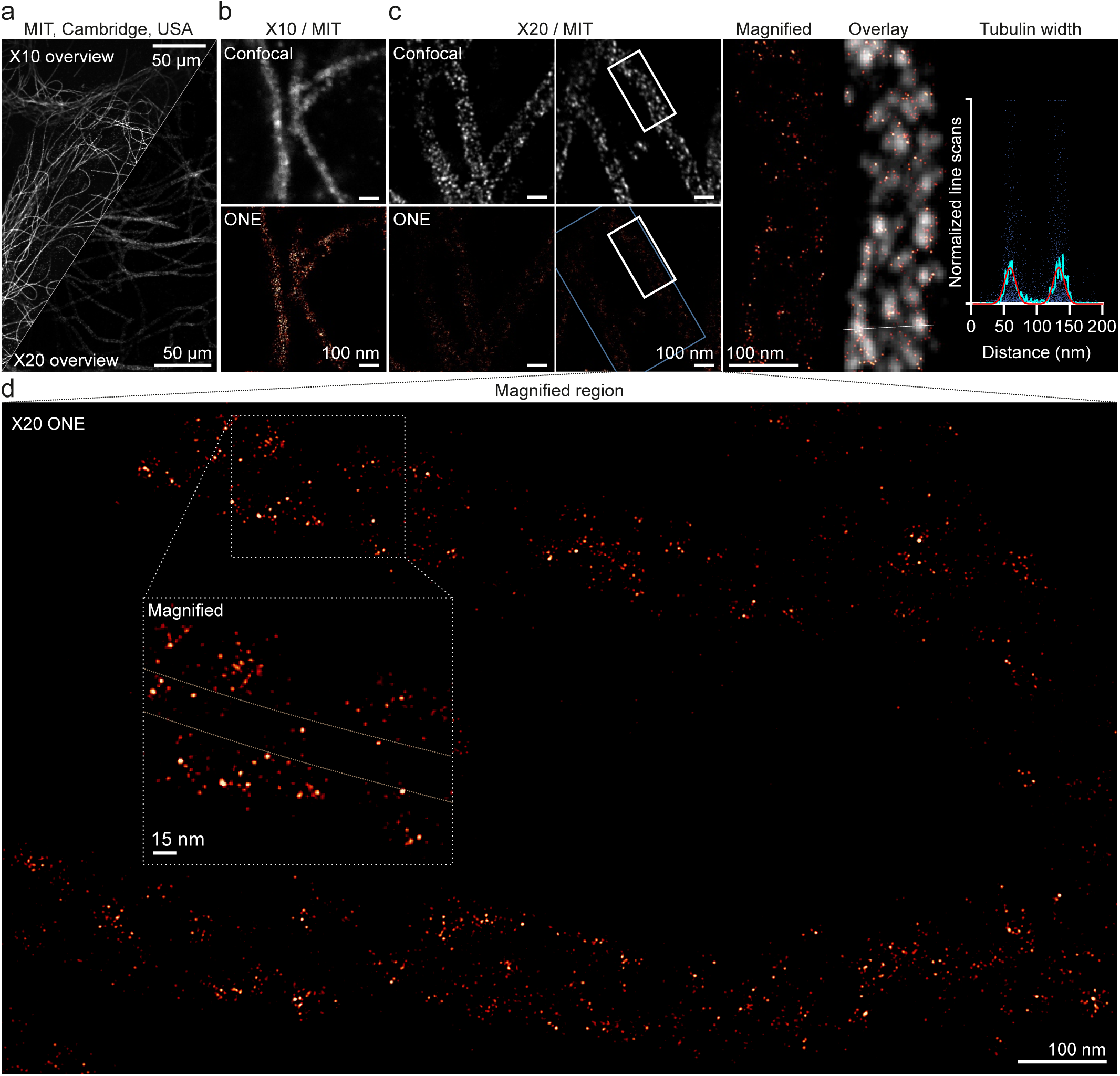
ONE microscopy applied at the MIT, Cambridge, USA. Pre-expansion labelled tubulin specimens were expanded X10 and X20 and were imaged on a STELLARIS 8 system. The X20 gel recipe was modified from the ExR protocol^49^, as follows. The first expansion gel components are 17.25% (w/v) sodium acrylate, 5% (w/v) acrylamide, 1mM BIS, 1x PBS, 0.05% (w/w) APS/TEMED. The re-embedding gel is composed of 10% acrylamide, 2.5 mM BIS, water, 0.05% (w/w) APS/TEMED. The second expansion gel is composed of 17.25% (w/v) sodium acrylate, 5% (w/v) acrylamide, 1mM BIS, 1x PBS, 0.05% (w/w) APS/TEMED. **a**, An overview of tubulin from X10 and X20. **b**, The upper panel shows an X10 confocal image, with its respective ONE image in the lower panel. **c,** The first upper two panels show X20 confocal images of tubulin. Their respective ONE images are shown in the lower panel. Note the significantly dimmer images, as the expansion factor gets higher. The third panel shows a magnified region of X20 ONE, followed by an overlay of confocal and ONE images. The graph shows normalized line scans across the tubulin width. The cyan curve shows the average line scan and the red curve is a fit using a double Gaussian formula. N = 26 line scans. **d**, A magnified region of X20 ONE image is shown. Another portion was magnified and displayed in a white box. The dotted pale-yellow lines are an estimation of the tubulin structure.

**Supplementary Figures 32.**
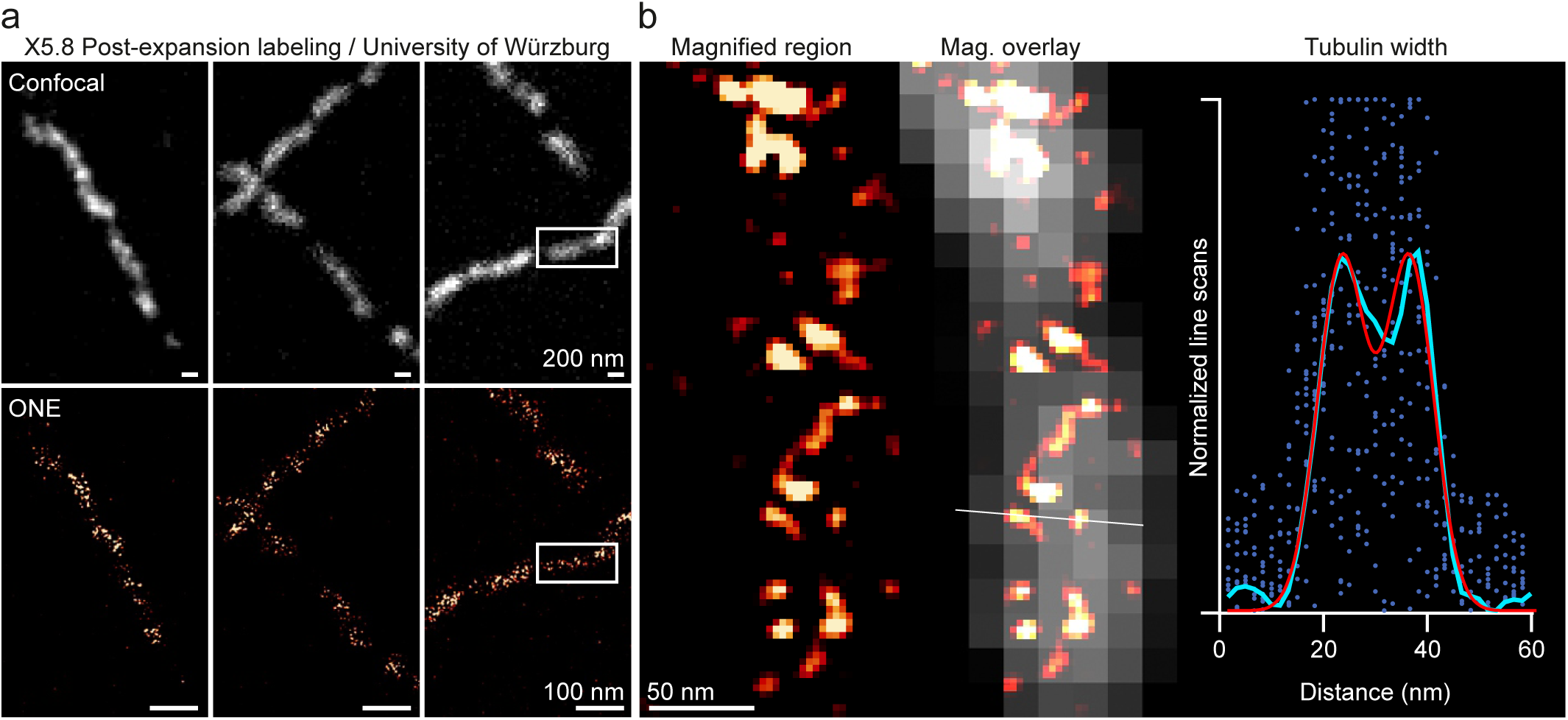
ONE microscopy applied at the University of Würzburg, Germany. Specimens were expanded X5.8 and were post-expansion labelled for tubulin, before imaging using an LSM900 microscope. **a**, The upper panels show a confocal overview and the lower panels show the respective ONE images. **b**, The upper panel shows an X5.8 confocal image and its respective ONE image in the lower panel. **c**, The first image is a magnified region of X5.8 ONE followed by an overlay of a confocal and a ONE image. The graph shows normalized line scans across tubulin width. The cyan curve shows the average line scan and the red curve shows the respective fit. N = 20 line scans.

**Supplementary Figures 33.**
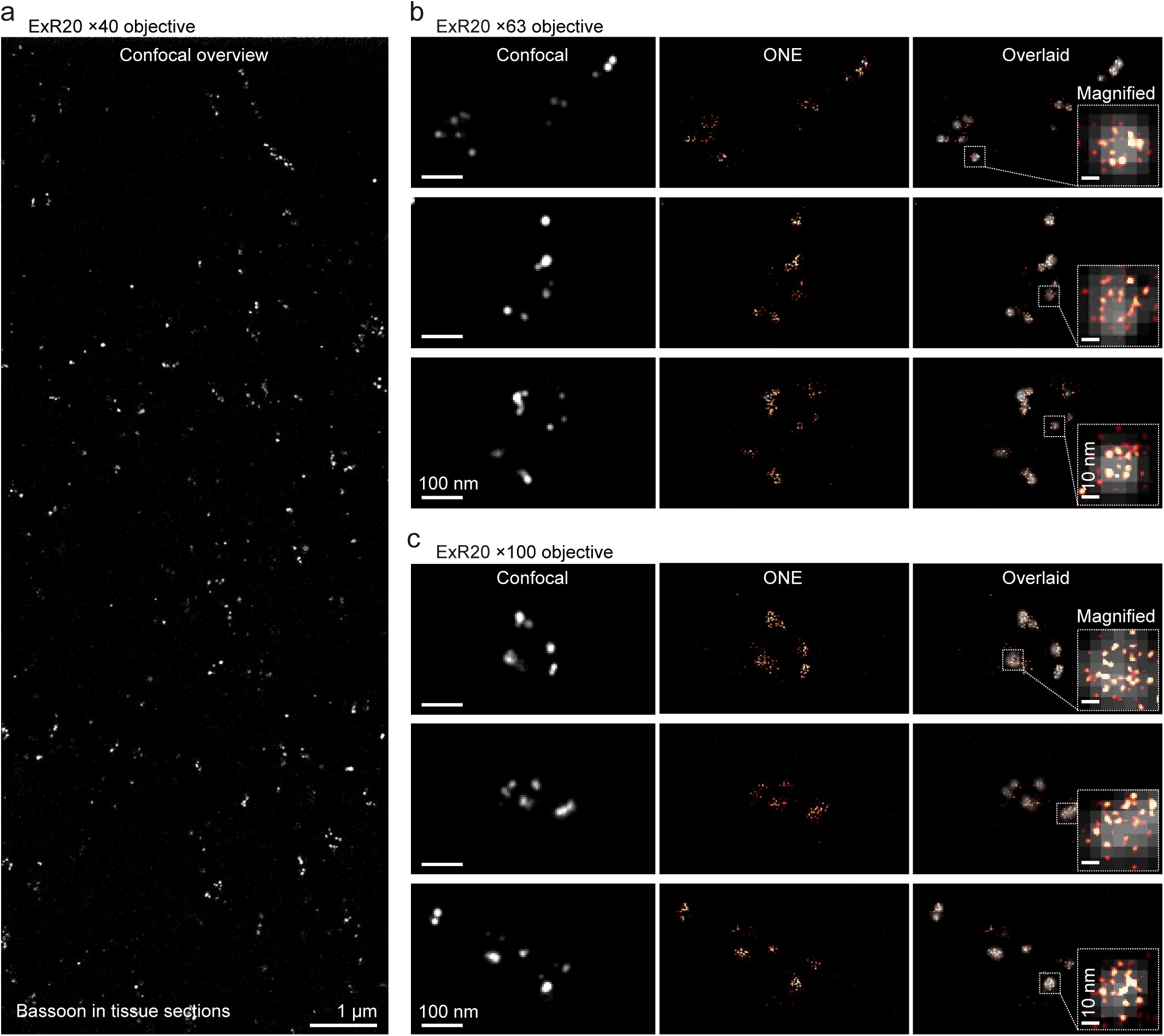
Post-expansion bassoon labeling ONE images. Tissue sections were expanded and then labelled against bassoon following the expansion revealing (ExR) protocol^49^ at the MIT, Cambridge, USA. **a**, An ExR20 (X20 expansion) confocal overview imaged with a ×40 objective. **b**, Three different exemplary ONE images of bassoon using a ×63 objective. The first image is a resonant scan MIP of 20 frames, followed by a ONE image, and an overlay with its respective confocal image. The white square indicates the magnified region to the right. **c**, Similar to (**b**) but using a ×100 objective.

## Supplementary Tables

**Supplementary Table 1.** Patient details.

| ID | Sex | Age | Diagnosis |
| --- | --- | --- | --- |
| 1180 | m | 74 | PD |
| 1407 | m | 82 | PD |
| 1698 | m | 83 | PD |
| 1057 | f | 74 | PD |
| 1081 | m | 69 | PD |
| 1100 | m | 71 | PD |
| 1119 | f | 84 | PD |
| 861 | f | 60 | RLS |
| 906 | m | 73 | CBD |
| 1059 | m | 70 | PSP |
| 1223 | f | 77 | PNP |
| 1382 | f | 75 | PNP |
| 1529 | m | 84 | PNP |
| 1606 | f | 65 | PNP |

Average ages: $76.7 \pm 2.3$ years (PD), $72.0 \pm 2.9$ years (controls); no significant difference (Mann-Whitney Ranksum test).
PD: Parkinson's disease. CBD: Corticobasal degeneration. PNP: Peripheral neuropathy. PSP: Progressive supranuclear palsy. RLS: Restless legs syndrome.

**Supplementary Table 2.**
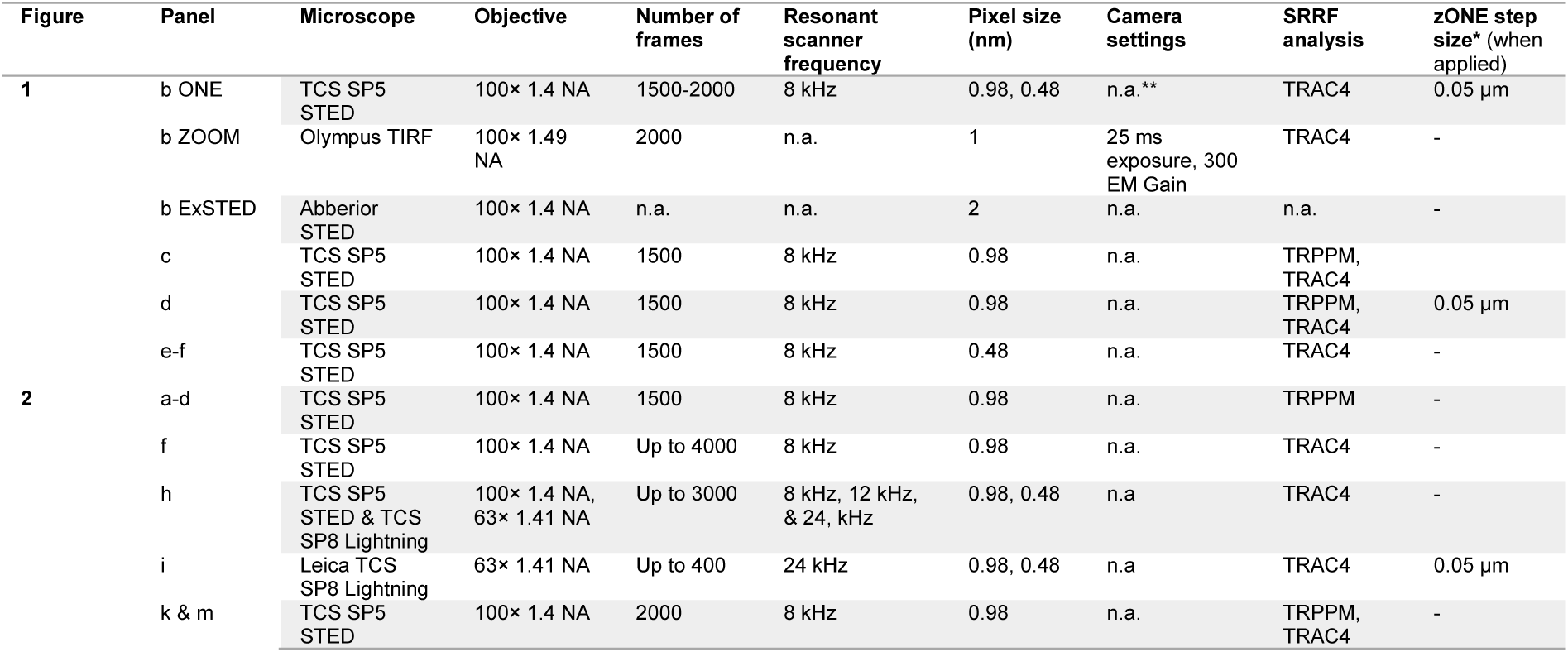
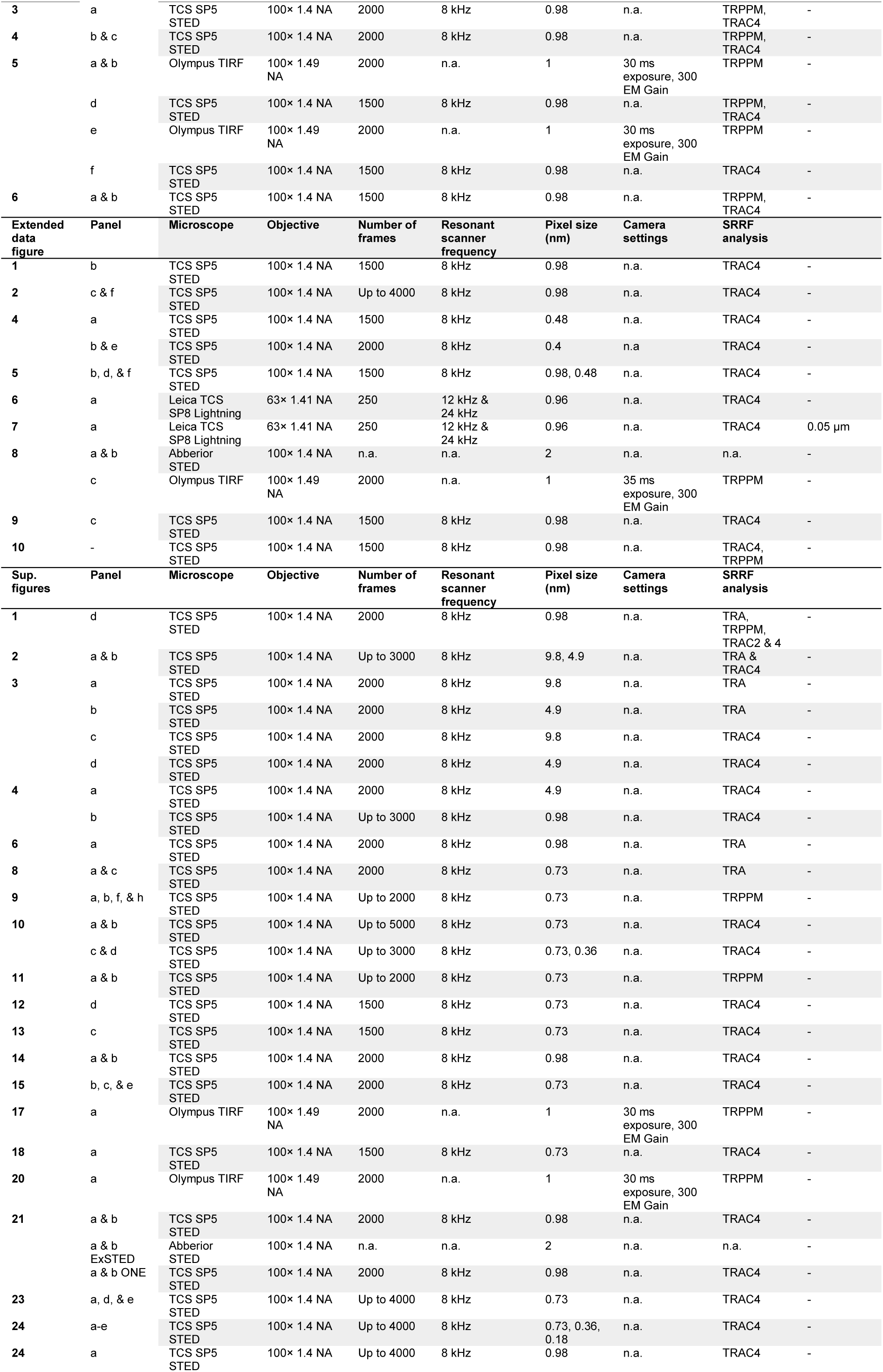
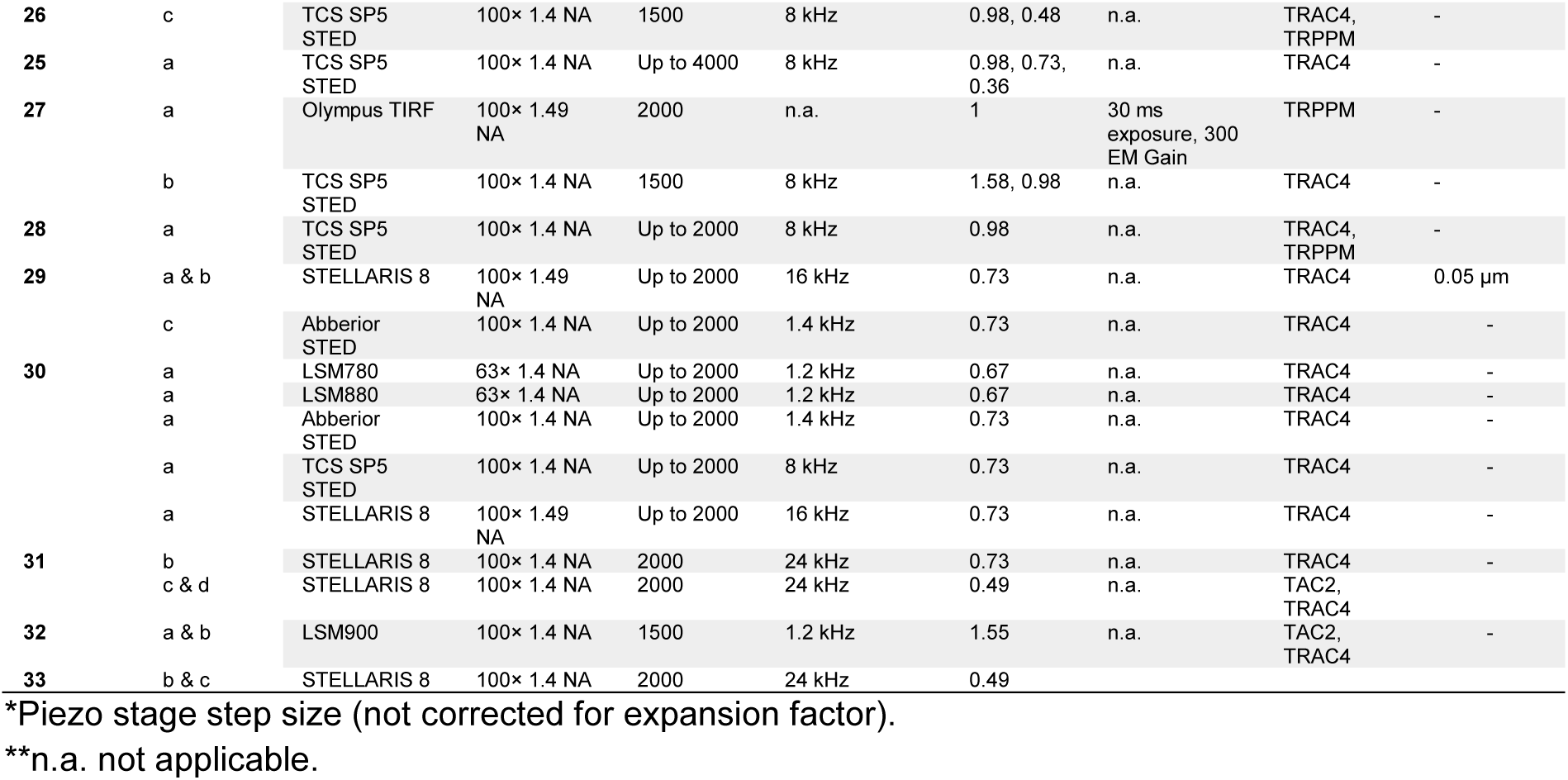
Image format and analysis technical information.

## Supplementary File 1

### ONE Platform plugin manual

This file consists of a series of screen views explaining how the one plugin functions. Please follow the instructions in the respective images.

## Supplementary Discussion

### Resolution

As presented in the main text, the ONE resolution enhancement relates almost exclusively to the lateral (XY) plane. Resolution along the Z axis depends on the expansion factor of the gel, being equivalent to the axial resolution of the confocal microscope used, divided by the expansion factor. This results in a difference of more than 20-fold between the axial and the lateral resolution, which will have significant effects on the image quality. This situation parallels conventional transmission electron microscopy (TEM), in which the thickness of the specimen limits the axial resolution to a similar 20-to 40-fold above the lateral resolution.

This situation implies that the optimal samples for ONE imaging would have a limited number of objects within the axial imaging volume of 40-60 nm (pre-expansion; volume calculated for a conventional confocal microscope and a 10-15x expansion factor). Denser structures will cause a signal overlap that will confuse the identification of individual structures. The use of purified proteins, which can be diluted to the desired signal density, is an optimal application for ONE microscopy, since the dilution factor avoids the potential issues with axial resolution. The axial resolution problem is especially evident for microtubules, whose thickness is not sufficiently large, under 10x expansion, to avoid imaging the entire microtubule structure within one confocal volume. Therefore, the entire “tube” of the microtubule appears as a band of fluorescence in the images shown in Fig. 1. When the expansion factor is raised beyond 10x, this is no longer a problem, and the sides of the microtubule become evident.

We have not encountered any issues relating to the sample density in the lateral (XY) plane: the shape of individual proteins is maintained well, and all measurements we performed provided results within the expected boundaries.

### Sample anchoring into the gel

We are currently relying on NHS-ester chemistry to anchor proteins into the gels, using the well-established chemical Acryloyl-X. This molecule reacts to amine groups on lysines and on the N-termini of the proteins in the sample. As lysines make up ∼5% of all amino acids in proteins, most proteins should have sufficient anchor points for accurate gel anchoring. The only problem we can envision is the fact that aldehyde fixatives also modify amine groups. If gel anchoring appears faulty in specific samples, possibly due to excessive fixation, we suggest using an epitope retrieval strategy, in which the sample is heated to 95°C in basic buffers (pH 8-9). This strategy should eliminate some of the fixative effects, and should enable accurate gel anchoring. Performing the anchoring in basic buffers, overnight, should also assist with this issue.

### Homogenization

The heat-based homogenization is optimal for retaining fluorophores already present in the samples (pre-expansion labeling), since it breaks the proteins, but it does not proceed, in the version we optimized, to the removal of every amino acid. At the same time, it does not rely on the diffusion of an enzyme deep into tissues, so it is optimal for these preparations. In contrast, the proteinase K presumably removes all amino acids that are not anchored into the gel. This approach is optimal for single proteins, since the fluorophore positions become quite precise, being always near the anchor points. However, proteinase K diffusion in thick tissues is poor, and therefore this approach is not suited for tissue slices of over ∼10 µm.

### SRRF performance

The initial implementation of SRRF resulted in a 50-70 nm resolution^6^, leading to the impression that this is the best achievable resolution for this technique, as it is implied by some subsequent works (e.g.^32^). This is not the case, as demonstrated in our work on nanorulers (**Supplementary Figs. 2-4**). The name SRRF serves as an umbrella term for a number of different analyses, including the temporal radiality average (**TRA**) and temporal radiality auto-correlation (**TRAC**)^6^. The latter method is a higher-order statistical analysis (following the procedures initially introduced for SOFI^26^, whose contrast, accuracy and final resolution are substantially higher than those of the TRA method. The TRA analysis does not consider higher-order temporal correlations, which makes it comfortable to use with limited numbers of frames (*e.g*. 100-300 frames), thus rendering it a method of choice for live-cell SRRF^6, 27^. TRA is heavily dependent on the distance between the fluorophores, and performs best when the different fluorescent objects are separated by more than 70% of the full width at half maximum (FWHM) of the point-spread-function (PSF^32^). This implies that this procedure is not intended to produce a very high resolution, unlike the TRAC analyses. These analyses do produce better resolutions, but require larger numbers of frames for optimal performance, something that does not seem to be clear in the literature, since all SRRF implementations are often performed with as few as 100 frames. Nevertheless, the optimal resolution obtained with TRAC analyses can be pushed towards 20 nm, and maybe even beyond this value, under ideal imaging conditions (**Supplementary Figs. 2-4**). We therefore conclude that SRRF should not be considered to be limited to 50-70 nm resolutions, as explained in the Supplementary Notes of the original SRRF publication^6^.

As for most other super-resolution approaches, the pixel size limits the resolution to a value of approximately its double^52^. This limitation can be overcome, as indicated in **Supplementary Fig. 2**, by reducing the initial pixel size. This, however, will result in a lower signal-to-noise ratio (SNR), which is an essential parameter for all fluctuation-based analyses. Even when applied to low numbers of frames, SRRF provided excellent images when the signal-to-noise ratio surpassed 10-15^6, 32^. Below these values, SRRF will perform more poorly than many other related methods, as MUSICAL or ESI^32^, implying that users should consider carefully the noise levels of their images, as explained in **Supplementary Fig. 4**.

The ONE procedure is designed to alleviate two of the main problems of the TRAC analysis, the fluorophore distance and the SNR. First, the distance between the fluorophores increases in all dimensions, leading to their dilution by the third power of the expansion factor. Second, the SNR increases profoundly (**Extended Data Fig. 3**). This is an important side effect of removing all cellular materials that are not embedded into the gels.

The remaining problem, that of acquiring sufficient frames for optimal performance, depends on 1) sample stability, and 2) fluorophore bleaching. The solutions to these issues come in the form of an improved gel-holding chamber (**Supplementary Fig. 5**) and of rapid resonant scanning, which reduces fluorophore bleaching (**Supplementary Fig. 7**). The latter effect is known from other super-resolution fields, as STED^77^ and is probably due to the fact that rapid scanning lowers the light dose received continuously by every fluorophore, thereby reducing the possibility of excessive excitation and damage.

## Materials and Methods

### Nanorulers

Custom-designed linear nanorulers of varying length (80, 60, 50, 30, 20, and 10 nm), carrying one Atto647N molecule on each end, were purchased from GATTAquant GmbH, Gräfelfing, Germany.

### Conventional cell cultures

Tubulin immunostaining was performed in the U2OS cell line, obtained from Cell Lines Service (CLS, Eppelheim, Germany). The cells were grown in a humidified incubator (5% CO_2_, 37°C), in Dulbecco’s Modified Eagle Medium (DMEM #D5671, Merck, Darmstadt, Germany), with the addition of 10% FCS (fetal calf serum, #S0615, Merck) and 4 mM glutamine (#25030-024, ThermoFisher Scientific, Waltham, USA), with an antibiotic mixture added at 1% (penicillin/streptomycin (ThermoFisher Scientific). For imaging purposes, cells were grown overnight on poly-L-lysine-coated coverslips (#P2658, Merck).

### Hippocampal cultured neurons

Animals (Wistar rats, P0 to P1) were treated according to the regulations of the local authority, the Lower Saxony State Office for Consumer Protection and Food Safety (Niedersächsisches Landesamt für Verbraucherschutz und Lebensmittelsicherheit), under the license Tötungsversuch T09/08. In brief, the hippocampi were dissected from the brains, were washed with Hank’s Balanced Salt Solution (HBSS, #14175-053, Invitrogen, Waltham, MA, USA), before being incubated under slow rotation in a digestion solution containing 15 U/ml papain (#LS003126, Worthington, Lakewood, USA), with 1 mM CaCl_2_ (#A862982745, Merck), 0.5 mM EDTA and 0.5 mg/ml L-cysteine (#30090, Merck), in DMEM. This procedure is performed for 1 hour at 37°C, before enzyme inactivation with a buffer containing 10% FCS and 5 mg/ml bovine serum albumin (BSA, #A1391, Applichem, Darmstadt, Germany) in DMEM. The inactivation solution is replaced after 15 minutes with the growth medium, containing 10% horse serum (#S900-500, VWR International GmbH, Darmstadt, Germany), 1.8 mM glutamine and 0.6 mg/ml glucose in MEM (#51200046, ThermoFisher Scientific), which is used to wash the hippocampi repeatedly. The neurons are then isolated by trituration using a glass pipette, and are sedimented by centrifugation at 800 rpm (8 minutes). The cells are then resuspended in the same medium and are seeded on PLL-coated coverslips, for several hours, before replacing the buffer with Neurobasal-A culture medium (#10888-022, ThermoFisher Scientific), containing 0.2% B27-supplement (#17504-044; ThermoFisher Scientific) and 2 mM GlutaMAX (#35050-038, ThermoFisher Scientific). The neurons are then maintained in a humidified incubator (5% CO_2_, 37°C) for at least 14 days before usage.

### Brain slices

We dissected rat brains from P0-P1 rat pups (Wistar), as above. The brains were then fixed with 4% PFA (#30525894, Merck) in PBS, for 20 hours. The fixed brains were then placed in agarose (4% solution, #9012366, VWR Life Science, Hannover, Germany), before cutting to the desired thickness (100-200 µm) using a vibratome.

### Patients

Patients were in treatment at Paracelsus Elena Klinik, Kassel, Germany. They had been diagnosed with Parkinson’s disease according to standard criteria^78–80^. Neurological control patients had been diagnosed with a variety of non-neurodegenerative disorders. For a detailed presentation of patients, their ages and diagnoses, see Supplementary Table 1. The informed consent of all of the participants was obtained at the Paracelsus Elena Klinik, following the principles of the Declaration of Helsinki.

### CSF samples

CSF samples were collected at the Paracelsus Elena Klinik, Kassel, Germany, following identical standard operating procedures (SOPs). CSF was gained by lumbar puncture in the morning with the patients fasting and in sitting position. The CSF was processed by centrifugation at 2000 x g for 10 minutes at room temperature and aliquots of supernatant frozen within 20-30 minutes and stored at -80 °C until analysis. Samples with red blood cell count>25/µl or indication for an inflammatory process were excluded.

### Preparation of microtubule samples

We reconstituted unlabelled tubulin (T240-A, TebuBio, Offenbach, Germany) to a concentration of 100 μM. We prepared stabilized microtubules by polymerizing through step-wise increase of the tubulin concentration. Initially, a 3 μM tubulin solution in M2B (magnesium 2X buffer: 80 mM PIPES with 1 mM EGTA and 2 mM MgCl_2_, pH 6.8, adjusted with KOH), in the presence of 1 mM GMPCPP (NU-405L, Jena Bioscience, Jena, Germany) was prepared at 37 °C to nucleate short microtubule seeds. Next, the total tubulin concentration was increased to 9 μM in order to grow long microtubules. To avoid further microtubule nucleation, we added 1 μM tubulin at a time from a 42 μM stock solution and waited for 15 min between the successive steps. We centrifuged the polymerized microtubules for 10 min at 13,000 × g to remove any non-polymerized tubulin and short microtubules. We discarded the supernatant and carefully resuspended the pellet in 800 μl M2B-taxol (M2B supplemented with 10 μM taxol (T7402, Merck, Darmstadt, Germany).

### Immunostaining procedures

#### *Tubulin* immunostaining

U2OS cells were first incubated with 0.2% saponin (#47036, Sigma Aldrich), to extract lipid membranes. This procedure was performed for 1 minute in cytoskeleton buffer, consisting of 10 mM MES (#M3671, Merck), 138 mM KCl (#K42209636128, Merck), 3 mM MgCl_2_ (#M8266-100G, Sigma-Aldrich), 2 mM EGTA (Merck 324626-25GM) and 320 mM sucrose, at pH 6.1. The cells were then fixed, using 4% PFA and 0.1% Glutaraldehyde (#A3166, PanReac, Darmstadt, Germany), in the same buffer. Unreacted aldehyde groups were quenched using 0.1% NaBH4 (#71320, Sigma Aldrich now Merck), for 7 minutes in PBS, followed by a second quenching step with 0.1 M glycine (#3187, Carl Roth), for 10 minutes in PBS. The samples were blocked and simultaneously permeabilized using 2% BSA and 0.1% Triton X-100 (#9036-19-5, Sigma Aldrich), in PBS (room temperature, 30 minutes). Primary tubulin antibodies (#T6199 Sigma Aldrich, #302211 Synaptic Systems, Göttingen, Germany, #302203 Synaptic Systems, #ab18251 Abcam, Cambridge, UK) were applied for 60 minutes at room temperature, and were then washed off with permeabilization buffer, followed by an incubation of the samples with secondary antibodies (#ST635P-1001, Abberior, Göttingen, Germany). Alternatively, the primary antibodies were saturated with secondary nanobodies (#N1202-Ab635P-S and #N2402-Ab635P-S, both NanoTag Biotechnologies GmbH, Göttingen, Germany) for 30 minutes at room temperature, using a ratio of 1:5 for the primary antibody:secondary nanobody, respectively. Afterwards, the antibody mixture was diluted in the blocking buffer, and was applied onto the cells for 60 minutes at room temperature. Five washes with permeabilization buffer followed by three PBS washes (each one for 10 minutes), before continuing with cellular expansion.

#### Neuronal immunostainings

Neurons were fixed with 4% PFA in PBS (#D8537-500ML, TheromFisher), for at least 30 minutes, before quenching with 50 mM glycine (in PBS) for 10 minutes, and blocking/permeabilizing using 2.5% BSA (#9048-46-8, Sigma-Aldrich), 2.5% NGS, and 0.1% Triton X-100 (#1003287133, Sigma-Aldrich) in PBS (30 minutes at room temperature, unless specified elsewhere otherwise). The antibodies and/or primary nanobodies were diluted in 2.5% BSA, 2.5% NGS in PBS, and they were added to the coverslips for 60 minutes at room temperature. This was followed by washing with the permeabilization buffer (30 minutes, three buffer exchanges), and by the application of secondary antibodies or nanobodies, in the same buffer, for 45 minutes at room temperature. Specimens were then washed five times with permeabilization buffer and a final wash with PBS was then performed (15-30 minutes, three buffer exchanges). The primary antibodies used were anti-synaptotagmin1 (SYT1, #105011 Synaptic Systems), anti-Homer1 (#160 003, Synaptic Systems), anti-Shank2 (#162204 Synaptic Systems), anti-GluR2 (Alomone Labs, #AGC-005, Jerusalem, Israel), anti-GluN2b (Neuromab 75-101, California, USA), anti-MAP2 (Novus Biologicals #NB300-213), anti-vGluT1 (#135304, Synaptic Systems), anti-Bassoon (#ADI-VAM-PS003-F, Enzo, New York, USA). Primary nanobodies were FluoTag-X2 anti-PSD95 (clone 1B2, #N3702, NanoTag Biotechnologies GmbH). Secondary antibodies were conjugated to Alexa 405 (#ab175674, Abcam), Alexa Fluor 488 (AF488, #706-545-148, Dianova), Cy3 (#711-165-150, Jackson ImmunoResearch), Abberior STAR580 (AS580 #ST580-1006, Abberior), Abberior STAR635P (#2-0112-007-1, Abberior), FluoTag-X2 STAR635P #N2002-Ab635P and #N2402-Ab635P (NanoTag Biotechnologies GmbH). For post-expansion immunostainings of PSD95, tubulin and bassoon, the gels were blocked with 5% NGS in PBS + 0.1% Triton X-100 (PBST) for 2 h, and were then incubated with either PSD95-nanobody, tubulin-or bassoon-antibody at concentrations of 5-10 µg/ml in PSBT overnight at 4 °C. The gels were washed 4x30 min each. PSD95 gels were expanded by adding water, while tubulin and bassoon gels were then incubated with secondary antibodies in PBST overnight. On the next day, the gels were washed 4x30 min each in PBST and we then expanded the gels by adding water.

#### Live immunostaining using synaptotagmin 1 antibodies

Surface Synaptotagmin 1 (Syt1) molecules were first blocked using unconjugated 604.2 Syt1 antibodies (#105311 Synaptic Systems), for 10 minutes at room temperature, in Tyrode buffer lacking Ca^2+^ (to reduce drastically both exo-and endocytosis; the Tyrode buffer contained 124 mM NaCl (#K52190904041, Merck), 5 mM KCl, 2 mM CaCl_2_ (A862982, Merck), 1 mM MgCl_2_, 30 mM glucose, 25 mM HEPES (K45408310520, Merck), at pH 7.4). The neurons were then wash with room temperature-Tyrode buffer and incubated over an ice water bath and exposed to fluorescently-conjugated Syt1 antibodies (#105311AT1, Synaptic Systems) for 40 minutes, to enable limited exo-and endocytosis. The neurons were then washed with ice-cold Tyrode buffer and then were fixed with 4% PFA for 20 minutes, and quenched with 50 mM glycine for 10 minutes. The samples were then blocked with 2.5% BSA in PBS for 30 minutes and vGluT1 antibody was added prior to permeabilization for 1 h. Three brief washing steps with blocking buffer preceded the half-an hour permeabilization step (0.1% Triton, 2.5% BSA, 2.5% NGS in PBS), and neurons were labelled for PSD95 using the FluoTag-X2 anti-PSD95 nanobody (NanoTag Biotechnologies GmbH), as indicated above. Synapses were identified as regions in which vGluT1 and Syt1 signals were found adjacent to the PSD95 staining.

#### Immunostaining of cerebro-spinal fluid (CSF) samples

Cerebro-spinal fluid probes were obtained from PD patients and controls at the Paracelsus Elena Klinik (Kassel, Germany), and were stored at -80°C before use. 20 µl amounts of CSF were placed on BSA-coated coverslips, enabling the sedimentation of multiprotein species overnight at 4° C. Fixation with 4% PFA (10 minutes, room temperature) and quenching with 50 mM glycine (10 minutes, room temperature) was followed by the application of either antibodies (Alpha-synuclein #128211 and 128002, Synaptic Systems) or Alpha-synuclein nanobody2, SynNb2^68^, custom produced and fluorescently-conjugated by NanoTag) for 1 h at room temperature, in 2.5% BSA in PBS buffer. For the case of antibodies, secondary Aberrior STAR635P was applied for 1 h at room temperature. Five washes with 2.5% BSA in PBS were followed by mild post-fixation with 4% PFA for 4 min, and by the expansion procedures.

#### Brain slice immunostaining

The fixed brain slices were first quenched using 50 mM glycine (in PBS), followed by three washes with PBS (each for 5 minutes), and blocking and permeabilization in PBS containing 2.5% BSA and 0.3% Triton X-100, for 120 minutes at room temperature. The primary antibodies used (Bassoon, #ADI-VAM-PS003-F, Enzo Life Sciences GmbH, Lörrach, Germany; Homer1, #160003, Synaptic Systems) were diluted in the same buffer (lacking Triton X-100) to 2 µg/ml and were added to the slices overnight, at 4°C. Three washes with PBS (each for 5 minutes) removed the primary antibodies, enabling the addition of secondary antibodies conjugated with Abberior Star635P (#ST635P-1001, Abberior, Göttingen, Germany) for Basson identification, or with Cy3 (#711-165-152, Dianova, Hamburg, Germany) for Homer1 identification. The secondary antibodies were diluted to 1 µg/ml in PBS containing 2.5% BSA, and were incubated for 3 hours at room temperature. The brain slices were finally subjected to five washes with PBS containing 2.5% BSA (each wash for 5 minutes), followed by two final 5-minute washes in PBS.

#### Immunostaining of SARS-CoV-2 particles

Intact SARS-CoV-2 samples deposited by the Center for Disease Control and Prevention were obtained through BEI resources, NIAID, NIH: isolate USA-WA1/2020, NR-52281 (Cat# NATSARS(COV2)-ERC, ZeptoMetrix, USA). The samples consisted of patient serum containing viral particles, fixed chemically using aldehydes, in a buffer containing BSA. An average of 9200 viral particles were allowed to adsorb onto single BSA-coated coverslips overnight at 4° C. Samples were mildly fixed with 4% PFA for 4 min before immunostaining using anti SARS-CoV-2 Spike Protein S1 (Cat# PA5-114447, ThermoFisher Scientific) and anti-human IgG (Fc)-Alexa 488 (Cat# 109-545-170, Jackson ImmunoResearch), as described above.

#### GFP-nanobody complex (TSR) generation

The monomeric (A206K) and non-fluorescent (Y66L) EGFP (mEGFP*) was modified to have an ALFA-tag on the N-Terminus and a HaloTag on its C-terminus (ALFA-EGFP-HaloTag). This construct was expressed in a NebExpress bacterial strain, and it had an N-terminal HisTag, followed by a bdSUMO domain, which enables the specific cleavage of the HisTag^39^ later on, after the purification procedures. Bacteria were grown at 37°C with shaking at 120 rpm in terrific broth (TB) supplemented with kanamycin. When reaching an optical density (OD) of ∼3, the temperature was reduced to 30°C and bacteria were induced using 0.4 mM isopropyl β-D-1-thiogalactopyranoside (IPTG), with shaking for another ∼16h. Bacteria lysates were incubated with Ni^+^ resin (Roche cOmplete) for 2h at 4°C. After several washing steps, the ALFA-tag-mEGFP(Y66L)-HaloTag protein was eluted by enzymatic cleavage on the column by using 0.1 µM of SENP1 protease for 15 minutes. Protein concentration was determined using Nanodrop (ThermoFisher), and purity was assessed by Coomassie gels. Complex formation was performed by mixing, for 1h at room temperature, in a final volume of 40 µl, the following: 25 pmol of ALFA-EGFP-HaloTag and 30 pmol of 3 different single-domain antibodies: FluoTag-Q anti-ALFA (Cat# N1505), FluoTag-X2 anti-GFP (clone 1H1, Cat# N0301) and FluoTag-X2 anti-GFP (clone 1B2), all from NanoTag Biotechnologies GmbH. The control experiments were performed by a similar procedure, without including the target protein ALFA-EGFP-HaloTag. Expression and purification of eGFP used in Supplementary Fig. 8 were performed as before^81^. Briefly, Neb Express *E. coli* strain (New England Biolabs) was cultured in terrific broth at 37°C and induced using 0.4 mM IPTG for 16h at 30°C. Bacteria pellets were sonicated on ice in 50 mM HEPES pH 8.0, 500 mM NaCl, 5 mM MgCl_2_, and 10% glycerol. After removing cell debris by centrifugation, the lysate was incubated for 1h with cOmplete His-Tag Purification Resin (Roche) at 4°C. After washing the resin in batch mode with more than 10 column volumes (CV), eGFP was enzymatically eluted using 0.1 µM of SUMO protease. Concentration was determined by absorbance at 280, using the molecular weight and extinction coefficient of eGFP. Purified protein was diluted in 50% glycerol and stored in small aliquots at -80°C.

#### Polyacrylamide gel electrophoresis (PAGE)

A primary mouse monoclonal antibody against synaptobrevin 2 (Cat# 104 211, Synaptic Systems) and a secondary antibody conjugated to Abberrior Star635P (Cat#ST635P-1002-500UG) were mixed with reducing 2x Laemmli buffer (63 mM Tris-HCl pH 6.8, 2% SDS, 100 mM DTT, 20% glycerol) and heated for 10 minutes at 96°C. The denatured and reduced samples were then loaded in a self-cast Tris-glycine 12% polyacrylamide gel, and 10µg of total protein was loaded per lane. Electrophoresis was run at low voltage, at room temperature. The gel was briefly rinsed using distilled water and fluorescence was read on a GE-Healthcare AI-600 imager using a far-red filter (Cy5 channel). Next, the gel was submerged for 4 hours in Coomassie Brilliant Blue solution to stain all proteins, following by incubation with destaining solutions, before finally being imaged using the same GE-Healthcare AI 600 gel documentation system.

#### Dot Blot

In a stripe of nitrocellulose membrane (GE Healthcare), 5 mg of bovine serum albumin (BSA) and 1 µg of ALFA-tagged EGFP(Y66L)-HaloTag were spotted and let to dry at room temperature. Membranes were then blocked in PBS supplemented with 5% skim milk and 0.05% Tween-20 for 1 h with tilting/shaking. FluoTag-X2 anti-GFP Cy3 (clone 1B1), FluoTag-X2 anti GFP-AberriorStar635P (clone 1H1) and Fluotag-X2 anti-ALFA AbberiorStar635P (all from NanoTag) were used at 2.5 nM final concentration in PBS with 5% milk and 0.05% Tween-20 for 1h with gentle rocking. After 1 h incubation at room temperature and protected from light, 5 washing steps using 2 ml each were performed with PBS supplemented with 0.05% Tween-20 for a total of 30 minutes. Membranes were finally imaged using a GE-Healthcare AI 600 system.

#### 1,6-hexanediol treatments

This compound (#240117-50G, Aldrich) was diluted in the neuronal Neurobasal-A culture medium at 3% for 2 minutes, and 10% for 12 minutes, before fixation and further processing for immunostaining.

#### Purified proteins

Immunoglobulins A and M were purchased from Jackson ImmunoResearch and Immunoglobulinss G from Abberior, Göttingen, Germany (AffinityPure IgA 109-005-011, ChromePure IgM 009-000-012, and ST635P-1001, respectively) and were diluted in PBS, before expansion procedures. Otoferlin was produced according to standard procedures^82^, and was diluted in 20 mM HEPES, 100 mM KCl, 0.05% DDM buffer, before being used at 0.4 mg/ml concentration. For GABA_A_ receptors a construct encoding the full-length human GABA_A_ receptor b3 subunit (Uniprot ID P28472), with an N-terminus TwinStrep tag, was cloned into the pHR-CMV-TetO2 vector^83^. A lentiviral cell pool was generated in HEK293S GnTI-TetR cells as described previously^84^. Cells were grown in FreeStyle 293 expression medium (#12338018, Gibco) supplemented with 1% fetal bovine serum (#11570506, Gibco), 1mM L-Glutamine (#25030149, Gibco), 1% NEEA (Gibco #11140050) and 5 mg/ml blasticidin (Invivogen #ant-bl-5b) at 37 °C, 130 r.p.m., 8% CO2 and induced as described^85^. Following collection by centrifugation (2,000 g, 15 min), the cell pellets were resuspended in PBS, pH=8 supplemented with 1% (v/v) mammalian protease inhibitor cocktail (Sigma-Aldrich). Cell membranes were solubilized with 1% (w/v) n-dodecyl β-D-maltopyranoside (#D3105GM, DDM, Anatrace) for 1h. The insoluble material was removed by centrifugation (12,500g, 15 min) and the supernatant was incubated with 300 mL Strep-Tactin® Superflow® resin (IBA lifesciences) while rotating slowly for 2h at 4°C. The beads were collected by centrifugation (300g, 5 min) and washed with 150mL of 0.04% (w/v) DDM, PBS pH=8. The sample was eluted in 2.5 mM Biotin, 0.02% (w/v) DDM, PBS pH=8 and used for imaging at 1 mg/ml concentration. For the purification of the GABA_A_ receptor in complex with the β3-β3 specific nanobody (Nb25)^86^, Nb25 was fluorescently labelled with STAR635P at the N-and C-termini generating Nb25-STAR635P. 20 μl of 10 μM Nb25-STAR635P was added to the sample prior to the elution step and incubated for 2h at 4°C while rotating. The excess Nb25-STAR635P was removed by washing the beads with 6 bed volumes of 0.04% (w/v) DDM, PBS pH=8, eluted with 2.5 mM Biotin, 0.02% (w/v) DDM, PBS pH=8 and used for imaging at 3mg/mL concentration. The same procedure was applied for the negative control anti-eGFP nanobodies. To test that Nb25-STAR635P could still bind the receptor, 2 μM of Nb25-STAR635P was added to the β3 homomeric receptor reconstituted in nanodisc as described previously^87^. 3.5 μl of the sample was applied to a freshly glow-discharged (PELCO easiGlow, 30 mA for 120s) 1.2/1.3 UltrAuFoil grid (Quantifoil), which were blotted for 2.5s and plunge-frozen using a Leica EM GP2 plunger at 14 °C and 99% humidity. Imaging was performed on a Titan Krios G2 microscope at the MRC LMB equipped with a F4 detector in electron counting mode at 300kV at a nominal magnification of 96,000 corresponding to a calibrated pixel size of 0.824 Å. 300 movies were collected using EPU (Thermo Fisher Scientific, version 2.0–2.11) with total dose of 38 e^-^/Å^2^ and 6.43s exposure time. The movies were motion corrected using MotionCor2^88^. Contrast transfer function estimation was performed with CTFFIND-4.1.13^89^. Particle picking was performed using a retrained BoxNet2D neural network in Warp^90^, followed by 2D classification in cryoSPARC^91^. Calmodulin was purified as previously described^92^, and was used in calcium free buffer: 150 mM KCl, 10 mM HEPES, 5 mM EGTA, or calcium+ buffer: 150 mM KCL, 10 mM HEPES, 2 mM CaCl_2_, at pH = 7.2, before expansion procedures. In brief, calmodulin 1 (mRNA reference sequence number NM_031969.2) was tagged with mEGFP and an ALFA-tag, for affinity purification purposes. The construct was transfected in HEK293 cells using Lipofectamine 2000 (Invitrogen, Carlsbad, CA, USA #11668019), following the manufacturer’s protocol. After expression for ∼24 hours, the cells were lysed in a PBS buffer containing 1% Triton X-100, 2 mM EDTA and a protease inhibitor cocktail. The debris was removed by centrifugation, and the supernatant was added to an ALFA Selector PE resin (NanoTag Biotechnologies), where it was allowed to bind for 60 minutes (4°C, under rotation). After two washes with lysis buffer and one wash with PBS (ice-cold), the bound proteins were eluted by adding the ALFA peptide. The purified protein was analyzed by Coomassie gel imaging (published in^92^).

#### Expansion procedures

X10 expansion of cultured cells was performed using proteinase K exactly as described in the following protocol article:^29^. X10 expansion relying on autoclaving (X10ht^47^) was performed as follows. The samples were incubated with 0.3 mg/ml Acryloyl-X (SE; #A-20770, Thermo Fisher Scientific) in PBS pH 7.4, overnight, at room temperature. The samples were then subjected to three PBS washes (5 minutes each), while preparing the gel monomer solution, exactly as described^29^. The solution was pipetted on parafilm and was covered by upside-down coverslips containing cells, or with brain slices that were then also covered with fresh coverslips. Polymerization was allowed to proceed overnight at room temperature, in a humidified chamber. Homogenization of proteins and single molecules were performed using 8 U/ml proteinase K (PK, #P4850 Sigma Aldrich now Merck) in digestion buffer (800 mM guanidine HCl, 2 mM CaCl_2_, 0.5 % Triton X-100, in 50 mM TRIS, #8382J008706, Merck), overnight at 50°C. Homogenization of cell cultures and brain slices was done by autoclaving for 60 minutes at 110°C in disruption buffer (5% Triton-X and 1% SDS in 100 mM TRIS, pH 8.0) followed by a 90 minutes incubation for temperature to cool down to safe levels. Before autoclaving, the gels were first washed using 1 M NaCl, and were then washed at least four times in disruption buffer, for a total time of at least 120 minutes. Gel expansion was then performed by ddH_2_O washing, for several hours, with at least five solution exchanges. Expansion was performed in 22 x 22 cm square culture dishes, carrying 400-500 ml ddH_2_O. When desired, the samples were labelled using 20-fold molar excess of NHS-ester fluorescein (#46409, ThermoFisher Scientific) in NaCHO_3_ buffer at pH = 8.3 for 1 h, before the washing procedure that induced the final expansion.

#### ZOOM expansion procedures

Fixed U2OS cultured cells were incubated in anchoring solution (25 mM Acrylic acid N-hydroxysuccinimide ester in 60% v/v DPBS and 40% v/v DMSO) for 60 minutes. Afterward, cells were moved to monomer solution (30% w/v Acrylamide and 0.014% w/v N-N’-methylenbisacrylamide in PBS buffer). After 60 min, the gelation process was started by adding initiators (0.5% w/v TEMED and 0.5% w/v APS) to the monomer solution. The hydrogel-cell hybrid was homogenized in detergent solution (200 mM SDS, 50 mM boric acid in DI water, pH titrated to 9.0), at 95 °C for 15 min, following by 24 h at 80 °C. ZOOM-processed samples were then stained using the previously mentioned anti α-tubulin antibodies (1:400 in PBST).

#### Microscope systems

For image acquisition, small gel fragments were cut and were placed in the imaging chamber presented in Supplementary Fig. 2. Paper tissues were used to remove any water droplets around the gels, before enabling the gels to equilibrate for at least 30 minutes on the microscope stage. Epifluorescence imaging was performed using an Olympus IX83 TIRF microscope equipped with an Andor iXon Ultra 888, 100× 1.49 NA TIRF objective, and an Olympus LAS-VC 4-channel laser illumination system. Confocal imaging was performed, for most experiments, using a TCS SP5 STED microscope (Leica Microsystems, Wetzlar, Germany), using a HCX Plan Apochromat STED objective, 100×, 1.4 NA, oil immersion. The LAS AF imaging software (Leica) was used to operate imaging experiments. Excitation lines were 633, 561, and 488 nm, and emission was tuned using an acousto-optical tunable filter. Detection was ensured by PMT and HyD detectors. Images were taken using a resonant scanner at 8 kHz frequency. 5D-stacks for zONE were performed using a 12 kHz resonant scanner mounted on a Leica TCSSP8 Lightning confocal microscope. Samples were excited with a 40% white light laser (WLL) at wavelengths of 633, 561 and 488 nm, and acquisitions were carried out using HyD detectors in unidirectional-xyct line scans or in uni-and bi-directional xyczt line scans.

#### Image acquisition

Objectives of 1.4, 1.45 and 1.51 NA were used to acquire images with a theoretical pixel size of 98 nm. For a higher resolution, the theoretical pixel size was set to 48 nm, at the cost of slightly lower detection rate. Images acquired on camera-based system had a predetermined pixel size of 100 nm. The acquisition speeds were ranging between 20 to 40 ms and 25 ms on a resonant scanner of 8 kHz and on a camera, respectively, for xyct. For hyperstacks of xyczt acquisitions, images were acquired using 8 kHz and 12 kHz scanners in bidirectional mode (after the necessary alignments) to compensate for speed loss. Images of 8 bit depth were acquired at a line format ranging from 128x128 to 256x256. The scanning modality on a confocal was set to “minimize time interval” (Leica LAS software). To maintain natural fluctuations of fluorophores, we did not use line accumulation or line averaging during scanning. A frame count starting from 200 and up to 4000 were acquired. We recommend a frame count of at least 1500 to 2000 for optimal computed resolution.

#### Image processing

ONE image processing is enabled through a Java-written ONE Platform under “ONE microscopy” in Fiji. The ONE microscopy plugin utilizes open-source codes from Bioformats Java library, NanoJ-Core, NanoJ-SRRF, NanoJ-eSRRF, and Image Stabilizer^6, 7, 93, 94^. ONE plugin supports multiple video formats of single or batch analyses in xyct. Hyperstacks with 5-dimensions xyczt format are processed with zONE module. This module allows the user to select the optical slices and channels to resolve at ultra-resolution. Upon irregularities in resolving one or more channels within one or more planes, zONE leaves a blank image, and computes the remaining planes within a stack. The image processing is fully automated and requires minimal initial user input. Aside from the expansion factor, preset values and analysis modalities are automatically provided (for more details, see Supplementary Fig. 1). The ONE plugin has a pre-installed safety protocol to skip failures in computations or uncompensated drifts, without affecting the progress of batch analysis. Data analysis, parameters and irregularities are reported in log files. The ONE plugin automatically linearizes the scale, based on radiality magnification and expansion factor corrections. In addition, ONE offers the possibility to correct for chromatic aberration by processing multi-channel bead images as a template that is applied to super-resolved images of the biological samples. The correction is performed by applying the Lucas-Kanade algorithm^93^. For the ONE Microscopy plugin to store complex multi-dimension images from hyperstacks, we modified the Java code of the ImageJ library and adapted it locally. ONE Platform source code and plugin are available on https://www.rizzoli-lab.de/ONE. For best performance, we recommend to download a preinstalled version on Fiji available via the same link.

#### Image analysis and statistics

For single-object analyses, such as synaptic vesicle or antibody analyses, signal intensities and distances between objects were analyzed manually using ImageJ (Wayne Rasband and contributors, National Institutes of Health, USA). Line scans were also performed and analyzed using ImageJ. For the analysis of PSDs (Fig. 2), spots were identified by thresholding band-pass filtered images, relying on empiric thresholds and band-pass filters, organized in the form of semi-automated routines in Matlab (version 2017b, The Mathworks, Inc, Natick, MA, USA). Spots were either overlaid, to determine their overall signal distributions, or their center positions were determined, to measure distances between spots (in either the same or different channels). The same procedure was used for the averaging analysis of CSF samples (Fig. 4) and for the analysis of spot distances for the GFP-nanobody assemblies (Extended Data Fig. 5). FWHM values were measured after performing line scans over small but distinguishable spots, as indicated in Fig. 1, followed by Gaussian fitting, using Matlab. The averaging analysis of GABA_A_ receptors is presented in detail in the main text, and was performed using Matlab. In brief, receptors were detected automatically, as particles with intensities above an empirically-derived threshold. To remove particles with uncompensated drift, we eliminated all receptors coming from images in which a large proportion of the particles were oriented similarly. We then inspected visually all of the remaining particles, to choose those that appeared to be in a “front view”, showing a reasonably round appearance, with nanobodies placed at the edges of the receptor (visible in the second color channel). All particles were centered on the intensity maxima of the respective GABA_A_R channel images. The particles were subjected to an analysis of the peaks of fluorescence, using a bandpass procedure, followed by identification of maxima^95^, the positions of the peaks were calculated to below-pixel precision and were rounded off to a pixel size of 0.384 nm (the starting pixel size was 1 nm). These positions were then mapped into one single matrix, which represents the “averaged receptor”, as indicated in the main text. Averaging analyses of tubulin and actin were performed similarly. In brief, microtubule segments, tubulin dimers or actin strands were selected manually and were overlaid, to generate average views. For the tubulin dimers, we calculated the peaks of fluorescence, as performed for the GABA_A_ receptors, above. Model objects were generated, as a comparison, by convoluting the amino acid positions in the respective PDB structures with empirically-derived ONE spots. All of these analyses were performed using Matlab. The signal to noise ratio (SNR) for single nanobodies was determined by measuring the average pixel intensities within the nanobody “spots” and away from them, and dividing the two measurements. Identically-sized circular regions-of-interest, sufficient to capture the nanobody spots completely, were used for both signal and background (noise) regions. Plots and statistics were generated using GraphPad Prism 9.3.1 (GraphPad Software Inc., La Jolla, CA, USA) or SigmaPlot 10 (Systat Software Inc., San Jose, CA, USA), or using Matlab. Statistics details are presented in the respective figures. Figures were prepared with CorelDraw 23.5 (Corel Corporation, Ottawa, Ontario, Canada).

## Ethics statement

Animals (Wistar rats, P0 to P1) were treated according to the regulations of the local authority, the Lower Saxony State Office for Consumer Protection and Food Safety (Niedersächsisches Landesamt für Verbraucherschutz und Lebensmittelsicherheit), under the license Tötungsversuch T09/08. For human patients, the informed consent of all of the participants was obtained at the Paracelsus Elena Klinik, following the principles of the Declaration of Helsinki.

## Acknowledgements

We thank Christina Zeising, Gabriele Klaehn, Anita-Karina Jaehnke, and Jannik Hentze (University Medical Center Göttingen) and Ulf Schwarz (Leica Microsystems) for excellent technical assistance. We thank Ali Kassem (Beirut, Lebanon) for driver debugging and integration. We acknowledge the University of Massachusetts Medical School Sanderson Center for Optical Experimentation Core Facility (RRID: SCR_022721), Massachusetts Life Sciences Center, and Christina Baer for assistance with imaging. We thank Nils Brose (Max Planck Institute for Multidisciplinary Sciences, Göttingen, Germany) for help with access to his fluorescence microscopy facility. We thank Markus Krone (Max Planck Institute for Multidisciplinary Sciences, Göttingen, Germany) and Frank Kötting (European Neuroscience Institute) for their support in building the gel stabilization chambers. We thank Eugenio Fornasiero and Christian Tetzlaff (University Medical Center Göttingen, Germany) and G.B. Rizzoli for comments on the initial manuscript. The work was supported by grants from the German Ministry for Education and Research, 13N15328/NG-FLIM and from the German Research Foundation (Deutsche Forschungsgemeinschaft, DFG), SFB1286/A03/B02/Z04, SFB1190/P09 to S.O.R., the European Research Council (ERC) under the European Union’s Horizon 2020 research and innovation programme (grant agreements No 835102 and 964016), and the DFG under Germany’s Excellence Strategy - EXC 2067/1-390729940. Support from the DFG grant SFB894/A9 (to UB) and the UK Medical Research Council grants MR/L009609/1 and MC_UP_1201/15 (to A.R.A.) is also acknowledged.

## Data availability statement

Image data are available from the corresponding authors on reasonable request.

## Code availability statement

The ONE platform plugin software (source code) will be available soon.

## Author contributions

AHS and SOR conceived the project. AAC and UB developed the ONE Platform plugin. AHS and SOR designed and performed the experiments. UB supervised the ONE experiments at CIPMM. ESB supervised the ONE experiments at MIT. SK supervised the ONE experiments at institute for X-Ray Physics. MS supervised the ONE experiments at Würzburg University. AHS, VI and SOR analyzed the data. RC, NM and PF contributed to the tubulin experiments. CZ performed the ONE experiments at the MIT. JK contributed to the bassoon in tissues experiment. SVG contributed to the 1,6-hexanediol experiments. TCM generated the *in vitro* microtubules. NA and UB performed the ONE experiments at the CIPMM. MM and JP purified otoferlin protein. HC and TM generated the otoferlin AlphaFold model. DM and ARA generated the GABA_A_ receptors and the Cryo-EM data. SR purified calmodulin protein. JE performed the ONE experiments at Würzburg University. RC and LA contributed to the experiments at Leica GMBH, Mannheim. FO generated the TSRs. DC generated the tissue sections. KAS assisted with the initial implementation of X10ht experiments. CT and BM provided the patient CSF specimens. TFO verified the analysis of the PD data. MS contributed to the understanding of photophysics fluctuations and ESB contributed to the understanding of ExM gel behavior. SOR wrote the manuscript, which was revised by all other authors, with especially strong contributions from AHS, ARA, MS and ESB.

## Competing interests

S.O.R. and F.O. are shareholders of NanoTag Biotechnologies GmbH. The remaining authors declare no competing interests. E.S.B. is an inventor on multiple patents related to expansion microscopy, and co-founder of a company working on commercial applications of expansion microscopy.

## Notes

### Summary of Updates

Shaib et al., 2022, has been updated to include many types of new data, all presented in new figures: -A demonstration of ONE feasibility in many laboratories in Germany and USA (Supp. Figs. 29 to 33) -Averaging analysis for single GABAa Receptors, to derive their protein shape in vitro, shown in Fig. 3, and supported by extensive new data (Supp. Figs. 12 and 13) -An averaging analysis of tubulin in vitro, to also derive its molecular shape (Supp. Fig. 15) -Averaging analysis for actin filaments in cells, again for the purpose of deriving their molecular shape (Fig. 4) -Analyses of ONE microscopy for post-expansion immunolabeling of PSD95 (Supp. Fig. 21), microtubules (Supp. Fig. 32) and bassoon (Supp. Fig. 33) -Analyses of ONE images in the sub-nanometer range (Supp. Fig. 24) -Quantifications of the SRRF precision, using DNA Origami nanorulers (Supp. Fig. 2) -An in-detail explanation of why ONE microscopy is able to provide molecular resolution (Supp. Fig. 3 and 4) -New analyses of single proteins, as eGFP (Supp. Fig. 8) -New analyses of synapses (Supp. Fig. 20) -Extensive FRC analyses of many samples (Supp. Figs. 23 and 28) -Technical updates, including structure analyses (Supp. Fig. 25), examples of bleaching curves (Supp. Fig. 7), drift correction (Supp. Fig. 14), analysis explanations and quantifications (Supp. Figs. 17 and 18), comparisons with current technologies (Supp. Fig. 22) Other updated elements: authors and author affiliations, main text, main figures, Materials and Methods, Extended Data Figures and Supplementary Figures.

